# SpatialScope: A unified approach for integrating spatial and single-cell transcriptomics data using deep generative models

**DOI:** 10.1101/2023.03.14.532529

**Authors:** Xiaomeng Wan, Jiashun Xiao, Sindy Sing Ting Tam, Mingxuan Cai, Ryohichi Sugimura, Yang Wang, Xiang Wan, Zhixiang Lin, Angela Ruohao Wu, Can Yang

## Abstract

The rapid emergence of spatial transcriptomics (ST) technologies are revolutionizing our under-standing of tissue spatial architecture and their biology. Current ST technologies based on either next generation sequencing (seq-based approaches) or fluorescence in situ hybridization (image-based approaches), while providing hugely informative insights, remain unable to provide spatial characterization at transcriptome-wide single-cell resolution, limiting their usage in resolving detailed tissue structure and detecting cellular communications. To overcome these limitations, we developed SpatialScope, a unified approach to integrating scRNA-seq reference data and ST data that leverages deep generative models. With innovation in model and algorithm designs, SpatialScope not only enhances seq-based ST data to achieve single-cell resolution, but also accurately infers transcriptome-wide expression levels for image-based ST data. We demonstrate the utility of SpatialScope through comprehensive simulation studies and then apply it to real data from both seq-based and image-based ST approaches. SpatialScope provides a spatial characterization of tissue structures at transcriptome-wide single-cell resolution, greatly facilitating the downstream analysis of ST data, such as detection of cellular communication by identifying ligand-receptor interactions from seq-based ST data, localization of cellular subtypes, and detection of spatially differently expressed genes.

## Introduction

Single-cell RNA sequencing (scRNA-seq) characterizes the whole transcriptome of individual cells within a given organ, providing remarkable opportunities for broad and deep biological investigations of diverse cellular behaviors [1, 2, 3]. However, scRNA-seq does not capture the spatial distribution of cells due to samples having to undergo tissue dissociation [4]. As spatial information is so critical to understanding communication between cells, many related scientific questions related to cellular communication cannot be fully addressed by scRNA-seq alone [5].

Current ST approaches are predominantly based on either next-generation sequencing (seq-based) or fluorescence in situ hybridization (image-based). Seq-based approaches, such as 10x Visium [6] and Slide-seq [7], can detect transcriptome-wide gene expression within spatial spots, but each spot often contains multiple cells. Therefore, the resolution of present seq-based approaches do not achieve single-cell resolution, which limits their usage in resolving detailed tissue structure and in characterizing cellular communications (e.g., identifying ligand-receptor interactions [8]). Image-based approaches such as seqFISH [9] and MERFISH [10] achieve single-cell resolution, but are limited to profiling panels of tens to hundreds of genes per sample, leaving the majority of the transcriptome unmeasured. Users of these image-based methods need to have well-defined biological hypotheses to design an appropriate and useful gene panel, and it is unlikely to generate incidental discoveries in this scenario.

Ideally, the integration of single-cell and ST data should allow us to characterize the spatial distribution of the whole transcriptome at single-cell resolution, by combining their complementary information. However, existing integration methods are far from satisfactory in real data analysis [11]. There are now several cell-type deconvolution methods for ST data, including RCTD [12], Cell2location [13], CARD [14] and spatialDWLS [15]. When these deconvolution methods are applied to seq-based ST data, they only estimate the proportions of cell types in each spatial spot but cannot achieve single-cell resolution. Therefore, the aforementioned limitations of not having single-cell resolution remain unresolved. For imagebased ST data, methods developed to infer unmeasured gene expressions, such as Tangram [16], gimVI [17] and SpaGE [18] are not sufficiently accurate, especially when ST expression data are sparse [11]. Therefore, there remains a need for accurate statistical and computational methods for integrating single-cell and ST datasets [4].

Herein we introduce SpatialScope, a unified approach to integrating scRNA-seq reference data and ST data generated from various experimental platforms, applicable to both seq-based ST data (e.g., 10x Visium and Slide-seq) and image-based data (e.g., MERFISH). By leveraging deep generative models, SpatialScope can resolve the spot-level data composed of multiple cells to single-cell resolution when it is applied to seq-based ST data. There are two key features of SpatialScope. First, it can greatly improve cell type identification by exploiting spatial information of cells through Potts model and properly correcting for batch effect between ST and scRNA-seq reference data. Second, unlike alignment-based methods such as Tangram [16] and CytoSPACE [19] that assign existing cells from scRNA-seq data to spatial spots, SpatialScope can generate the gene expressions of pseudo-cells using the learned deep generative model to match the observed spot-level gene expression in space. Consequently, SpatialScope can decompose the observed gene expression at each spot into the single-cell level gene expression accurately. In addition, for image-based ST data, SpatialScope can learn the distribution of gene expressions from the scRNA-seq data and then infer transcriptome-wide expression of the unmeasured genes in the sample, conditioned on the observed tens to hundreds of genes in that sample. With the above features, SpatialScope allows more in-depth and informative downstream analyses at single-cell resolution. Using ST data generated from various experimental platforms, such as 10x Visium, Slide-seq and MERFISH data, we show that the results of SpatialScope enable spatially resolved cellular communications mediated by ligand-receptor interactions and spatially differentially expressed genes expression, highlighting SpatialScope’s utility in elucidating underlying biological processes. By applying SpatialScope to human heart data, ligand and receptor pairs that are essential in vascular proliferation and differentiation are detected using higher resolution ST data generated by SpatialScope. Some meaningful genes absent in MERFISH data are detected as DE genes through the imputation of SpatialScope. Very recently, SpatialScope has been applied to enhance the resolution of ST data generated from human embryonic hematopoietic organoids, producing single-cell resolution ST data which was then used to detect spatially resolved cell-cell interactions and co-localization of different cell types [20]. This single-cell resolution decomposition of the original data has allowed us to identify additional biological findings that were not possible at spot-level.

## Results

### Overview of the SpatialScope method

By leveraging the deep generative model, SpatialScope enables the characterization of spatial patterns of the whole transcriptome at single-cell resolution for ST data generated from various experimental platforms. To present our key idea better, we begin our formulation with gene expression decomposition of seq-based ST data from the spot level to the single-cell level. Let 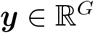 be the expression levels of *G* genes (after batch effect corrections) at a spot in seq-based data. Suppose that we already know that the spot-level gene expression ***y*** comes from two cells of different types, ***y*** = ***x***_1_ + ***x***_2_ + ***ϵ***, where ***x***_1_ and ***x***_2_ are the true gene expression levels of cells 1 and 2 whose cell types are denoted as *k*_1_ and *k*_2_, respectively, and the independent random noise ***ε*** is assumed to be 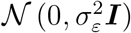 for convenience. We aim to decompose ***y*** into ***x***_1_ and ***x***_2_, and thus obtain the single-cell resolution gene expression at the given spot. To achieve this, we use a deep generative model [21, 22, 23] to learn the expression distributions of cell types *k*_1_ and *k*_2_ from the scRNA-seq reference data, denoted as *p* (***x***_1_ | *k*_1_) and *p* (***x***_2_ | *k*_2_). Based on Langevin dynamics [24, 22], we can obtain the decomposition by sampling ***X*** = [***x***_1_; ***x***_2_] from the posterior distribution *p* (***X*** | ***y***, *k*_1_, *k*_2_),

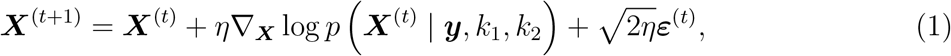

Where 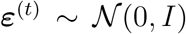 and *η* > 0 is the step size, *t* = 1,…, ∞. By Bayes rule, we have 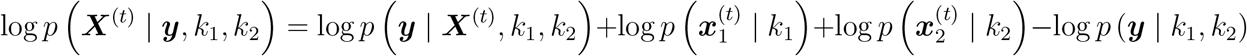

Noting that ∇_*X*_ log *p* (***y*** | *k*_1_, *k*_2_) = 0, this makes it easy to obtain posterior samples from the Langevin dynamics as

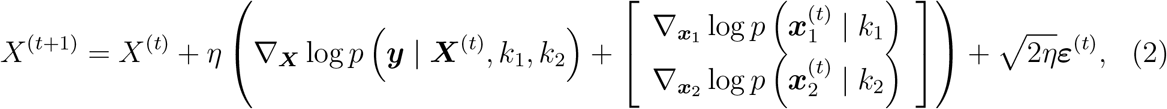

where 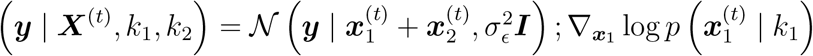 and 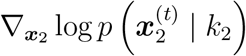 are known as the score function which can be learned from the scRNA-seq reference data. The samples from the posterior distribution *p* (***X***^(*t*)^ | ***y***, *k*_1_, *k*_2_) recover gene expression levels of the two cells, achieving single-cell resolution.

To implement the key idea formulated above, SpatialScope comprises three steps of real data analysis (Fig. 1): (i) Nucleus segmentation; (ii) cell type identification; and (iii) gene expression decomposition with a score-based generative model. Specifically, we first perform nuclei segmentation on the hematoxylin and eosin (H&E)-stained histological image to count the number of cells at each spot. Second, for cell type identification (i.e., *k*_1_, and *k*_2_) at each spot, we develop a fast and accurate method by integrating scRNA-seq and ST data. Third, we learn the conditional score generative model (i.e., 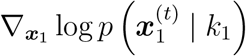 and 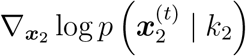) in a coherent neural network to approximate the expression distribution of different cell types from scRNA-seq data, and then use the learned model to decompose gene expression from the spot level to the single-cell level, as outlined above. Based on the same modeling principle, we generalize SpatialScope to infer the unmeasured gene expression for image-based ST data, conditional on the observed gene expression levels. We introduce the details of SpatialScope in the method section.

**Figure 1:**
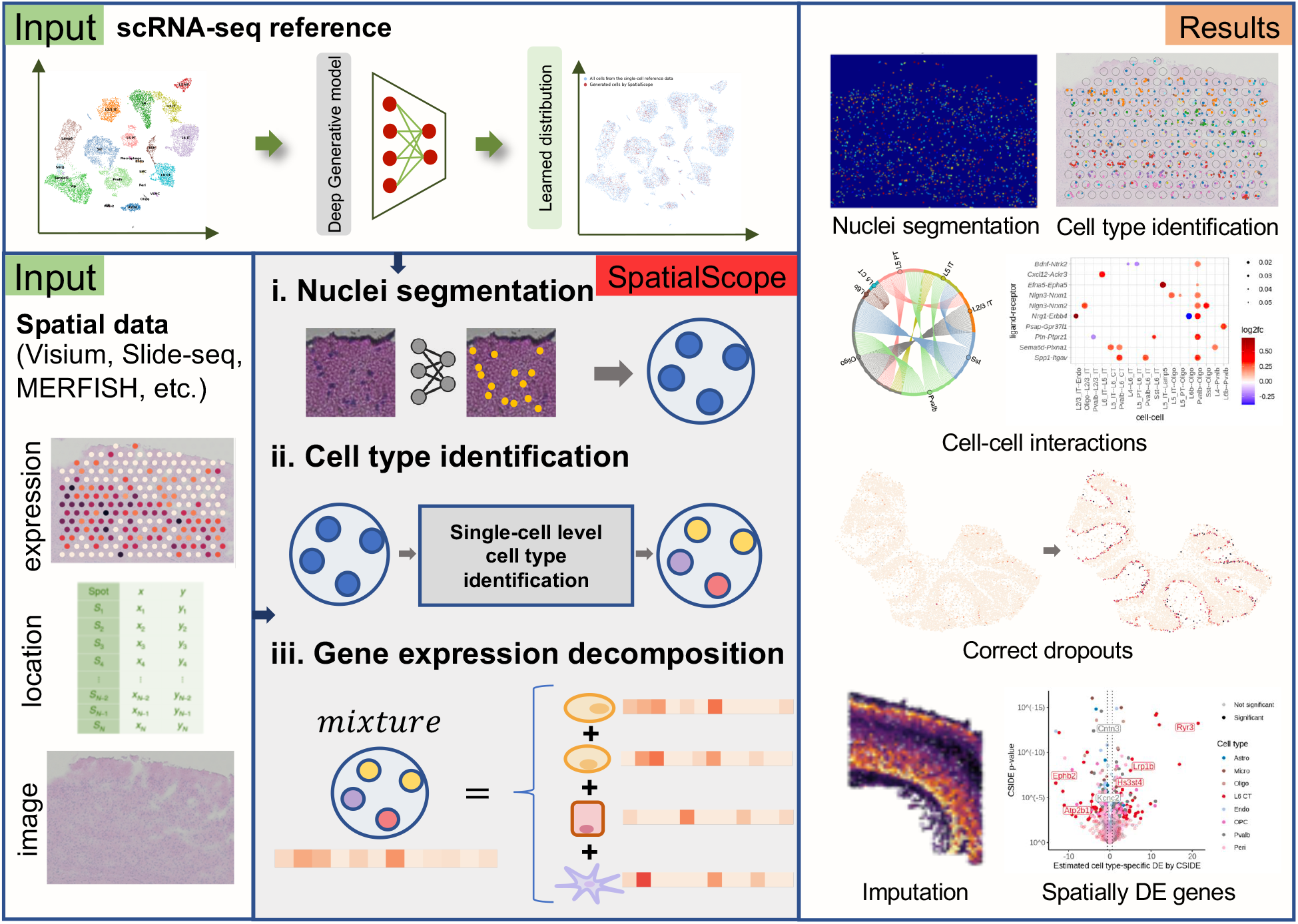
Overview of SpatialScope. SpatialScope is designed to infer the spatially resolved single-cell transcriptomes based on the distribution learned from scRNA-seq reference data. SpatialScope first segments single cells in low-resolution ST data such as Visium, then identifies the cell type label for each segmented cell within the spot. Finally, conditioning on the infer cell type labels, SpatialScope decomposes the spot-level gene expression profile into single-cell level, allowing more in-depth and informative downstream analyses at single-cell resolution.

### SpatialScope accurately identifies cell types and enables the integration of multiple slices

Without single-cell resolution in ST datasets, fine-grained cell gradients cannot be visualized, and understanding cellular communication within tissues is not possible. Ideally, tools for ST data analysis should leverage single-cell expression profiles either generated from paired samples or from the growing body of published tissue atlases to infer single-cell types and their comprehensive gene expression profiles, which would in turn enable interpretation of cell-cell interactions, localization, and spatial trajectories. To get reliable inferences of gene expression profiles at single-cell resolution, accurately identifying single-cell types is fundamental. Each spot in the ST data may contain multiple single cells whose signals are mixed at the spot level and we can only observe the low-depth mixed signal for each spot. Successful cell type identification means assigning cell type labels to the individual cells within each spot. SpatialScope enables accurate cell type identification by integrating annotated scRNA-seq data and ST data. Further, SpatialScope takes advantages of the inclusion of spatial information, enabling reliable single-cell type identification. Although a few alignment-based methods, such as Tangram and CytoSPACE, also can achieve single-cell type identification, the spatial information is missing. SpatialScope utilizes spatial information by factoring in neighbor cells’ identities when assigning each cell’s cell-type, which is achieved by introducing a spatial neighbor structure into the prior (Fig. 2a, Supplementary Method). To demonstrate the strengths of SpatialScope in this aspect, we compared its ability to identify cell types on spot-level ST data against several other computational tools for ST data. We compared SpatialScope with Tangram and CytoSPACE, which can also perform cell type identification at single-cell resolution. Since ground truth cell type of individual cells are unknown in real low-resolution ST data, we used a single-cell resolution MERFISH dataset [25] to generate a simulated dataset: The MERFISH data is partitioned into two regions, with one region serving as the single-cell reference data, and the second region serving to mimic low-resolution ST data by aggregating the cells on uniform grids to simulate spots (Fig. S1). Since the cell type of each cell in MERFISH dataset is annotated, we used the annotated cell type labels for individual cells within the simulated spots as ground truth.

**Figure 2:**
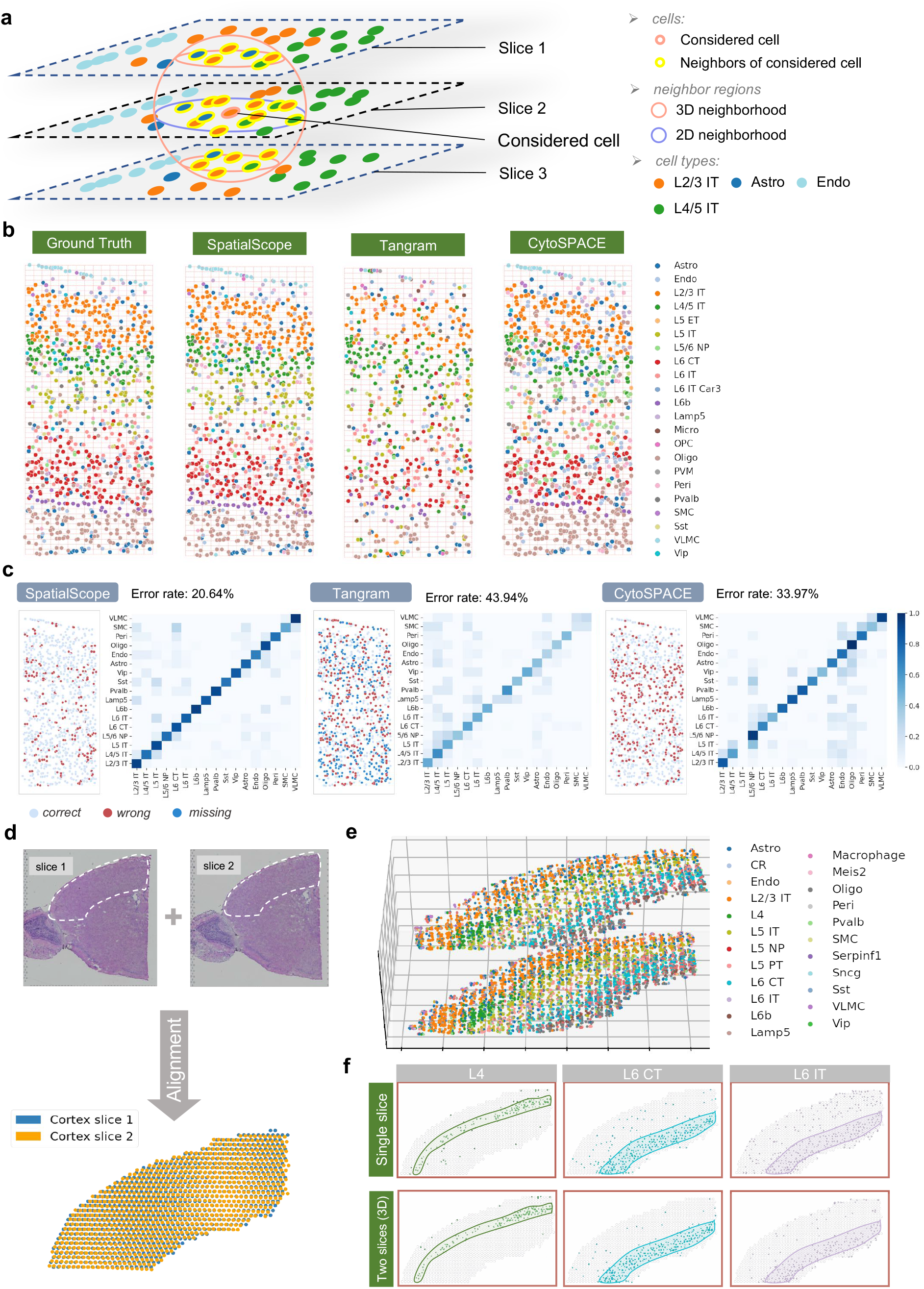
Simulated study of cell type identification and SpatialScope applying on a real multiple slices mouse brain cortex data. **a**, SpatialScope utilizes spatial information by encouraging neighboring cells to belong to the same cell type. Blue big circle shows the neighborhood when considering a single slice. Red big circle shows the neighborhood when considering multiple slices, resulting in integrating information from multiple slices. **b**, A spatial scatter plot displays identified single cell types on each cell location from ground truth and different methods. Each grid represents a simulated spot containing multiple cells. **c**, Scatter plot of single cells in simulated spatial data (left). Different colors show cells with correct/wrong cell type identification and cells missed by each method. Confusion matrix of true versus identified single cell types by different methods. Color in the confusion matrix represents the proportion of the cell type on the y-axis identified as the cell type on the x-axis. **d**, H&E staining histology of two adjacent slices of mouse brain cortex from 10x Visium (top). White polygons indicate the region of interest. Alignment results using PASTE [36] of the two slices (bottom). **e**, The SpatialScope identified cell type labels for the stacked 3D ST data constructed by two slices. **f**, Comparison cell type identification results of using single slice (top) and multiple slices (bottom). The first column to third column show spatial cell locations that identified as L4, L6 CT and L6 IT by SpatialScope, respectively. Circled area indicates regions of different layers: L4, L6 CT and L6 IT.

We evaluated the performance of cell type identification within simulated spots of different grid sizes (representing spatial data with different resolutions) and total UMI counts per spot (Fig. 2b, Fig. S2). The number of cells in the spots ranges from 1 to 6 for grid size 42 × 42 μm and from 1 to 3 for grid size 16 × 16*μ*m. The total UMI counts per spot range from 130 to 520, which respectively represent averaged half-cell UMIs and averaged two-cell UMIs in the original MERFISH data. The cell type identification results are similar across settings, with SpatialScope identifying clear layer structure at single-cell resolution at all settings. The results under the setting, grid size = 34 × 30(μm); total UMI counts per spot = 260, are visualized in Fig. 2b-c. Fig. 2b shows the ground-truth cell type labels and the cell type identification results of SpatialScope, Tangram and CytoSPACE. SpatialScope produced closer results to ground truth compared to Tangram and CytoSPACE.

By contrast, Tangram missed many cells because it does not use the actual number of nuclei in the histological image to determine the output cell number. CytoSPACE inaccurately assigned cell type labels in several layers, including the L5 IT layer (Fig. 2b). This is also reflected by the confusion matrix, with Tangram having a lower on-diagonal correlation and CytoSPACE showing noise and inappropriate off-diagonal correlation (Fig. 2c). The overall error rates of cell type identification for SpatialScope, Tangram and CytoSPACE are 20.64%, 43.94% and 33.97%, respectively (Fig. 2f), indicating that SpatialScope substantially outperforms Tangram and CytoSPACE in terms of cell type identification. We also performed cell type identification on other three simulated datasets generated by various single-cell resolution ST data (e.g., STARmap [26], MERFISH data from different tissues, etc.) and SpatialScope also compares favorably to Tangram and CytoSPACE on all datasets with respect to cell type identification error rate (Fig. S3b-d). To compare SpatialScope with other deconvolution methods, we calculated Pearson correlation (PCC) and root-mean-square error (RMSE) between true and estimated cell type proportion of spots and found that SpatialScope is the top-performing (Fig. S3a) and computationally efficient (Fig. S3e) method in all three simulated datasets. Furthermore, when there are missing cell types in the reference dataset, we showed that SpatialScope achieved the highest robustness over the compared methods by predicting the cells as the most transcriptionally similar cell type in the reference (supplementary note section 2.7.2)

Recently, ST data with multiple parallel slices in a tissue from one or more samples is become more widely available and is being generated at an accelerated pace. SpatialScope is also applicable to the data with multiple slices and its performance is further enhanced by integrating multiple slices. SpatialScope leverages 3D spatial information by adding a prior similar to 2D spatial neighbor structure. In the 3D case, the neighbors of a cell also include those across slices in the z-direction (Fig. 2a), and taking this into account improves the accuracy of cell type identification. To illustrate how SpatialScope is applied to multi-slices data, we considered a real two-slice dataset of spot-level mouse brain cortex data (Fig. 2d). Both of the two slices are from 10x Visium and they are adjacent slices from the mouse brain cortex, and these two-slices data serve as the spatial data. Separately, we used a published mouse brain scRNA-seq data (Smart-seq2) [27] as the single-cell reference, which is comprised of 14,249 cells across 23 cell types (Fig. S4). We first segmented single cells independently in the two corresponding H&E-stained histological images from the same tissue sections and located 3,777 and 3,034 cells within 812 and 794 spots from the brain cortex of slice 1 and slice 2, respectively. Using PASTE [28] to align multiple adjacent tissue slices, we then successfully constructed a 3D aligned ST data for the mouse brain cortex tissue (Fig. 2d). We applied SpatialScope to the 3D aligned ST data for cell type identification. We evaluated the accuracy of the inferred cell type labels based on the known spatial organization of cell types in the brain cortex: The mouse brain cortex consists of four main layers of glutamatergic neurons (L2/3, L4, L5 and L6), and cell type labels identified by SpatialScope accurately reconstructed these multi-layer structures in both slices of mouse brain cortex (Fig. 2e and Fig. S5). Alignment-based and deconvolution methods can only handle one slice at a time, and as a result the tissue layer structure can be misidentified (see Fig. S6 for more details). By incorporating 3D spatial structure and borrowing information from adjacent slices, SpatialScope reduces cell mis-identification compared to when only a single slice is used by taking into account neighboring cell types (Fig. 2f). For example, some mis-identified L6 CT and L6 IT cells in the upper layer of brain cortex are corrected when using two slices (Fig. 2f).

### SpatialScope generate single-cell resolution gene expression profiles and enable interpretation of cell-cell interactions

The goal for gene expression recovery is to decompose mixed reads within each spot and generate single-cell resolution gene expression profiles, which overcomes the limitation of low-resolution ST data. For SpatialScope, this step follows the step of cell type identification. In the gene expression decomposition step, SpatialScope first learns a conditional score-based generative model, which is conditioned on cell type, to approximate the distribution of gene expressions from a given scRNA-seq reference data (Fig. S7). Then based on the learned generative model and cell-type label of each single cell, SpatialScope decomposes the spot level gene expressions into single-cell level (see Methods). Accurate cell type identification is necessary for reliable inference of single-cell comprehensive gene expression profiles for SpatialScope as well as other alignment-based methods. For example, Tangram and CytoSPACE complete cell type identification and gene expression decomposition at the same time, and in this process, alignment of cells with the wrong cell type naturally leads to unreliable single-cell gene expression inference. As such, the results shown in the previous section and those of single-cell resolution gene expression decomposition are closely related.

To evaluate the performance of SpatialScope in gene expression decomposition, we used the same simulated data (Fig. S1) and compared SpatialScope with Tangram and CytoSPACE. The gene expression profiles for individual cells within the simulated spots were used as ground truth. We compared the gene expressions inferred by the three methods with the ground truth gene expression. All the results of SpatialScope are produced based on the cell type identification results in the last section.

Fig. 3a shows an example of how SpatialScope decompose spot-level gene expression profiles into cell-level gene expression profiles, where a spot containing two cells is decomposed to two single cell gene expression profiles conditioned on the fact that SpatialScope has already identified those two cells as endothelial and oligodendrocyte respectively. The decomposed gene expression of the two individual cells by SpatialScope (red dots) are clustered closely with the ground truth expression levels (green dots), with mean cosine similarity as high as 0.90. In contrast, inferred cells generated using Tangram (purple dots) and CytoSPACE (orange dots), which both use alignment-based methods, are more dissimilar to ground truth with mean cosine similarities of 0.44 and 0.57, respectively (Fig. 3a). By accurate gene expression decomposition, SpatialScope recovers the higher spatial resolution afforded by the original MERFISH data that is lost in the simulated ST data (Fig. 3b and Fig. S8a). As some cells with weak signals may be missed in the nucleus segmentation step, we also evaluated the performance of SpatialScope and the compared method when the ground truth cell number is inconsistent with the estimated cell number in the spots. In brief, we observed that SpatialScope is robust over inconsistent cell numbers and capable of identifying the remaining ground truth cells with highly matched transcriptional profiles (supplementary note section 2.7.3).

**Figure 3:**
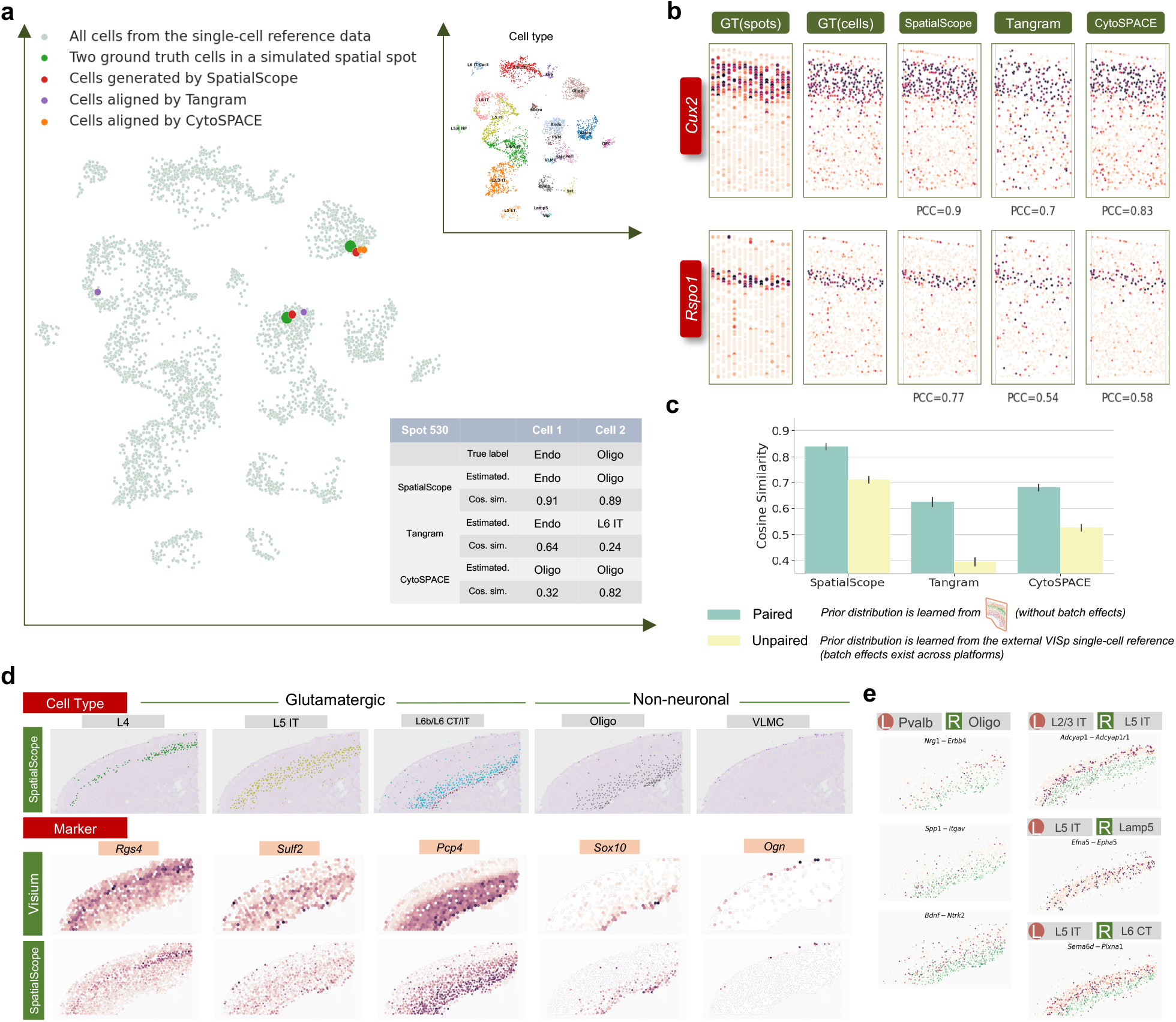
SpatialScope generates single-cell resolution gene expression profiles and enable interpretation of cell-cell interactions. **a**, The UMAP plot of gene expression decomposition example for a simulated spot with two cells from endothelial and oligodendrocyte, respectively. Dots of different colors represent gene expression profiles of two single cells decomposed from this simulated spot by different methods. The table shows the cosine similarity between the ground truth and decomposed gene expressions of two single cells in the simulated spot. In the table, “True label” represents ground truth cell type label; “Estimated.” represents estimated cell type label by different methods; “Cos. sim.” represents cosine similarity between ground truth gene expression profiles and estimated gene expression profiles by different methods. **b**, The spatial expression pattern of two layer’s marker genes before and after applying SpatialScope’s gene expression decomposition are displayed. SpatialScope decomposes spots to single-cell resolution, resulting in a refined spatial map of gene expression. **c**, The cosine similarities between the ground truth and decomposed single-cell level gene expression profiles of different methods are displayed. Two different single-cell reference data are used to evaluate the robustness of gene expression decomposition. **d**, Top, spatial cell locations identified as L4, L5 IT, L6b/L6 CT/L6 IT, Oligo and VLMC by SpatialScope using multiple slices. Middle, spot-level expression levels of the corresponding cell-type-specific marker genes in original Visium data. Bottom, refined single-cell resolution expression levels of the corresponding marker genes by SpatialScope **e**, Visualization of some representative molecular interactions detected in the 3D aligned single-cell resolution spatially resolved transcriptomic data produced by SpatialScope. The scatter plot shows the expression level of ligand-receptor pairs in corresponding cell type pairs. The expression of ligands and receptors are colored by orange and green, respectively.

In real use cases, it may be impractical to generate paired scRNA-seq data for each ST profiling experiment, either due to budget or sample availability. Thus, we also tested the accuracy and robustness of different tools in generating single-cell gene expression decomposition from ST-data based on either paired scRNA-seq data or an independently generated scRNA-seq reference of the same tissue type. We calculated cosine similarities between the inferred single-cell level gene expression and ground truth for all cells under different scenarios as a measure of accuracy (Fig. 3c). As shown in Fig. 3c, SpatialScope achieved significantly higher accuracy irrespective of paired/unpaired single-cell reference; the performances of alignment-based methods, Tangram and CytoSPACE, decreased dramatically when the reference data is generated from the same tissue type but different biological samples (unpaired setting in Fig. 3c). Furthermore, when varying the UMI subsample rate in the construction of simulated spots, we found that SpatialScope’s performance remains robust even when the simulated ST data is very sparse (Fig. S8b). This is significant because the per cell transcript capture rate of ST methods is known to be much lower than scRNA-seq, and ST data is generally sparse in nature, which poses challenges in data analysis [29, 30, 31]; accurate decomposition that is robust to sparsity alleviates this problem. The overall superior performance of SpatialScope is likely due to leveraging the learned distribution to generate pseudo/unobserved cells that fit the spot-level expression data, while alignment-based methods can only align existing cells from scRNA-seq reference data to spatial data. By fitting the spot-level expression, SpatialScope exhibits robustness to different reference data and adaptively improves when there is more information (e.g. more UMIs).

To further investigate SpatialScope’s gene expression decomposition performance in real data applications, we applied SpatialScope to analyze 10x Visium data of mouse brain cortex that have two adjacent slices, which is also used in cell type identification analysis. Further, we showed that gene expression decomposition enables more in-depth and informative cell-cell interactions analyses at single-cell resolution. We also found that multiple slices increased the power of cell-cell interactions detection by integrating information across multiple slices.

The same that the gene expression decomposition was performed after cell type identification described in last section. Through decomposition of gene expressions from the spot-level into the single-cell level, we refined the spatial transcriptomic landscape of mouse brain cortex at singlecell resolution while maintaining accurate spatial patterns of gene expressions (Fig. 3d). By contrast, Tangram and CytoSPACE cannot reconstruct the expected spatial expression patterns for a few marker genes at single-cell resolution (Fig. S6d). Next, we show that spatially resolved transcriptomic data at single-cell resolution produced by SpatialScope, with the additional help of 3D alignment, enabled us to infer reliable spatially proximal cell-cell communications. Compared to very few ligand-receptor signaling detected in a single slice only (Fig. S9d-e), we detected widespread proximity interactions between Parvalbumin-positive neurons (Pvalb) and Oligodendrocytes (Oligo) when analyzing the 3D aligned ST data (Fig. 3e and Fig. S9a). The identified ligand and receptor pairs show strong enrichment in multiple biological processes/pathways that are essential in the regulation of neuronal development in the cortex, including synapse organization and assembly, oligodendrocyte differentiation, and regulation of gliogenesis (Fig. S9b) [32]. For example, the cell-cell communication mediated by *Nrg1* and *Erbb4* interaction is well documented, and Neuregulin ligands play a role in proliferation, survival and maturation of oligodendrocytes through *Erbb4* pathways [33]. *Spp1–Itgav* was also suggested to mediate communication and migration between oligodendrocytes and microglia [34]; our analysis shows that this molecular interaction might also occur between Pvalb neurons and oligodendrocytes, providing a possible direction for further investigation. We also detected extensive cellular communications between subtypes of neurons, such as *Adcyap1-Adcyap1r1* between L2/3 IT and L5 IT, *Efna5-Epha5* between L5 IT and Lamp5, and *Sema6d-Plxna1* between L5 IT and L6 CT (Fig. 3e). These molecular interactions have been reported to be vital for cortical development in brain [35, 36]. The interacting cell types detected by SpatialScope provide more comprehensive understanding of cellular/molecular interactions in the cortex.

### SpatialScope enables high resolution identification of cell types and candidate pathways for cellular communication in human heart tissue

The human heart is a highly functionally coordinated organ, and different cell types within the same tissue must act in concert with precise feedback and control. Previous single-cell profiling of the human heart identified cellular subtypes with high levels of specialization in their gene expression corresponding to their roles in regeneration/renewal or as fully differentiated cells that participate in blood circulation and pacing [37]. With spatial transcriptomics, there is additional opportunity to understand these cellular specializations in the context of the complex architecture of the human heart. We applied SpatialScope to a real ST dataset of adult heart tissue profiled at spot-level [38], and show that decomposed single-cell transcriptomes enables localization of cellular subtypes at high resolution. Furthermore, assessment of ligand-receptor co-expression in neighboring cells reveals candidate pathways that facilitate cellular communication in a given tissue region.

First, we segmented single cells in the corresponding H&E stained image and located 10,734 cells within 3,813 spots in the whole slice (Fig. 4a, Fig. S10). As the paired single-cell reference (produced from the same sample as the ST data) is not available, we used as reference another human heart snRNA-seq atlas [37] consisting of 10 major cell types including cardiomyocytes to less common adipocytes and neuronal cells (Fig. S11). SpatialScope learned the distribution of the gene expression in each cell type from this atlas via a deep generative model. The “pseudo-cells” generated using this learned model are indistinguishable from existing real cells in the reference data (Fig. 4c), laying the foundation for SpatialScope to accurately resolve spot-level ST data containing multiple cells to single-cell resolution. The overall cell-type composition across all spots identified by SpatialScope was highly consistent with that of the snRNA-seq reference from the same tissue type (the heart left ventricle) (Fig. 4b). These results further validate the performance of SpatialScope on real data beyond the simulated dataset. Alignment-based methods, on the other hand, did not provide satisfactory estimations of cell-type composition. Tangram mis-identified many cells in the left ventricle as atrial cardiomyocytes, and CytoSPACE was unable to identify pericytes, a major cell type in human heart tissue. The SpatialScope estimated cell-type compositions remained highly consistent even when using different non-paired human heart snRNA-seq atlases as reference (Fig. S12), suggesting that it is robust to the choice of reference data in real data analyses, which is important during practical implementation.

**Figure 4:**
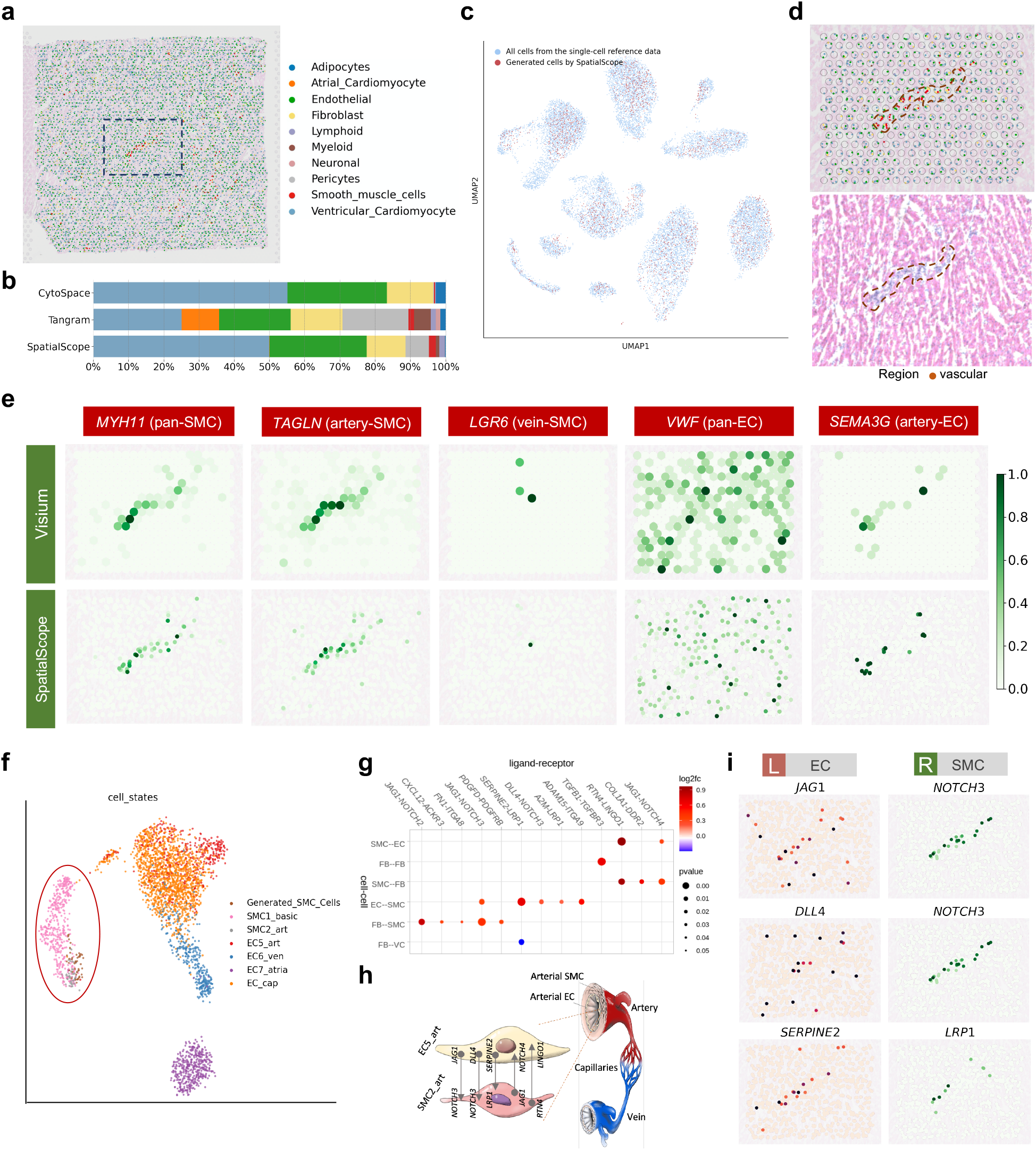
Analysis of vascular region in spot-based human heart ST data. **a**, Cell type identification result at single-cell resolution for whole slice 10x Visium data of human heart using SpatialScope. The background is H&E staining of human heart. Black dotted line indicates ROI. **b**, Inferred cell type compositions across the whole slice by SpatialScope, Tangram and CytoSPACE. **c**, UMAP of scRNA-seq reference data (blue dots) and the pseudo cells (red dots) generated by the deep generative model. **d**, Top, cell type identification result at single-cell resolution in ROI. Bottom, H&E staining of the heart ROI. Black dotted line indicates vascular location. **e**, Expression of SMC and EC marker genes in raw Visium data (top) and single-cell transcriptomes generated by SpatialScope (bottom). **f**, The UMAP plot of SMCs in refined single-cell resolution spatial data generated by SpatialScope and cells from all subgroups of SMC and EC in snRNA-seq reference. Red circle highlights the overlap of inferred SMCs with real arterial SMC rather than venous ones. **g**, Dot plot of ligand-receptor pairs that exhibit spatially resolved cell-cell communications inferred from SpatialScope generated single-cell resolution spatial data. SMC: smooth muscle cells; EC: endothelial cells; FB: Fibroblast; VC: ventricular cardiomyocyte. **h**, Schematic of the vascular cells and inferred cell-cell interactions between SMC and EC in the arteries. **i**, Visualization of molecular interactions between EC and SMC using single-cell resolution gene expression profiles generated by SpatialScope. The scatter plot shows the expression level of ligand-receptor pairs in EC and SMC, respectively.

That SpatialScope can construct pseudo-cells with inferred gene expressions offers a unique advantage over other methods: through deep learning we recover additional information from each spot that is missing in the original ST data due to dropouts of low expression genes, and this enables statistically meaningful analysis of relative expression between cells (Fig. 4e). To illustrate this feature, we focused on a region of interest (ROI) that shows a spatial pattern characterized by vascular cells. Figure 4d shows that the SpatialScope inferred smooth muscle cells (SMC) accurately reside in the areas containing vascular structures, as marked by the pan-SMC marker gene *MYH11* in the original ST data and by the H&E staining in the histological image (Fig. 4d; Fig. 4e). In comparison, alignment-based methods Tangram and CytoSPACE were unable to identify SMCs in this region (Fig. S13a). Cell type deconvolution methods RCTD and spatialDWLS performed better and correctly identified SMCs, while CARD and Cell2location incorrectly identified many endothelial cells (EC) and atrial cardiomyocytes, respectively (Fig. S13b). The SpatialScope inferred results also indicate that this region has a much higher expression level of *TAGLN* than *LGR6* (Fig. 4e). In both brain [39] and cardiac vasculature [37], *TAGLN* was previously found to be highly expressed in arteriole SMCs while much lower in venous SMCs, and high *LGR6* expression was found to be associated with venous SMC [39]. Based on these previous atlas studies, which labelled *TAGLN*-high/*LGR6*-low SMCs as arterial, our result classifies this region as being arterial rather than venous. It is worth noting that the same conclusion could also be drawn from the raw ST data as their expression patterns were highly similar (Fig. 4e), but the single-cell resolution ST generated by SpatialScope allowed us to significantly increase the confidence of the conclusion. To see this, we projected the inferred single-cell level gene expression profiles of SMCs in this region onto the UMAP of all SMCs and ECs in the snRNA-seq reference, obtaining a global view of SMCs identified by SpatialScope. This reveals that the global gene expression of inferred SMCs are clustered with real arterial SMC rather than venous ones (Fig. 4f), indicating that SpatialScope accurately identified the arterial SMC. Other methods could not distinguish these subtypes (Fig. S13c).

Spatially resolved single-cell expression profiles inferred by SpatialScope can further facilitate downstream analysis, for example in exploring cell-cell communication between ECs and SMCs in arteries (Fig. 4h). We applied Giotto [8] to identify statistically significant ligand-receptor (LR) interactions between these two cell types when in close proximity (Fig. 4g) and found LR expression patterns that are consistent with previous studies [37]. Spatial co-expression patterns of these LR pairs was also verified by visual inspection (Fig. 4i, Fig. S14). We further noted that the interacting ECs are arterial, marked by *SEMA3G* expression (Fig. 4e), which is concordant with our previous observation that these SMC are the arterial subtype. Among the LR-pairs we identified as significant, Notch receptor–ligand interactions (e.g., *JAG1-NOTCH3*, *DLL4-NOTCH3*) are known to be essential for regulating vascular smooth muscle proliferation and differentiation [40, 41], and *SERPINE2-LRP1* has been reported to act as a protector of vascular cells against protease activity [42, 43]. *RTN4-LINGO1* is commonly detected in brain tissue due to its importance in regulating neuronal development [44]. With the inferred gene expression profiles with single-cell resolution, here our results indicate that *RTN4-LINGO1* has a spatially strong co-expression pattern in the human heart vascular region (Fig. S14). The RTN family of genes are also known by another name, the Nogo family, and *RTN4* protein products are widely expressed in many cell types but most highly expressed on the surface of glial cells. Both *RTN4* and *LINGO1* are found to be expressed in multiple cell types including smooth muscle cells and endothelial cells [45]. Literature has reported the interaction of this ligand-receptor in the brain [44, 46, 47], and the Nogo-B isoform was found to be important in regulating vascular homeostasis and remodeling in mouse models [48]. Further research is need to uncover the tissue-specific mechanisms and roles of the *RTN4-LINGO1* pair in the human heart.

### SpatialScope enables accurate correction of dropouts in spot-level ST data

Various spatial technologies differ in their resolution; as an example, Slide-seqV2 can achieve a higher spatial resolution than the Visium technology but the trade-off is a lower transcript capture rate [49]. For example, in a cerebellum Slide-seq V2 dataset with 10,975 cells within 8,952 spots (Fig. 5a), 98.55% entries of the gene expression matrix are zero and the median UMI counts per spot is about 300 [12]. In this dataset, some marker genes exhibit unusual sparsity (Fig. 5d, Fig. S15), with total UMIs across all spots as low as 25 in some cases (*Klf2*). We can also leverage SpatialScope to correct the low-detection in situ transcripts, inferring the missing signals using the gene expression distribution learned from the single-cell reference (Fig. S16). As shown in Fig. 5a, SpatialScope correctly assigned cell type labels and captured the three-layer architecture (molecular layer, Purkinje cell layer and granular layer) of the cerebellum [50, 51]; these high resolution single-cell level results are consistent with spot-level RCTD results [12]. Other methods produced noisy results and even incorrectly estimated cell type proportions: Cell2location missed most Astrocytes; SpatialDWLS wrongly detected a large number of Fibroblasts in the Purkinje cell layer; alignment-based methods Tangram and CytoSPACE could not reconstruct the granular layer, suggesting that alignment-based methods are not robust to low capture rate data (Fig. S17).

**Figure 5:**
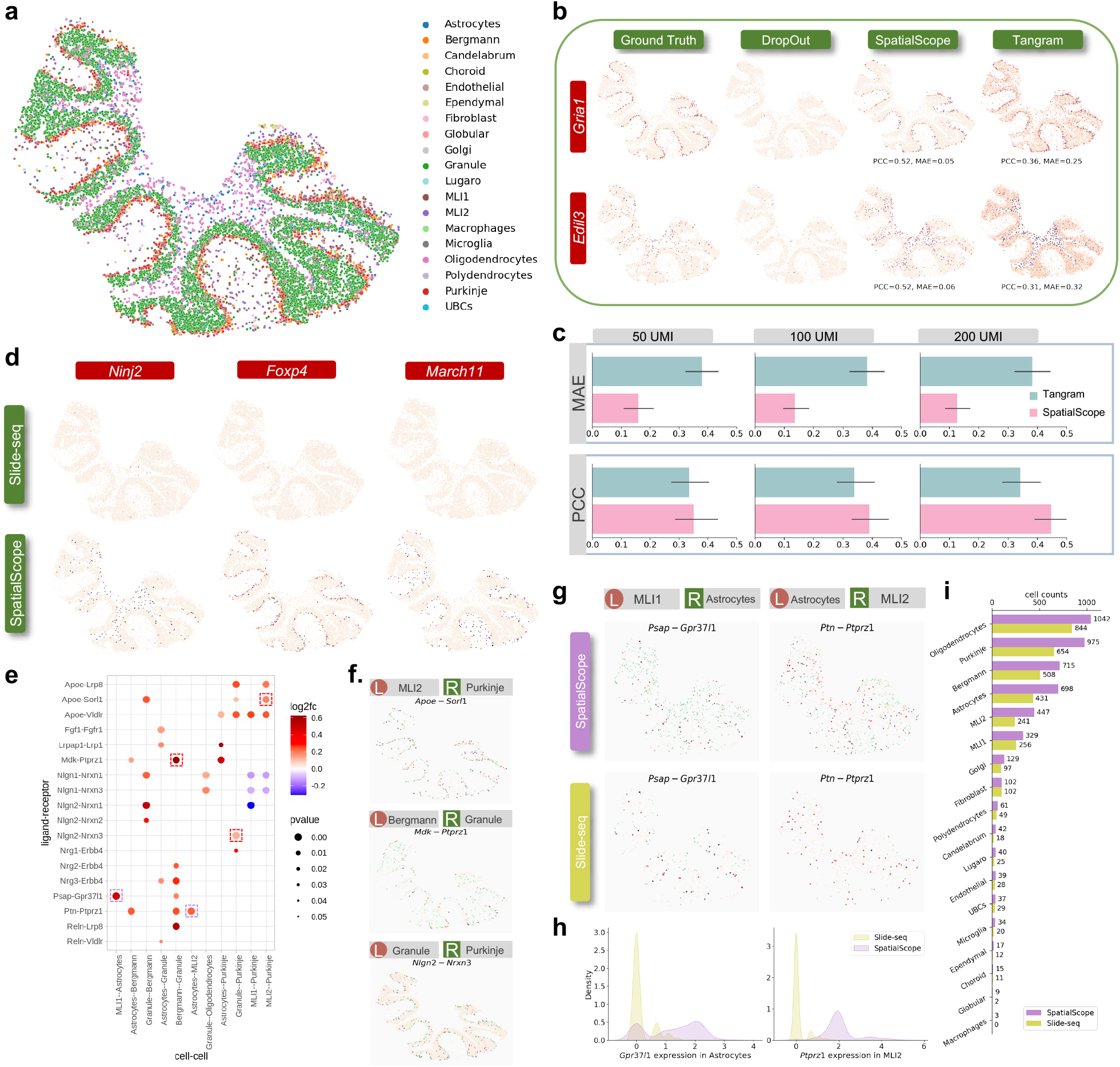
Application of SpatialScope to Slide-seq V2 cerebellum data. **a**, Cell type identification results of Slide-seq V2 cerebellum data by SpatialScope. **b**, Correction of two simulated low-quality spatial measurements genes. Slide-seq V2 measurements (first column), simulated low-quality spatial measurements after processing subsampling (second column), SpatialScope-corrected gene expression level (third column), and Tangram-corrected gene expression level (Fourth column). **c**, Accuracy of dropouts correction on simulated low expression genes under different subsampling level. Two different metrics are used to evaluate correction accuracy in terms of the similarity between the corrected expression and true expression: Pearson’s correlation (PCC) and Mean Absolute Error (MAE). **d**, Correction of low-quality spatial measurements on real data. Slide-seq measured (top) and SpatialScope corrected genes (bottom). **e**, Dot plot of ligand-receptor pairs that exhibit spatially resolved cell-cell communications inferred from corrected gene expression profiles by SpatialScope. Red frame indicates ligand-receptor pairs further visualized in **f**. Purple frame indicates ligand-receptor pairs that are newly found after dropouts correction by SpatialScope and further visualized in **g**. **g**, Visualization of ligand-receptor expression in both corrected Slide-seq data by SpatialScope (first row) and raw Slide-seq data (second row). **h**, Comparison of gene expression level for ligand/receptor between raw Slide-seq data and corrected Slide-seq data by SpatialScope, displayed by density plots. **i**, Comparison of cell counts between raw Slide-seq data and corrected Slide-seq data by SpatialScope for each cell type.

To evaluate the performance of dropouts correction for SpatialScope, we randomly subsampled the UMIs of existing marker genes with high-capture rates to mimic the technical “dropouts”, and then applied SpatialScope to check if we could accurately recover the spatial expression patterns of these marker genes (Fig. 5b).

Specifically, we selected 22 marker genes with high-capture rates, where the median UMIs is about 3,600 across all spots, and then subsampled their UMIs to 50, 100, 200. Notably, SpatialScope showed the best performance of the dropout correction in all settings in terms of mean absolute error (MAE) and PCC (Fig. 5c, Fig. S18). As the subsampled UMIs increased, SpatialScope further improved the correction accuracy but the performance of Tangram plateaued. We then used SpatialScope to correct low-capture genes in Slide-seq data. A close inspection of the corrected sparse marker genes showed clear expression patterns concordant with spatial cell type organization, indicating that SpatialScope can effectively address the dropout issue in Slide-seq ST data (Fig. 5d, Fig. S15).

Low capture rates means that many ligand and receptor pairs are also sparsely captured, making it difficult to perform relevant downstream analyses. The SpatialScope-corrected Slide-seq data imputes genes with low-capture rates, enabling further calculation of cell-cell communications. For example, the cellular communication mediated by *Psap* and *Gpr37l1* between molecular layer interneuron type 1 (MLI1) cells and astrocytes was only detected in the corrected data (Fig. 5g). Astrocytes are reported to have neuroprotective effects on neurons through the *Gpr37l1* pathway [52, 53], supporting the cell-cell interactions we identified in the corrected data; In contrast, raw Slide-seq data was too sparse to detect this (Fig. 5h-i). We detected many cellular interactions that are concordant with existing literature (Fig. 5e). For example, basket cells (e.g., MLI1 and MLI2) in the molecular layer of the cerebellum is known to have a powerful inhibitory effect on Purkinje cells [50], and we indeed found the *Apoe-Sorl1* interaction between these two cell types (Fig. 5f). Notably, both Apoe and *Sorl1* are genes associated with Alzheimer’s disease risk, and play roles in regulating the clearance of amyloid protein *β* [54]; the interacting cell types detected by SpatialScope may help to elucidate the underlying genetic etiology behind Alzheimer’s disease.

### SpatialScope accurately imputes unmeasured genes on single molecule imaging-based ST dataset to enable global differential gene expression analysis

Finally, we investigated how SpatialScope could leverage deep generative models to impute unmeasured genes in image-based spatial transcriptomics data that only measures a panel of selected genes. We analyzed a MERFISH dataset, where the expression profiles for 254 genes were measured in 5,551 single cells in a mouse brain slice from the primary motor cortex (MOp) [25]. To perform cell type identification and learn the distribution of single-cell gene expression, we used a paired droplet-based snRNA-seq profiles from mouse MOp as the reference dataset (Fig. 6b) [16]. SpatialScope successfully learned the gene expression distribution of the single cell reference data (Fig. 6b), laying the groundwork for inferring the expressions of unmeasured genes. Using the 252 genes that were targeted by MERFISH and that overlap with snRNA-seq reference data, we assigned cell type labels for each cell on the slice. As shown in Fig. 6a, SpatialScope successfully reconstructed the known spatial organization of cell types in the MOp of the brain cortex. Specifically, glutamatergic neuronal cells showed distinct cortical layer patterns while GABAergic neurons and most non-neuronal cells were granularly distributed (Fig. 6a).

**Figure 6:**
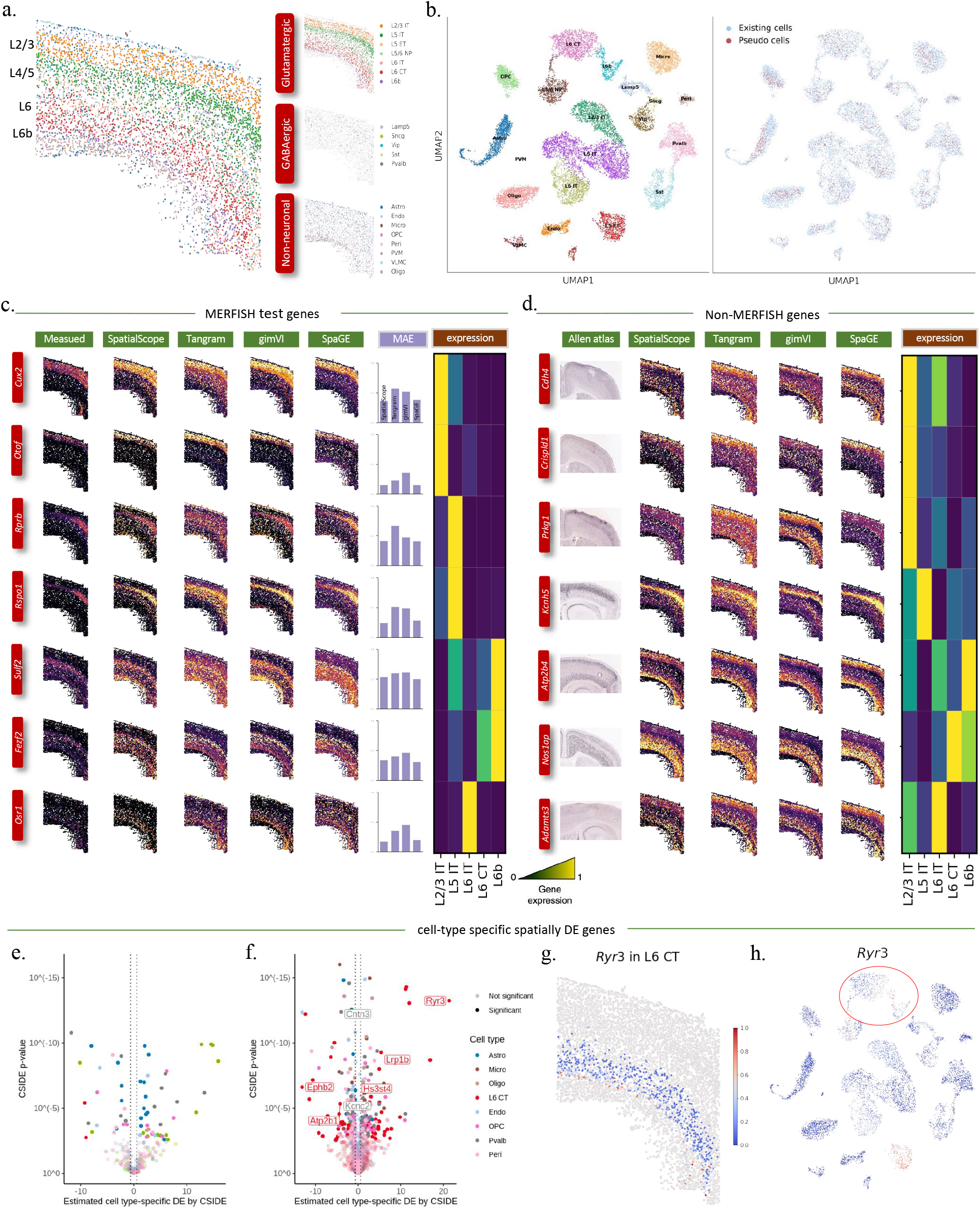
Application of SpatialScope to MERFISH data. **a**, Cell type identification results of MERFISH MOp data by SpatialScope. Cell type identification results in each of three major categories are shown on the right. **b**, UMAP plots of snRNA-seq reference data (left). UMAP of single cell reference data and the pseudo cells generated by deep generative model (right). **c**, Measured and imputed expressions of known spatially patterned genes in the MERFISH dataset. Each row corresponds to a single gene. First column from the left shows the measured spatial gene expression in the MERFISH dataset, while the second to fifth columns show the corresponding imputed expression pattern by SpatialScope, Tangram, gimVI, SpaGE. The imputation accuracy was evaluated by MAE and displayed with bar plots (sixth column). The marker gene expression signatures in snRNA-seq reference were displayed with heatmap plot (seventh column). **d**, Measured and imputed expressions of Non-MERFISH genes. Each row corresponds to a single gene. First column from the left shows the ISH images from the Allen Brain Atlas, while the second to fifth columns show the corresponding imputed expression pattern by SpatialScope, Tangram, gimVI, SpaGE. The gene expression signatures in snRNA-seq reference were displayed with heatmap plot (sixth column). **e**, Volcano plot of C-SIDE cell-type specific spatially DE results for MERFISH dataset, where a total of 252 genes in MERFISH dataset are considered. Color represents cell type, a subset of significant genes is labeled, and dotted lines represent 1.5x fold-change cutoff. **f**, Volcano plot of C-SIDE spatially DE results for imputed MERFISH dataset by SpatialScope, where a total of 1938 genes including genes in original MERFISH dataset and imputed Non-MERFISH genes by SpatialScope are considered. **g**, Spatial Visualization of Ryr3, identified by C-SIDE as DE in L6 CT. Color shows the expression change of *Ryr3* across L6 CT. **h**, The expression profile of *Ryr3* in single cell reference data. The expression level change of *Ryr3* in L6 CT cell type are outlined in the red circle.

Then we compared the performances of SpatialScope and other methods in imputing spatial expression patterns. Specifically, we split the genes overlapping with the reference data into training and testing genes. Then, all the compared methods used the training genes as inputs to predict the gene expression patterns of the testing genes. Here we selected the cortical layer-specific makers (*Cux2*, *Otof*, *Rorb*, *Rspo1*, *Sulf2*, *Fezf2* and *Osr1*) as testing genes for better visualization of the predicted spatial gene expression patterns. As correlation metrics are often sensitive to noise in gene measurements [16], we evaluated the imputation performance by computing MAE between the real measurement and the predicted gene expression of the testing genes. Overall, we showed that SpatialScope achieved state-of-the-art imputation accuracy with an average MAE as low as 0.126, with 34% improvement compared to Tangram (0.191) (Fig. 6c). For example, as a marker gene of the L5 IT layer, the spatial gene expression of *Rspo1* imputed by SpatialScope is in accordance with the real measurement specific to the L5 layer.

In contrast, other methods overestimated the expression of *Rspo1* outside the L5 layer. Next, we used all overlapped genes as training genes and evaluated the imputation performance of non-MERFISH genes using an ISH dataset [55] as validation, and found that other methods tended to overestimate the spatial expression in some layer-specific marker genes (e.g., *Cdh4*, *Prkg1*) (Fig. 6d).

SpatialScope increases the gene throughput of MERFISH from 254 to thousands of genes, enabling us to conduct wide-ranging downstream analysis such as detection of spatially differentially expressed (DE) genes. We first applied a recently developed tool, C-SIDE [56], to detect cell-type specific spatially DE genes on the imputed MERFISH dataset. As expected, compared to 63 cell-type specific spatially DE genes detected in MERFISH genes under an FDR of 1% (Fig. 6e), the number of significant genes with FDR < 1% increases to 293 by incorporating the imputed Non-MERFISH genes (Fig. 6f). For example, *Ryr3* encodes a calcium release channel that affects cardiac contraction, insulin secretion, and neurodegeneration by altering the levels of intracellular *Ca*^2+^ [57]. The expression of *Ryr3* in L6 CT shows a spatial pattern that coincides with the L6b cell boundary (Fig. 6g and Fig. S19a), suggesting the potential communication between L6 CT and L6b through *Ryr3*. Interestingly, the expression signature of *Ryr3* in the single-cell reference data also suggests its diverse expression in L6 CT and L6b (Fig. 6h), and the transition region between these two cell types in single-cell reference shows higher expression, which is perfectly concordant with what we observed in the imputed spatial expression pattern of *Ryr3* in MERFISH data. This concordance highlights the value of SpatialScope in integrating the merits of both single cell reference and lower throughput high precision spatial transcriptomic data such as MERFISH. Next, we considered the spatially DE genes across the entire MERFISH data instead of restricting to specific cell types. We applied SPARK-X [30] and identified 243 genes that exhibit spatially DE patterns in a global perspective, which was 2.3 times more than the number of DE genes detected in MERFISH genes (Fig. S19c). Visualizing a few representative non-MERFISH DE genes clearly shows their significantly spatially distinct expression patterns (Fig. S19d). For example, *Lingo2* encodes a transmembrane protein that positively regulates synapse assembly [58], and the genetic variants of *Lingo2* have been reported to be linked to Parkinson’s disease (PD) and essential tremor (ET) [59, 60]. The spatial expression pattern of *Lingo2*, highly expressed in the upper cortical layer, imputed by SpatialScope may shed light on the genetic etiology of PD/ET in brain MOp.

## Discussion

Fine-grained cell gradients are critical for understanding cellular communication within tissues, which requires that ST technologies achieve the detection of the whole transcriptome at singlecell resolution. However, existing ST technologies often have limitations either in spatial resolution, capture rate of the genes, or the number of genes that can be profiled in one experiment. Here we developed a unified framework SpatialScope to address these limitations.

SpatialScope is applicable to different ST technologies and can achieve several important functions. First, SpatialScope recovers single-cell resolution data from seq-based technologies (e.g. 10X Visium) that do not have single-cell resolution. Consequently, single-cell resolution ST data produced by SpatialScope enables the detection of spatially resolved cellular communication, which is almost impossible for ST data that does not have cellular resolution. Spatially resolved cell-cell communications between each paired cell mediated by ligand–receptor interactions can be robustly inferred and visualized, leading to decoding spatial inter-cellular dynamics in tissues. Second, SpatialScope improves the power and precision of molecular interaction by correcting for genes that has low capture rate in higher-resolution spatial data, such as Slide-seq. Some signals missing in the raw ST data can be detected after the correction for dropouts by SpatialScope. Third, SpatialScope imputes unmeasured genes for image-based ST technology that cannot measure the whole transcriptome, such as MERFISH, allowing the discovery of more biologically meaningful signals. Fourth, SpatialScope can integrate multiple slices of ST data, which enables better cell type identification and the detection of cell-cell communication by increasing the effective sample size.

SpatialScope’s ability to accurately and robustly resolve the spot-level data towards higher resolution and expand from signature to transcriptome-wide scale expression comes from the fact that SpatialScope leverages the deep generative model to approximate the distribution of gene expressions accurately from the scRNA-seq reference data. Rather than directly applying learned distribution from single-cell reference data, SpatialScope accounts for the platform effects between single-cell reference and ST data. Cell type identification and gene expression decomposition results would not be satisfactory if the platform effects are not appropriately corrected. With these innovations in its model design, SpatialScope serves as a unified framework which is applicable to ST data from various platforms. In the step of cell type identification, SpatialScope leverages spatial information to improve the accuracy of cell type identification for each single cell. The inclusion of spatial information also allows straightforward extrapolation of SpatialScope to data with multiple slices, where 3D spatial information across slices is well exploited.

While SpatialScope has shown its superior performance, it can be time-consuming to train a generative model to approximate the distribution from single-cell reference data. In our real data analysis, the training time of the deep generative models is measured in hours or days. Although the generative model can be pre-trained using single-cell atlas data sets and we only need to train each dataset once, ST data analysis can still benefit a lot from more computationally efficient methods that reduce the computational complexity for learning deep generative models [61].

While our work focused on the analysis of cell-cell communications and spatially DE genes detection, we anticipate that refined single-cell resolution spatial transcriptomic data generated by SpatialScope can be very useful in many other downstream applications. Examples include unraveling spatiotemporal patterns of cells [62], analysis of cellular interactions between tumor and immune cells in disease or cancer tissue, and inference of differentiation trajectories [63]. We believe that SpatialScope can serve as a very useful tool in providing single-cell resolution ST data, facilitating detailed downstream cellular analysis, and generating biological insights.

## Methods

To characterize spatially-resolved transcriptome-wide gene expression at single-cell resolution, we introduce SpatialScope as a unified framework to integrate single-cell and ST data. For 10x Visium ST data, the SpatialScope method comprises of three steps: nucleus segmentation, cell type identification, and gene expression decomposition. SpatialScope can also be applied to dropout correction for Slide-seq data and transcriptome-wide gene expression imputation for image-based ST data, such as MERFISH data.

### Nucleus segmentation

Accurate segmentation of nuclei/cells in microscopy images is an important step to locate cells and count the number of cells within a spot. In this study, we compared the performance of two deep learning tools for nucleus segmentation, i.e., StarDist [64] and Cellpose [65]. We observe that StarDist performs better for hematoxylin and eosin (H&E)-stained histological images of Visium data (see Supplementary Fig. S20 for more details). Therefore, we use StarDist as the default tool to perform nucleus segmentation on the H&E-stained histological image. After segmentation, we denote *M_i_* as the number of detected cells at the i-th spot, *i* = 1,…, *I*, where *I* is the total number of spots.

### Cell type identification

Suppose we have *K* cell types in a single-cell reference data. The expression counts of *G* genes have been measured to capture the whole transcriptome in the scRNA-seq data. Let *k_i,m_* ∈ {1, 2,…, *K*} be the cell type of the *m*-th cell at spot *i*, where *m* = 1,…, *M_i_*. Our goal is to infer the cell type vector **k**_*i*_ = {*k_i,m_*} at spot *i* by integrating scRNA-seq and ST data.

As inspired by RCTD [12], we consider the following probabilistic model for cell type identification in ST data by incorporating scRNA-seq reference data,

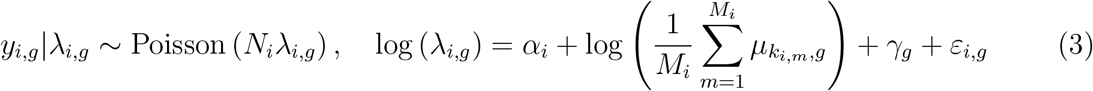

where *y_i,g_* is the observed gene expression counts of gene *g* at spot *i*, *N_i_* is the total number of unique molecular identifiers (UMIs) of spot *i*,λ_*i,g*_ is the relative expression level of gene *g* at spot *i*, *M_i_* is the number of cells in spot *i* inferred from the last step, 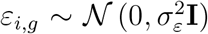 is a random effect to account for additional noise, and *μ_k,g_* represents the mean expression level of cell type *k* and gene *g*, which can be estimated from annotated single-cell reference data. Both *γ_g_* and *α_i_* are designed to address the batch effect between single-cell reference and ST data. More specifically, 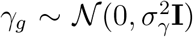 represents a gene-specific random effect to account for expression differences of a gene *g* between single-cell and ST platforms, and *α_i_* is the spot-specific effect to account for differences of a gene set across platforms.

Recall that the RCTD model is given as 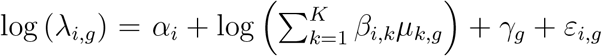, where *β_i,k_* is the proportion of cell type *k* at spot *i*. Our model differs from RCTD in the term 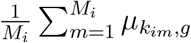, which is the average of the mean expression level of cell types corresponding to the *M_i_* cells at spot *i*. In other words, our model can be viewed as a discrete version of RCTD which was developed to estimate the continuous cell type proportions. The benefits of our discrete version are two-fold. First, given the accurate number of detected cells from image segmentation, it allows us to achieve cell type identification at single-cell resolution. Second, it also enables the incorporation of spatial smoothness constraints to improve the accuracy of cell type identification. In contrast, RCTD can only impose the simplex constraint (i.e., 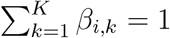 and *β_i,k_* ≥ 0) when estimating *β_i,k_*, leading to suboptimal results. To incorporate spatial smoothness in the distribution of the cell types, we assume a prior given by the Potts model for cell types **K** = {*k_i,m_*},

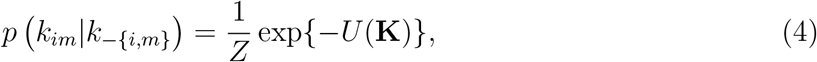

where 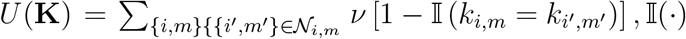 is the indicator function which equals to 1 when 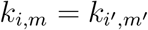 and 0 otherwise, *Z* is a normalization constant, 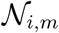 is the set of neighbors of the *m*-th cell in spot *i* and –{*i*, *m*} denote all the cells other than (*i*, *m*) cell. Parameter *ν* controls the smoothness of cell type labels. The larger the *ν*, the smoother the cell type labels.

Now we develop an iterative algorithm to identify cell type label *k_i,m_* based on maximum a posterior (MAP) estimate, where *i* = 1, 2,…, *I*, and *m* = 1, 2,…, *M_i_*. Meanwhile, we are also interested in *γ_g_* which will used to correct gene-level batch effects between different platforms. First, we estimate *μ_k,g_* by calculating mean expression of gene *g* and cell type *k* from single-cell reference data. Next, we follow RCTD’s strategy to accurately approximate *γ_g_* by viewing ST data as a bulk RNA-seq data using the convenient property of Poisson distribution [12]. Other parameters, including *α_i_*, can be obtained accordingly (see Supplementary Methods for details). Then we iteratively find MAP estimate of {*k_i,m_*} and the estimate of *σ_ε_*. The derivation of the MAP estimate for {*k_i,m_*} is as follows. Let 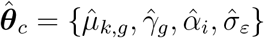 be the collection of these estimates in the above cell type identification model, where *k* = 1, 2,…, *K*, *g* = 1, 2,…, *G*, and *i* = 1, 2,…, *I*. Given 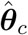, we can obtain the MAP estimate for **K**:

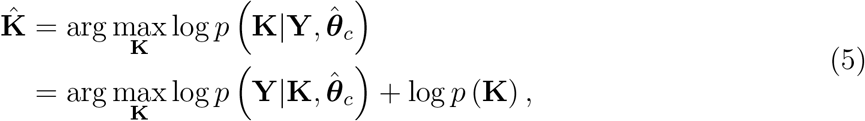

where **Y** = {*y_i,g_*} represents all observed gene expression counts. The term 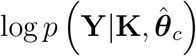 is the likelihood term of model (3) and log *p* (**K**) is the prior term given by Potts model (4). For computational efficiency and scalability, we adopt iterative-conditional-mode-based scheme [66] to infer **K** by updating two labels 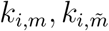 at a time. Then the distribution becomes,

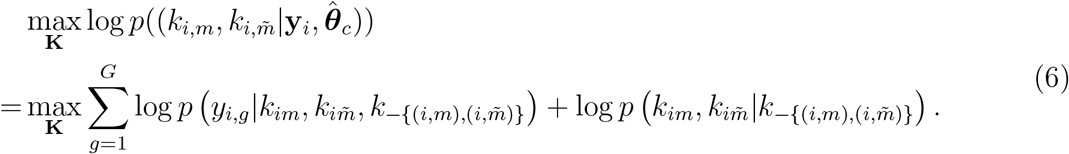

By finding the MAP estimate, we not only use information from gene expression levels *y_i,g_* to determine the cell type labels *k_i,m_*, but also incorporate information from its neighbors. We use the hyperparameter *ν* = 10 as the default setting, and we found that *ν* = 10 ~ 100 yielded similar results. The details of the optimization process are given in Supplementary Methods.

### Gene expression decomposition

We first learn a score-based generative model to approximate the expression distribution of different cell types from the single-cell reference data. Then we use the learned model to decompose gene expression from the spot level to the single-cell level, while accounting for the batch effect between single-cell reference and ST data.

#### Learning conditional score-based generative models from single-cell reference data

There are two major challenges in learning score-based generative models for the scRNA-seq data. First, while score-based generative models [22, 21, 67, 68, 69] can accurately approximate the distribution of images, the nature of the scRNA-seq count data, such as sparsity in the expression matrix, may hinder the capacity of score-based generative models. Second, as given in Eq. (2), we need to learn a conditional score function ∇_x_ log *p*(**x**|*k*) rather than an unconditional score function ∇_x_ log *p*(**x**), where *k* represents the cell type. The reason for learning a conditional score function has been demonstrated in the supplementary note section 2.7.5. It largely remains unknown how to learn the conditional score function across different cell types using a coherent neural network. Let’s begin with the key idea of learning the unconditional score function, and then show how to learn a conditional score function based on the single-cell reference data.

Consider the vanilla score matching problem which aims to find a neural network **s_*θ*_**(**x**) to approximate 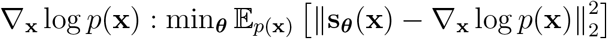, where ***θ*** represents the parameter set of the neural network. The challenge of vanilla score matching comes from the fact that high dimensional data **x** often tends to concentrate on a low dimensional manifold embedded of the entire ambient space. For data points not on the low-dimensional manifold, they would have zero probability, the log of which is undefined. Moreover, the score function can not be estimated accurately in the low density region. Fortunately, these challenges can be addressed by adding multiple levels of Gaussian noise to data. The perturbed data with Gaussian noise will not concentrate on the low-dimensional manifold, and the multiple levels of noise will increase training samples in the low-density region. Specifically, a sequence of data distributions perturbed by *L* levels of Gaussian noise is given as 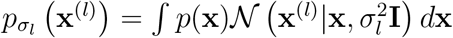, where **x**^(*l*)^ represents a sample perturbed by the noise level 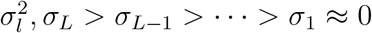. To learn the score function **s_*θ*_** (**x**, *σ_l_*), we consider the following problem,

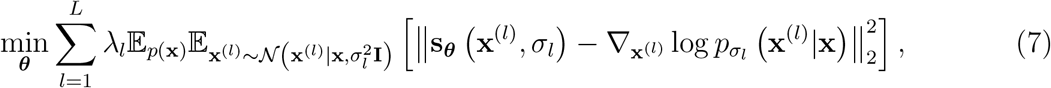

where 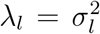 is the weight for noise level *l* and 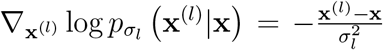. Based on (7), the score function can be estimated by stochastic gradient methods. Let **x**^(*l,t*)^ be the *t*-sample at level *l*. We run Langevin dynamics in (8) from *l* = *L* to *l* = 1 with initialization **x**^(*l,t*=1)^ = **x**^(*l*+1,*t*=*T*)^. In the meanwhile, we progressive reduce of noise level *σ_l_* and decrease the step size *η*:

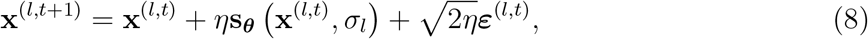

where 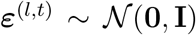. Then the obtained samples **x**^(*l,t*)^ at level *l* = 1, *t* = 1,…, *T*, will approximately follow the target distribution *p*(**x**) because *σ*_1_ ≈ 0.

Now we consider learning our score function conditional on cell types based on scRNA-seq data. For computational stability, we transform the count data to its log scale and remove the mean expression level of each cell type. Specifically, let 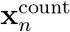 be the gene expression counts corresponding to cell *n* of cell type *k*. The transformation is given as 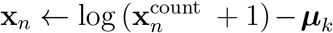, where 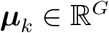 is the mean expression level of cell type *k*. Later on, we will learn the conditional score function based on the transformed expression level. To incorporate cell type information, we consider the following optimization problem:

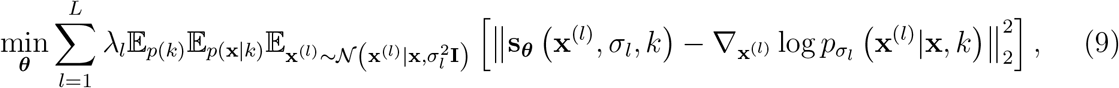

where the score function **s_*θ*_** (**x**^(*l*)^, *σ_l_*, *k*) explicitly takes cell type *k* ∈ {1, 2,…, *K*} as its input. In principle, the score function **s_*θ*_** (**x**^(*l*)^, *σ_l_*, *k*) can be estimated by solving the optimization problem given in Eq. (9). In practice, however, the learning process often tends to largely ignore the cell type information because the neural network will naturally focus on 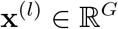 rather than the scalar *k*. To successfully incorporate cell type information, our key idea is to embed cell type information in a vector whose dimension is comparable to **x**^(*l*)^. Therefore, we propose to learn the score function **s_*θ*_** (**x**^(*l*)^, *σ_l_*, ***μ***_*k*_) which takes the mean expression level of cell type *k* as input. The benefits are two-fold. First, ***μ***_*k*_ provides precise information about cell type *k*. Second, 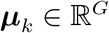 has the same dimension of **x**^(*l*)^ such that it will not be ignored. With this key idea, we can design a novel network architecture for learning the score function **s_*θ*_** (**x**^(*l*)^, *σ_l_*, ***μ***_*k*_). The details of the learning procedure are provided in Supplementary Methods. The learned score function **s_*θ*_** (**x**^(*l*)^, *σ_l_*, ***μ***_*k*_) is then used in “gene expression decomposition” step (next section). We use the score function at 7500 training epoch for all data analysis in the main text. We also investigate the performance of SpatialScope under score-based generative models with different training epochs and recommend that the number of epochs ranges from 5,000 to 10,000 due to the trade-off between the performance and the time cost (supplementary note section 2.7.4).

#### Decomposition with a conditional score-based generative model

Now we show how we obtain gene expression decomposition at single-cell resolution by leveraging the learned score-based generative model. One of the pertinent challenge for decomposition is the batch effects between single-cell reference and ST data. If the batch effects are not appropriately corrected, the decomposition results will not be satisfactory. Therefore, we adjust the batch effects between ST and single-cell reference data before we perform gene expression decomposition. Our batch effect correction includes two steps. Specifically, in the first step, we adjust for the gene-specific cross-platform effects using 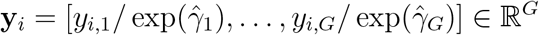, where *y_i,g_* are the observed expression counts of gene *g* at spot *i* and 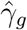 is the batch effect of gene *g* estimated under model (3). In the second step, we account for the difference in sequencing depth by normalizing the total count of **y**_*i*_ to the mean of the total transcript counts of individual cells from single-cell reference data. Next, we show how to decompose the normalized **y**_*i*_, which is corrected for batch effect, into single-cell resolution. Let **X**_*i*_ = [**x**_*i*,1_;…, **x**_*i,M_i_*_] be the expression level in the log scale, where **x**_*i,m*_ is the expression level of the *m*-th cell in spot *i*, and *M_i_* is the number of cells in spot *i* inferred in the nucleus segmentation step. Our goal is to decompose gene expression from the spot-level **y**_*i*_ to the single-cell level **x**_*i,m*_.

Let *p* (**x**|*k_i,m_*) be the gene expression distribution of cell type *k_i,m_*, where the cell type labels for the cells in spot *i*, **k**_*i*_ = {*k*_*i*,1_,…, *k_i,M_i__*} are inferred as in the “cell type identification”. As outlined in the methods overview, we consider the following probabilistic model for gene expression decomposition,

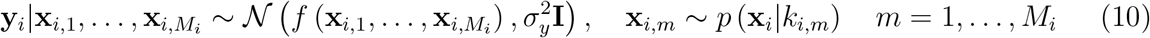

where 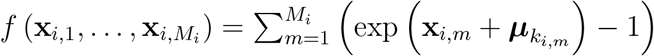 transforms the log-scale expression level to the count scale. To obtain the decomposition, we use Langevin dynamics to get samples from the posterior *p* (**X**_*i*_|**y**_*i*_, **k**_*i*_),

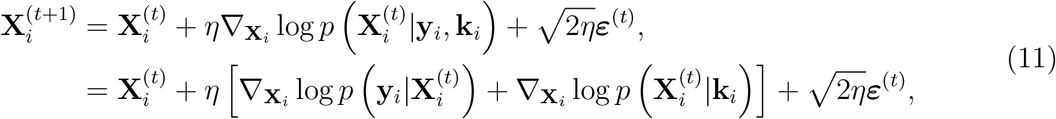

where 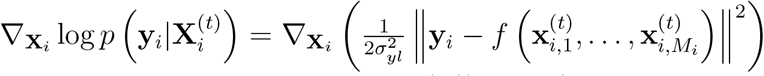 and 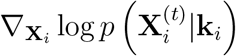 is given by the learned score function **s_*θ*_** (**x**^(*l*)^, *σ_l_*, ***μ***_*k*_). Similar to (8), we progressively reduce noise level *σ_l_* (from *l* = *L* to *l* = 1) and initialize later stage with samples from the previous stage **X**^(*l*,*t*=1)^ = **X**^(*l*+1,*t*=*T*)^,

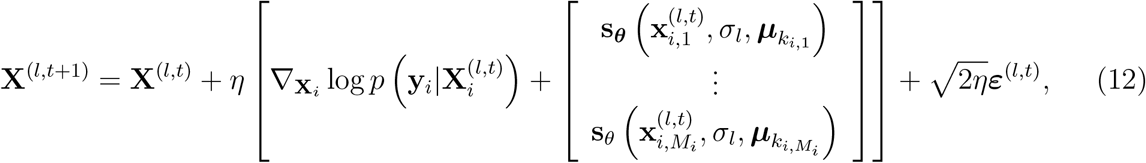

where 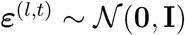. The obtained samples **X**^(*l*,*t*)^ at level *l* = 1, *t* = 1,…, *T*, will be posterior samples from *p* (**X**_*i*_|**y**_*i*_, **k**_*i*_). By averaging samples from Langevin dynamics (12), we use the posterior means as the decomposed gene expression levels at single-cell resolution. The posterior sampling process is summarized in Algorithm 1.

##### Algorithm 1 Annealed Langevin dynamics for gene expression decomposition

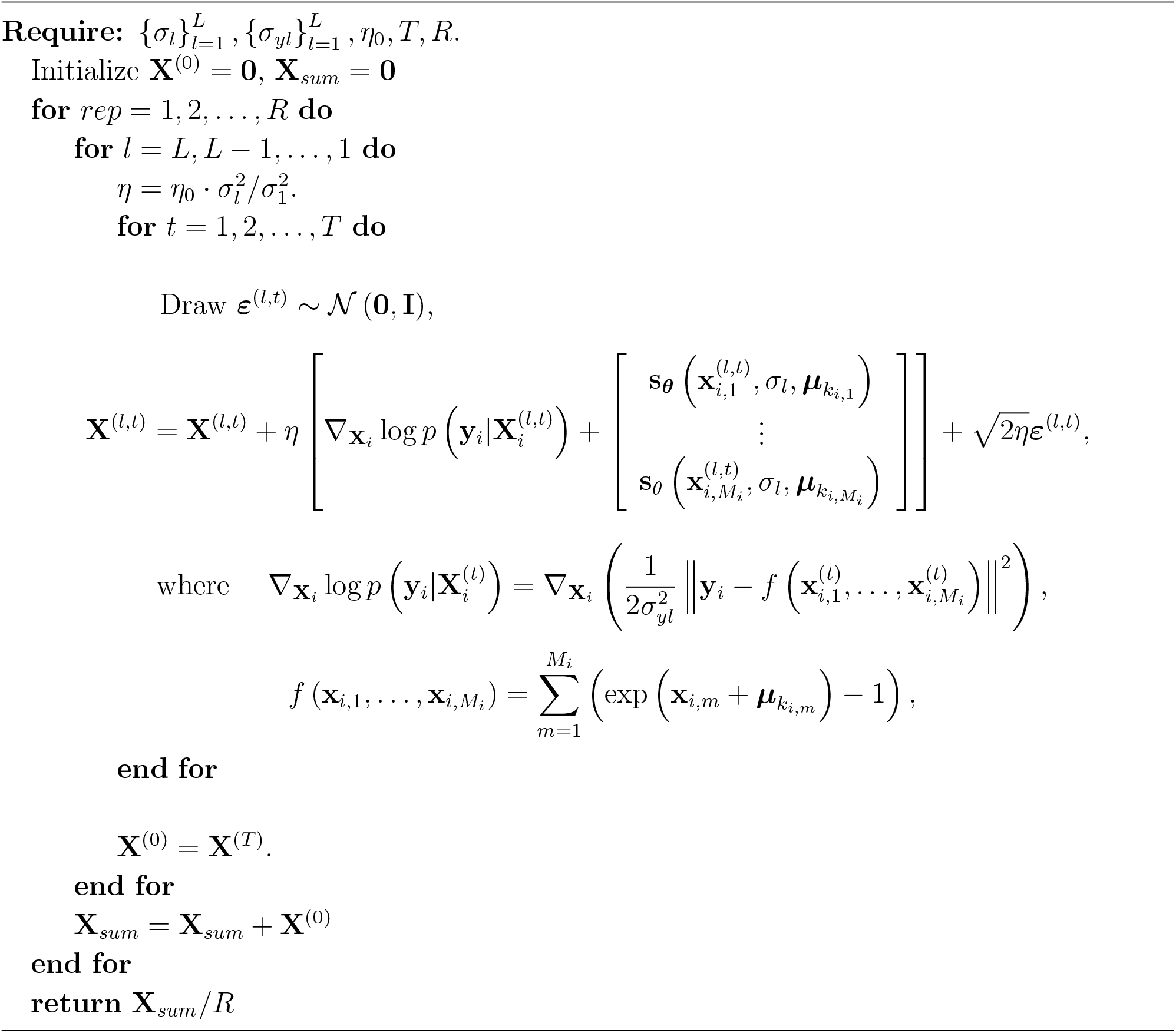

In practice, it is a common situation that there is a large variation in the proportions of different cell types for single-cell reference data. For example, the number of cells varies from tens to thousands for different cell types. To assess the robustness of SpatialScope to unbalanced cell types in single-cell reference data, we compared the performance of gene expression decomposition in different cell types using the MERFISH simulation dataset and found that SpatialScope exhibited robust decomposition performance on unbalanced cell types (supplementary note section 2.7.6).

### SpatialScope for ST data from other platforms

As a unified framework, SpatialScope not only can handle low-resolution ST data with histological images (e.g., 10x Visium), but also can serve as efficient analytical tools for spatial data from other experimental platforms. In this section, we demonstrate that SpatialScope can be applied to perform dropout correction for genes with low-detection rates in Slide-seq data, and imputation for unmeasured genes in MERFISH data or other *in-situ* hybridization-based ST data.

#### Sparse genes dropout correction for Slide-seq data

As a high-resolution approach, the pixel size of Slide-seq can achieve single cell level (10 μm [70]) but it may still contain the mRNA from multiple cells [12]. Slide-seq data can be highly sparse. About 99.46% entries are zero for Slide-seq V1 data and 98.35% for Slide-seq V2 data, compared to about 90% zero counts for 10x Visium data [30]. The framework of SpatialScope can also be applied to correct dropouts in Slide-seq data and recover transcriptome-wide gene expressions at single-cell resolution.

Because of the high resolution of Slide-seq data and lack of histological images, nucleus segmentation step in dealing with 10x Visium data can be skipped for Slide-seq data. As inspired by RCTD [12], we replace the first step nucleus segmentation by singlet/doublet classification, assuming that at most two cell types co-exist within a spot. Next, the steps of cell type identification and gene expression decomposition can be applied similarly as those for 10x Visium data. The correction of dropout for genes with low detection rate is achieved in gene expression decomposition based on the same procedure. Let’s consider the case where a pixel *i* contains two cells, i.e., *M_i_* = 2. In this case, **y**_*i*_ is the aggregated gene expression profiles from two cells, in which the expression levels of some genes are nearly zero. By the same modeling principle as that in (10), we can assume that

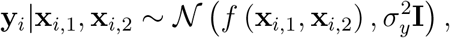

where 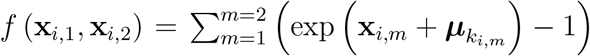. Because SpatialScope first learns the distribution of gene expressions from the single-cell reference data, it can output the posterior means of **x**_*i*,1_ and **x**_*i*,2_ by running Algorithm 1. We then use the posterior means of *x*_*i*,1_ and *x*_*i*,2_ as the de-noised data, where the dropouts are corrected.

#### Imputation for *in-situ* hybridization based ST data

*In-situ* hybridization-based ST data can provide localizations of gene expressions at the cellular level, resulting in single-cell resolution spatial transcriptomics. However, because of the limitation of the indexing scheme [9, 26, 25], the detected spatial transcriptomics by *in-situ* hybridization methods tend to have limited throughput in the number of genes (e.g., tens to hundreds of genes captured by

MERFISH [25]). Therefore, researchers begin to integrate *in-situ* hybridization-based ST data with single-cell reference data to impute the unmeasured genes, providing more complete spatial transcriptome information and cellular structures [18, 16, 17]. By learning the distribution of gene expressions from the single-cell reference data using a score-based generative model, SpatialScope can achieve accurate gene imputation as follows.

Suppose that the expression levels of *G* genes and *G*_0_ genes are measured in the single-cell reference and ST data, respectively. We assume that the set of *G*_0_ genes measured in ST data is a subset of *G* genes in the scRNA-seq data. Let 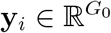 be the measured gene expression counts in ST data after batch effect correction, and 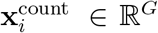 be the true expression at location *i*, respectively. Without loss of generality, we assume that the first *G*_0_ genes in 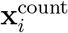 are measured. Then we have

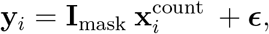

where 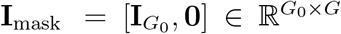, **I**_*G*_0__ is the *G*_0_ × *G*_0_ identity matrix, and 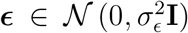 is the measurement noise. As the score function is estimated in the log scale, we denote 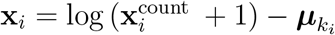 as the log scale expression, where ***μ***_*k_i_*_ is the mean expression level of cell type *k_i_*. Now we have 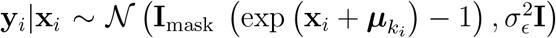 and **x**_*i*_ ~ *p* (**x**_*i*_|*k_i_*). To obtain the imputed expression, we can take samples from posterior *p* (**x**_*i*_|**y**_*i*_) based on the Langevin dynamics,

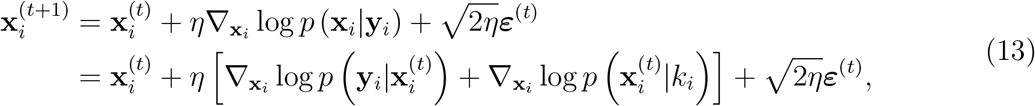

where 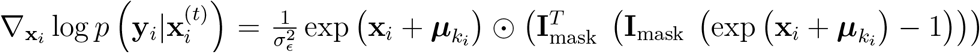 and ⊙ is element-wise product. Using the learned score function **s_*θ*_** (**x**, *σ_l_*, ***μ***_*k*_) given by (9), we begin with a random initialization and then run the Langevin dynamics by progressively reducing noise level *σ_l_*,

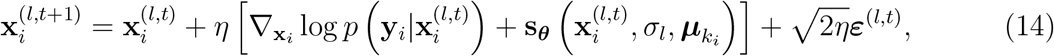

where the initial point at the later stage is given by the sample from the previous stage, i.e., **x**^(*l,t*=1)^ = **x**^(*l*+1,*t*=*T*)^. The obtained samples **X**^(*l,t*)^ at level *l* = 1, *t* =1,…, *T*, will be posterior samples from *p* (**x**_*i*_|**y**_*i*_, *k_i_*). By averaging samples of Langevin dynamics in (14), we use the posterior mean as the imputed gene expression.

### Real data analysis

In this study, we evaluated our SpatialScope on five publicly available spatial transcriptomics datasets.

#### Visium human heart dataset

The human heart sample was from BioIVT Asterand and profiled by 10x Visium that measured the whole transcriptome within 55 μm diameter spots. After removing spots that did not map to the tissue region or with total UMI counts less than 100, 3,813 spots were left for subsequent analysis. We then focused on an ROI of 331 spots that shows a spatial pattern characterized by vascular cells. Through the matched H&E image, we annotated the main vascular structure in the center of ROI that covers 18 tissue spots. For cell type identification, we used a paired human heart snRNA-seq atlas that consists of 10 major cell types, ranging from widespread cardiomyocytes to less common adipocytes and neuronal cells [37]. Following the standard pre-processing procedure, we normalized total counts per cell with median transcript count, then performed *log*(1 + *x*) transformation and selected the top 1000 most highly variable genes and 50 top marker genes for each cell type as training genes. For gene expression decomposition, we first included an additional 876 ligands/receptors provided by Giotto [8] into the training genes for the detection of cellular communication in the downstream analysis. Then we applied the deep generative model to learn the expression distribution of the training genes in the snRNA-seq reference data. Finally, by leveraging the learned single-cell gene expression distribution, we performed the gene expression decomposition for the low-resolution Visium data and generated the single-cell resolved spatial transcriptomics for human heart.

#### Visium mouse brain cortex dataset

The two adjacent sagittal slices of mouse brain anterior tissue were from BioIVT Asterand and profiled by 10x Visium. After removing spots that did not map to the tissue region, 2,695 and 2,825 spatial spots from the two slices were left for subsequent analysis. We first filtered out spatial locations that have less than 100 total read counts. Then, using the matched H&E-stained histological images, we segmented the cerebral cortex regions, resulting in 812 and 794 cortex spots left in slice 1 and slice 2, respectively. Finally, we used the recently developed tool, PASTE [28], to compute a pairwise slice alignment between these two segmented cortex slices, which allowed us to construct an aligned 3D mouse brain cortex ST data. We used mouse brain cortical scRNA-seq data as a reference [27]. This dataset was collected from the mouse Primary visual (VISp) area using the Smart-seq2 technology and contains 14,249 cells across 23 cell types. Similarly, we first performed total counts normalization and log(1 + *x*) transformation, and then selected the top 1000 most highly variable genes and 50 top marker genes from each cell type as training genes. In cell type identification, we incorporated the spatial information in 3D space and thus can produce more reliable spatial priors. Next, in the gene expression decomposition task, we included ligands/receptors and decomposed the gene expressions from the spot-level into the single-cell level using the learned gene expression distribution.

#### Slide-seq v2 mouse cerebellum dataset

The mouse cerebellum dataset was profiled by Slide-seq V2 and measured with the whole transcriptome within 10 μm diameter spots [12]. This dataset consists of gene expression measurements for 23,096 genes and 11,626 spatial spots. We filtered out genes that have zero counts across all spots and filtered out spots with total UMI counts less than 100, leading to 20,141 genes and 8,952 spots for subsequent analysis. As the paired histological images are not available for Slide-seq data, we replaced the first step “Nuclei segmentation” with “Singlet/Doublet classification” inspired by RCTD, which assumes that there are at most two cells per spot as the spot size (10 μm) almost matches the single cell size. Overall, we detected 10975 cells, including 6929 spots contain one cell and 2023 spots contain two cells. Following RCTD, we used a paired mouse cerebellum snRNA-seq data as the reference [71]. This dataset contains 24,387 genes and 15,609 cells from 19 cell types. Similarly, we first performed total counts normalization and log(1 + x) transformation, and then selected the top 1000 most highly variable genes and 50 top marker genes from each cell type as training genes in the cell type identification task. Finally, we generated the corrected high-throughput single-cell resolution Slide-seq data by leveraging the gene expression distribution learned from the snRNA-seq reference.

#### MERFISH MOp dataset

The mouse brain MOp dataset was profiled by the image-based ST approach, MERFISH, with single-cell resolution. This dataset comprised of 254 genes and about 300,000 single cells located in 64 mouse brain MOp slices from 12 different samples [25]. As a concrete example demonstrated in MERFISH paper, we used the slice ‘mouse1_slice180’ from ‘mouse1_sample4’ to evaluate the imputation performance of SpatialScope and the compared methods. We used a paired droplet-based snRNA-seq profiles collected from mouse MOp as the reference, which measures the expression of 26,431 cells across 20 cell types [16]. Following the standard pre-processing procedure, we normalized total counts per cell with median transcript count and performed and *log*(1 + *x*) transformation. Using the 252 genes that overlapped with snRNA-seq reference data as training genes, we first identified the cell type label for each cell in the MERFISH dataset. Then we applied the deep generative model to learn the distribution of gene expressions in the snRNA-seq reference data. Finally, using the learned high-throughput gene expressions distribution, we imputed the gene expressions of unmeasured genes in MERFISH dataset by conditioning on the observed expressions of 252 overlapped genes.

### Downstream analysis

#### Cell-cell interactions

Although ST is believed to be the best suited technology to elucidate cellular/molecular interactions [49], current ST dataset is still limited by either low resolution or low capture rate. Fortunately, the efficient in silico generation of single-cell resolution and high throughput spatially resolved transcriptomics by SpatialScope perfectly solved this issue. As a recently developed tool for detecting cellular communications mediated by ligand-receptor interactions, Giotto was applied on SpatialScope outputs following the protocol (https://rubd.github.io/Giotto_site/articles/tut14_giotto_signaling.html). Specifically, Giotto first ran pre-processing to remove low-quality genes/cells and create a spatial network connecting single cells using Delaunay triangulation, then ran ‘spatCellCellcom’ function to analyze the ligand-receptor signaling with parameters: spatial_network_name = ‘Delaunay_network’. Finally, we selected top and reliable ligand-receptor signaling with the following threshold: p.adj < 0.25, abs(log2fc) > 0.1, lig nr > 10, rec nr > 10, lig_expr > 0.5 & rec_expr > 0.5. For raw Slide-seq data, we detected cell-cell interactions by simply assuming each spot is a single cell whose cell type is determined by the majority proportion. Then the following ligand-receptor signaling analysis by Giotto is identical.

#### Cell-type specific spatially DE genes

We ran C-SIDE to detect cell-type specific spatially DE genes on the MERFISH dataset using “run.CSIDE.nonparametric” function. We followed the guidelines (https://raw.githack.com/dmcable/spacexr/master/vignettes/merfish_nonparametric.html) with parameters: gene_threshold = .001, cell_type_threshold = 10, and fdr = 0.2.

#### Spatially DE genes

Compared to C-SIDE, SPARK-X was developed to consider genes that exhibit spatially DE patterns in a global perspective instead of restricting to specific cell types. We applied SPARK-X to MERFISH dataset using the default parameters following the instruction (https://xzhoulab.github.io/SPARK/03_experiments/). Suggested by SPARK-X, we treated cell type labels as covariates to exclude the spatially DE genes explained by spatial distribution of cell types.

### Compared Methods

For cell type identification task, we compared SpatialScope with two single-cell alignment (Tangram and CytoSPACE) and four deconvolution (SpatialDWLS, RCTD, Cell2location and CARD) methods.

#### Tangram

We followed the instructions of Tangram: https://tangram-sc.readthedocs.io/en/latest. In order to constrain the number of mapped single cell profiles, we set the parameters as mode=“constrained”, target_count=the total number of segmented cells, and density_prior=fraction of cells per spot.

#### CytoSPACE

We followed the guidelines on GitHub repository: https://github.com/digitalcytometry/cytospace. We first used Seurat to obtain a overall cell type composition across all spatial spots, then the estimated fractional composition of each cell type was used as input for alignment.

#### SpatialDWLS

We followed the instructions on the SpatialDWLS website: https://rubd.github.io/Giotto_site/articles/tut7_giotto_enrichment.html. We set the parameter as n_cell = 20.

#### RCTD

We followed the guidelines on the RCTD GitHub repository: https://github.com/dmcable/spacexr. We set doublet_mode = “full”.

#### Cell2location

We followed the guidelines on the Cell2location Github repository: https://github.com/BayraktarLab/cell2location. The single-cell regression model was trained with parameters max_epochs = 250, batch_size=2500, and lr = 0.002. The cell2location model was trained with parameters max_epochs = 30,000.

#### CARD

We followed the guidelines and used the recommended default parameter setting on the CARD GitHub repository: https://github.com/YingMa0107/CARD.

For gene expression decomposition task, we only compared SpatialScope with the two single-cell alignment methods.

#### Tangram

According to the instructions, Tangram only provides a prediction of spot-level gene expression using the mapped single cell profiles through the function “project_genes”. To make it comparable with our SpatialScope in the task of gene expression decomposition, we provided a script that takes the Tangram output as input and generates the single-cell resolution spatial transcriptomics data. Specifically, with the single cells to spatial spots mapping matrix output by Tangram, we first obtained the most probable spot that each single cell belong to. Then we removed the noise cells with mapping probability less than 0.5, and grouped the remaining cells by spot ids. Finally, for each spot, we regarded the grouped cells are mapped single cells from scRNA-seq reference and used their gene expressions as the decomposed single-cell level gene expression profiles.

#### CytoSPACE

CytoSPACE provides the mapped single cells ids from scRNA-seq reference for each spatial spot, we therefore directly used the mapped single cell’s gene expressions as the decomposed gene expression profiles.

For gene expression imputation task, we compared SpatialScope with Tangram, gimVI and SpaGE.

#### Tangram

We followed the instructions of Tangram: https://tangram-sc.readthedocs.io/en/latest. We used function “project _genes” to generate the “new spatial data” with whole transcriptome using the mapped single cell.

#### gimVI

We followed the guidelines on the gimVI website: https://docs.scvi-tools.org/en/0.8.0/user_guide/notebooks/gimvi_tutorial.html. We used the “model.get_imputed_values” function with parameter normalized = False to impute the unmeasured gene expressions.

#### SpaGE

We followed the instructions on the GitHub repository of SpaGE: https://github.com/tabdelaal/SpaGE. We set the parameter n_pv = *N_gene_*/2 if the number of genes used for integration was greater than 50.

## Data availability

For the simulation dataset, the MERFISH data were download from the brain image library (https://doi.brainimagelibrary.org), the STARmap mouse cortex dataset was downloaded from the project page (http://clarityresourcecenter.org). For real data analysis, the 10x human heart and mouse brain cortex datasets were downloaded from the 10x official website (https://www.10xgenomics.com/resources/datasets), and the paired human heart and mouse brain cortex scRNA-seq reference are available from the project page (www.heartcellatlas.org) and (https://portal.brain-map.org), respectively. Both Mouse cerebellum Slide-seq V2 dataset and the paired scRNA-seq reference were download from single cell portal project (https://singlecell.broadinstitute.org/single_cell/study/SCP948).

## Code availability

The SpatialScope software package and source code are available in Github (https://github.com/YangLabHKUST/SpatialScope). We also uploaded all scripts and materials to reproduce all the analyses at the same website.

## References

[1] Angela R Wu, Norma F Neff, Tomer Kalisky, Piero Dalerba, Barbara Treutlein, Michael E Rothenberg, Francis M Mburu, Gary L Mantalas, Sopheak Sim, Michael F Clarke, et al. Quantitative assessment of single-cell rna-sequencing methods. Nature methods, 11(1):41–46, 2014.

[2] Camille Ezran, Shixuan Liu, Stephen Chang, Jingsi Ming, Olga Botvinnik, Lolita Penland, Alexander Tarashansky, Antoine de Morree, Kyle J Travaglini, Kazuteru Hasegawa, et al. Tabula microcebus: A transcriptomic cell atlas of mouse lemur, an emerging primate model organism. bioRxiv, 2021.

[3] Camille Ezran, Shixuan Liu, Jingsi Ming, Lisbeth A Guethlein, Michael FZ Wang, Roozbeh Dehghannasiri, Julia Olivieri, Hannah K Frank, Alexander Tarashansky, Winston Koh, et al. Mouse lemur transcriptomic atlas elucidates primate genes, physiology, disease, and evolution. bioRxiv, 2022.

[4] Sophia K Longo, Margaret G Guo, Andrew L Ji, and Paul A Khavari. Integrating single-cell and spatial transcriptomics to elucidate intercellular tissue dynamics. Nature Reviews Genetics, 22(10):627–644, 2021.

[5] Anjali Rao, Dalia Barkley, Gustavo S França, and Itai Yanai. Exploring tissue architecture using spatial transcriptomics. Nature, 596(7871):211–220, 2021.

[6] Ludvig Larsson, Jonas Frisén, and Joakim Lundeberg. Spatially resolved transcriptomics adds a new dimension to genomics. Nature methods, 18(1):15–18, 2021.

[7] Robert R Stickels, Evan Murray, Pawan Kumar, Jilong Li, Jamie L Marshall, Daniela J Di Bella, Paola Arlotta, Evan Z Macosko, and Fei Chen. Highly sensitive spatial transcriptomics at near-cellular resolution with slide-seqv2. Nature biotechnology, 39(3):313–319, 2021.

[8] Ruben Dries, Qian Zhu, Rui Dong, Chee-Huat Linus Eng, Huipeng Li, Kan Liu, Yuntian Fu, Tianxiao Zhao, Arpan Sarkar, Feng Bao, et al. Giotto: a toolbox for integrative analysis and visualization of spatial expression data. Genome biology, 22(1):1–31, 2021.

[9] Sheel Shah, Yodai Takei, Wen Zhou, Eric Lubeck, Jina Yun, Chee-Huat Linus Eng, Noushin Koulena, Christopher Cronin, Christoph Karp, Eric J Liaw, et al. Dynamics and spatial genomics of the nascent transcriptome by intron seqfish. Cell, 174(2):363–376, 2018.

[10] Jeffrey R Moffitt, Dhananjay Bambah-Mukku, Stephen W Eichhorn, Eric Vaughn, Karthik Shekhar, Julio D Perez, Nimrod D Rubinstein, Junjie Hao, Aviv Regev, Catherine Dulac, et al. Molecular, spatial, and functional single-cell profiling of the hypothalamic preoptic region. Science, 362(6416):eaau5324, 2018.

[11] Bin Li, Wen Zhang, Chuang Guo, Hao Xu, Longfei Li, Minghao Fang, Yinlei Hu, Xinye Zhang, Xinfeng Yao, Meifang Tang, et al. Benchmarking spatial and single-cell transcriptomics integration methods for transcript distribution prediction and cell type deconvolution. Nature Methods, pages 1–9, 2022.

[12] Dylan M Cable, Evan Murray, Luli S Zou, Aleksandrina Goeva, Evan Z Macosko, Fei Chen, and Rafael A Irizarry. Robust decomposition of cell type mixtures in spatial transcriptomics. Nature Biotechnology, 40(4):517–526, 2022.

[13] Vitalii Kleshchevnikov, Artem Shmatko, Emma Dann, Alexander Aivazidis, Hamish W King, Tong Li, Rasa Elmentaite, Artem Lomakin, Veronika Kedlian, Adam Gayoso, et al. Cell2location maps fine-grained cell types in spatial transcriptomics. Nature biotechnology, 40(5):661–671, 2022.

[14] Ying Ma and Xiang Zhou. Spatially informed cell-type deconvolution for spatial transcriptomics. Nature Biotechnology, pages 1–11, 2022.

[15] Rui Dong and Guo-Cheng Yuan. Spatialdwls: accurate deconvolution of spatial transcriptomic data. Genome biology, 22(1):1–10, 2021.

[16] Tommaso Biancalani, Gabriele Scalia, Lorenzo Buffoni, Raghav Avasthi, Ziqing Lu, Aman Sanger, Neriman Tokcan, Charles R Vanderburg, Åsa Segerstolpe, Meng Zhang, et al. Deep learning and alignment of spatially resolved single-cell transcriptomes with tangram. Nature methods, 18(11):1352–1362, 2021.

[17] Romain Lopez, Achille Nazaret, Maxime Langevin, Jules Samaran, Jeffrey Regier, Michael I Jordan, and Nir Yosef. A joint model of unpaired data from scrna-seq and spatial transcriptomics for imputing missing gene expression measurements. arXiv preprint arXiv:1905.02269, 2019.

[18] Tamim Abdelaal, Soufiane Mourragui, Ahmed Mahfouz, and Marcel JT Reinders. Spage: spatial gene enhancement using scrna-seq. Nucleic acids research, 48(18):e107–e107, 2020.

[19] Milad R Vahid, Erin L Brown, Chloé B Steen, Minji Kang, Andrew J Gentles, and Aaron M Newman. Robust alignment of single-cell and spatial transcriptomes with cytospace. bioRxiv, 2022.

[20] Yiming Chao, Yang Xiang, Jiashun Xiao, Shihui Zhang, Weizhong Zheng, Xiaomeng Wan, Zhuoxuan Li, Mingze Gao, Gefei Wang, Zhilin Chen, et al. Organoid-based single-cell spatiotemporal gene expression landscape of human embryonic development and hematopoiesis. bioRxiv, pages 2022–09, 2022.

[21] Jonathan Ho, Ajay Jain, and Pieter Abbeel. Denoising diffusion probabilistic models. Advances in Neural Information Processing Systems, 33:6840–6851, 2020.

[22] Yang Song and Stefano Ermon. Generative modeling by estimating gradients of the data distribution. Advances in Neural Information Processing Systems, 32, 2019.

[23] Nanxin Chen, Yu Zhang, Heiga Zen, Ron J Weiss, Mohammad Norouzi, and William Chan. Wavegrad: Estimating gradients for waveform generation. arXiv preprint arXiv:2009.00713, 2020.

[24] Max Welling and Yee W Teh. Bayesian learning via stochastic gradient langevin dynamics. In Proceedings of the 28th international conference on machine learning (ICML-11), pages 681–688. Citeseer, 2011.

[25] Meng Zhang, Stephen W Eichhorn, Brian Zingg, Zizhen Yao, Kaelan Cotter, Hongkui Zeng, Hongwei Dong, and Xiaowei Zhuang. Spatially resolved cell atlas of the mouse primary motor cortex by merfish. Nature, 598(7879):137–143, 2021.

[26] Xiao Wang, William E Allen, Matthew A Wright, Emily L Sylwestrak, Nikolay Samusik, Sam Vesuna, Kathryn Evans, Cindy Liu, Charu Ramakrishnan, Jia Liu, et al. Three-dimensional intact-tissue sequencing of single-cell transcriptional states. Science, 361(6400):eaat5691, 2018.

[27] Bosiljka Tasic, Zizhen Yao, Lucas T Graybuck, Kimberly A Smith, Thuc Nghi Nguyen, Darren Bertagnolli, Jeff Goldy, Emma Garren, Michael N Economo, Sarada Viswanathan, et al. Shared and distinct transcriptomic cell types across neocortical areas. Nature, 563(7729):72–78, 2018.

[28] Ron Zeira, Max Land, Alexander Strzalkowski, and Benjamin J Raphael. Alignment and integration of spatial transcriptomics data. Nature Methods, 19(5):567–575, 2022.

[29] Shiquan Sun, Jiaqiang Zhu, and Xiang Zhou. Statistical analysis of spatial expression patterns for spatially resolved transcriptomic studies. Nature methods, 17(2):193–200, 2020.

[30] Jiaqiang Zhu, Shiquan Sun, and Xiang Zhou. Spark-x: non-parametric modeling enables scalable and robust detection of spatial expression patterns for large spatial transcriptomic studies. Genome biology, 22(1):1–25, 2021.

[31] Nuha BinTayyash, Sokratia Georgaka, ST John, Sumon Ahmed, Alexis Boukouvalas, James Hensman, and Magnus Rattray. Non-parametric modelling of temporal and spatial counts data from rna-seq experiments. Bioinformatics, 37(21):3788–3795, 2021.

[32] Yingyao Zhou, Bin Zhou, Lars Pache, Max Chang, Alireza Hadj Khodabakhshi, Olga Tanaseichuk, Christopher Benner, and Sumit K Chanda. Metascape provides a biologist-oriented resource for the analysis of systems-level datasets. Nature communications, 10(1):1–10, 2019.

[33] Lin Mei and Klaus-Armin Nave. Neuregulin-erbb signaling in the nervous system and neuropsychiatric diseases. Neuron, 83(1):27–49, 2014.

[34] Weimin Luan, Xiqian Qi, Feng Liang, Xiaotao Zhang, Ziyang Jin, Ligen Shi, Benyan Luo, and Xuejiao Dai. Microglia impede oligodendrocyte generation in aged brain. Journal of Inflammation Research, 14:6813, 2021.

[35] Gregory C Johnson, Rodney Parsons, Victor May, and Sayamwong E Hammack. The role of pituitary adenylate cyclase-activating polypeptide (pacap) signaling in the hippocampal dentate gyrus. Frontiers in cellular neuroscience, 14:111, 2020.

[36] Katrin Gerstmann and Geraldine Zimmer. The role of the eph/ephrin family during cortical development and cerebral malformations. Medical Research Archives, 6(3), 2018.

[37] Monika Litviňuková, Carlos Talavera-López, Henrike Maatz, Daniel Reichart, Catherine L Worth, Eric L Lindberg, Masatoshi Kanda, Krzysztof Polanski, Matthias Heinig, Michael Lee, et al. Cells of the adult human heart. Nature, 588(7838):466–472, 2020.

[38] Genomics, 10x. 10x Gennomics Visium. Human Heart. https://www.10xgenomics.com/resources/datasets/human-heart-1-standard-1-1-0. Accessed: 2022-02-25.

[39] Michael Vanlandewijck, Liqun He, Maarja Andaloussi Mäe, Johanna Andrae, Koji Ando, Francesca Del Gaudio, Khayrun Nahar, Thibaud Lebouvier, Bàrbara Laviña, Leonor Gouveia, et al. A molecular atlas of cell types and zonation in the brain vasculature. Nature, 554(7693):475–480, 2018.

[40] Mark Sweeney and Gabor Foldes. It takes two: endothelial-perivascular cell cross-talk in vascular development and disease. Frontiers in Cardiovascular Medicine, 5:154, 2018.

[41] Lauren J Manderfield, Frances A High, Kurt A Engleka, Feiyan Liu, Li Li, Stacey Rentschler, and Jonathan A Epstein. Notch activation of jagged1 contributes to the assembly of the arterial wall. Circulation, 125(2):314–323, 2012.

[42] Celina Madjene, Alexandre Boutigny, Marie-Christine Bouton, Veronique Arocas, and Benjamin Richard. Protease nexin-1 in the cardiovascular system: Wherefore art thou? Frontiers in Cardiovascular Medicine, 8:652852, 2021.

[43] Marie-Christine Bouton, Yacine Boulaftali, Benjamin Richard, Véronique Arocas, Jean-Baptiste Michel, and Martine Jandrot-Perrus. Emerging role of serpine2/protease nexin-1 in hemostasis and vascular biology. Blood, The Journal of the American Society of Hematology, 119(11):2452–2457, 2012.

[44] Jie Wang, Yi Miao, Rebecca Wicklein, Zijun Sun, Jinzhao Wang, Kevin M Jude, Ricardo A Fernandes, Sean A Merrill, Marius Wernig, K Christopher Garcia, et al. Rtn4/nogo-receptor binding to bai adhesion-gpcrs regulates neuronal development. Cell, 184(24):5869–5885, 2021.

[45] Max Karlsson, Cheng Zhang, Loren Méar, Wen Zhong, Andreas Digre, Borbala Katona, Evelina Sjöstedt, Lynn Butler, Jacob Odeberg, Philip Dusart, et al. A single–cell type transcriptomics map of human tissues. Science Advances, 7(31):eabh2169, 2021.

[46] Sha Mi, Xinhua Lee, Zhaohui Shao, Greg Thill, Benxiu Ji, Jane Relton, Melissa Levesque, Norm Allaire, Steve Perrin, Bryan Sands, et al. Lingo-1 is a component of the nogo-66 receptor/p75 signaling complex. Nature neuroscience, 7(3):221–228, 2004.

[47] Kevin C Wang, Jieun A Kim, Rajeev Sivasankaran, Rosalind Segal, and Zhigang He. P75 interacts with the nogo receptor as a co-receptor for nogo, mag and omgp. Nature, 420(6911):74–78, 2002.

[48] Lisette Acevedo, Jun Yu, Hediye Erdjument-Bromage, Robert Qing Miao, Ji-Eun Kim, David Fulton, Paul Tempst, Stephen M Strittmatter, and William C Sessa. A new role for nogo as a regulator of vascular remodeling. Nature medicine, 10(4):382–388, 2004.

[49] Luyi Tian, Fei Chen, and Evan Z Macosko. The expanding vistas of spatial transcriptomics. Nature Biotechnology, pages 1–10, 2022.

[50] Francesca Prestori, Lisa Mapelli, and Egidio D’Angelo. Diverse neuron properties and complex network dynamics in the cerebellar cortical inhibitory circuit. Frontiers in Molecular Neuroscience, 12:267, 2019.

[51] Amanda M Brown, Marife Arancillo, Tao Lin, Daniel R Catt, Joy Zhou, Elizabeth P Lackey, Trace L Stay, Zhongyuan Zuo, Joshua J White, and Roy V Sillitoe. Molecular layer interneurons shape the spike activity of cerebellar purkinje cells. Scientific reports, 9(1):1–19, 2019.

[52] Beihui Liu, Valentina Mosienko, Barbara Vaccari Cardoso, Daria Prokudina, Mathew Huentelman, Anja G Teschemacher, and Sergey Kasparov. Glio-and neuro-protection by prosaposin is mediated by orphan g-protein coupled receptors gpr37l1 and gpr37. Glia, 66(11):2414–2426, 2018.

[53] Miho Taniguchi, Hiroaki Nabeka, Kimiko Yamamiya, Md Sakirul Islam Khan, Tetsuya Shimokawa, Farzana Islam, Takuya Doihara, Hiroyuki Wakisaka, Naoto Kobayashi, Fumihiko Hamada, et al. The expression of prosaposin and its receptors, grp37 and gpr37l1, are increased in the developing dorsal root ganglion. Plos one, 16(8):e0255958, 2021.

[54] Hyo Lee, Cheryl Pan, Sri Goberdhan, Jessica E Young, and Tracy Young-Pearse. Elucidating the role of sorl1 as an apoe receptor using ipsc-derived astrocytes: Molecular and cell biology/stem cells, ips cells. Alzheimer’s & Dementia, 16:e043860, 2020.

[55] Ed S Lein, Michael J Hawrylycz, Nancy Ao, Mikael Ayres, Amy Bensinger, Amy Bernard, Andrew F Boe, Mark S Boguski, Kevin S Brockway, Emi J Byrnes, et al. Genome-wide atlas of gene expression in the adult mouse brain. Nature, 445(7124):168–176, 2007.

[56] Dylan M Cable, Evan Murray, Vignesh Shanmugam, Simon Zhang, Luli S Zou, Michael Diao, Haiqi Chen, Evan Z Macosko, Rafael A Irizarry, and Fei Chen. Cell type-specific inference of differential expression in spatial transcriptomics. Nature methods, 19(9):1076–1087, 2022.

[57] Shaoqing Gong, Brenda Bin Su, Hugo Tovar, ChunXiang Mao, Valeria Gonzalez, Ying Liu, Yongke Lu, Ke-Sheng Wang, and Chun Xu. Polymorphisms within ryr3 gene are associated with risk and age at onset of hypertension, diabetes, and alzheimer’s disease. American Journal of Hypertension, 31(7):818–826, 2018.

[58] Paul W Sternberg, Julie Agapite, Laurent-Philippe Albou, Suzanne A Aleksander, Micheal Alexander, Anna V Anagnostopoulos, Giulia Antonazzo, Joanna Argasinska, Valerio Arnaboldi, Helen Attrill, et al. Harmonizing model organism data in the alliance of genome resources. Genetics, 220(4):iyac022, 2022.

[59] Yi-Wen Wu, KM Prakash, Tian-Yi Rong, Hui-Hua Li, Qin Xiao, Louis C Tan, Wing-Lok Au, Jian-qing Ding, Sheng-di Chen, and Eng-King Tan. Lingo2 variants associated with essential tremor and parkinson’s disease. Human genetics, 129(6):611–615, 2011.

[60] Min-Tzu Lo, Yunpeng Wang, Karolina Kauppi, Nilotpal Sanyal, Chun-Chieh Fan, Olav B Smeland, Andrew Schork, Dominic Holland, David A Hinds, Joyce Y Tung, et al. Modeling prior information of common genetic variants improves gene discovery for neuroticism. Human molecular genetics, 26(22):4530–4539, 2017.

[61] Robin Rombach, Andreas Blattmann, Dominik Lorenz, Patrick Esser, and Björn Ommer. High-resolution image synthesis with latent diffusion models. In Proceedings of the IEEE/CVF Conference on Computer Vision and Pattern Recognition, pages 10684–10695, 2022.

[62] Laleh Haghverdi, Maren Büttner, F Alexander Wolf, Florian Buettner, and Fabian J Theis. Diffusion pseudotime robustly reconstructs lineage branching. Nature methods, 13(10):845–848, 2016.

[63] F Alexander Wolf, Fiona K Hamey, Mireya Plass, Jordi Solana, Joakim S Dahlin, Berthold Göttgens, Nikolaus Rajewsky, Lukas Simon, and Fabian J Theis. Paga: graph abstraction reconciles clustering with trajectory inference through a topology preserving map of single cells. Genome biology, 20(1):1–9, 2019.

[64] Uwe Schmidt, Martin Weigert, Coleman Broaddus, and Gene Myers. Cell detection with star-convex polygons. In International Conference on Medical Image Computing and Computer-Assisted Intervention, pages 265–273. Springer, 2018.

[65] Carsen Stringer, Tim Wang, Michalis Michaelos, and Marius Pachitariu. Cellpose: a generalist algorithm for cellular segmentation. Nature methods, 18(1):100–106, 2021.

[66] Yi Yang, Xingjie Shi, Wei Liu, Qiuzhong Zhou, Mai Chan Lau, Jeffrey Chun Tatt Lim, Lei Sun, Cedric Chuan Young Ng, Joe Yeong, and Jin Liu. Sc-meb: spatial clustering with hidden markov random field using empirical bayes. Briefings in bioinformatics, 23(1):bbab466, 2022.

[67] Prafulla Dhariwal and Alexander Nichol. Diffusion models beat gans on image synthesis. Advances in Neural Information Processing Systems, 34:8780–8794, 2021.

[68] Yang Song and Stefano Ermon. Improved techniques for training score-based generative models. Advances in neural information processing systems, 33:12438–12448, 2020.

[69] Alexander Quinn Nichol and Prafulla Dhariwal. Improved denoising diffusion probabilistic models. In International Conference on Machine Learning, pages 8162–8171. PMLR, 2021.

[70] Samuel G Rodriques, Robert R Stickels, Aleksandrina Goeva, Carly A Martin, Evan Murray, Charles R Vanderburg, Joshua Welch, Linlin M Chen, Fei Chen, and Evan Z Macosko. Slide-seq: A scalable technology for measuring genome-wide expression at high spatial resolution. Science, 363(6434):1463–1467, 2019.

[71] Velina Kozareva, Caroline Martin, Tomas Osorno, Stephanie Rudolph, Chong Guo, Charles Vanderburg, Naeem Nadaf, Aviv Regev, Wade G Regehr, and Evan Macosko. A transcriptomic atlas of mouse cerebellar cortex comprehensively defines cell types. Nature, 598(7879):214–219, 2021.

