## Supplementary information for "SpatialScope: A unified approach for integrating spatial and single-cell transcriptomics data using deep generative models"

### 1 Supplementary Figures

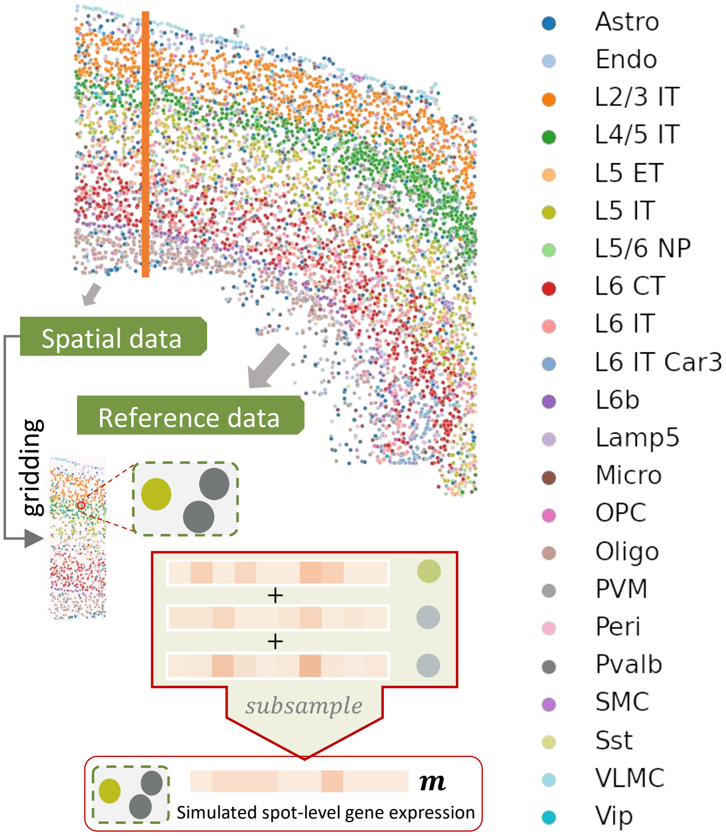

**Figure S1: The process of generating a simulated dataset using MERFISH dataset.** MERFISH cells are divided into two subsets. The left part is used to make pseudo-spots by aggregating the cells within each grid, and the right part is regarded as paired scRNA-seq reference. Uniform gridding is performed and cells in squares are aggregated to generate simulated spots.

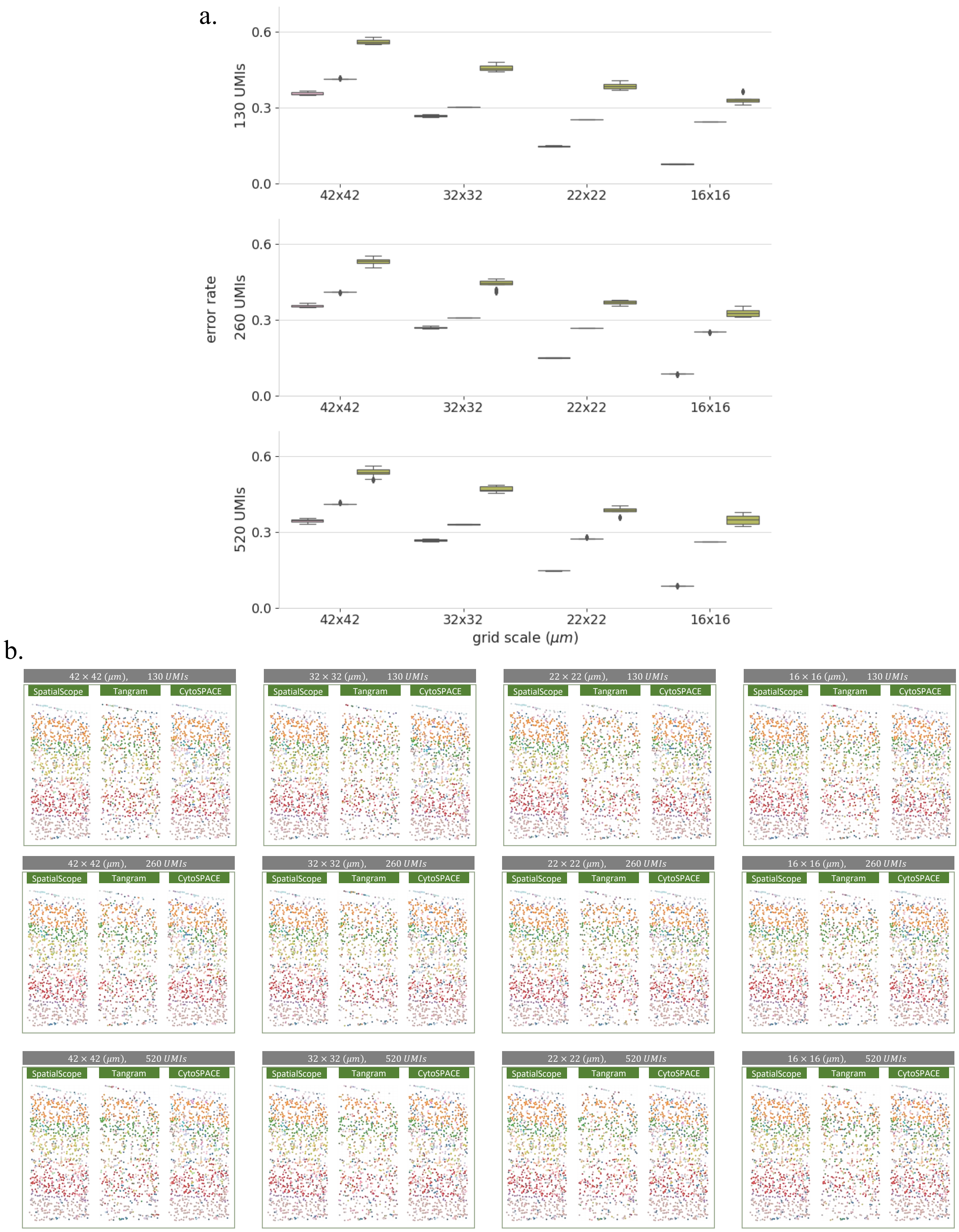

**Figure S2: Comparison of cell type identification performance under more simulation settings. a**, Summary of cell type identification results by error rates of the compared methods under different combination scenarios of grid scales and UMIs. **b**, The cell type identification results of SpatialScope, Tangram and CytoSPACE.

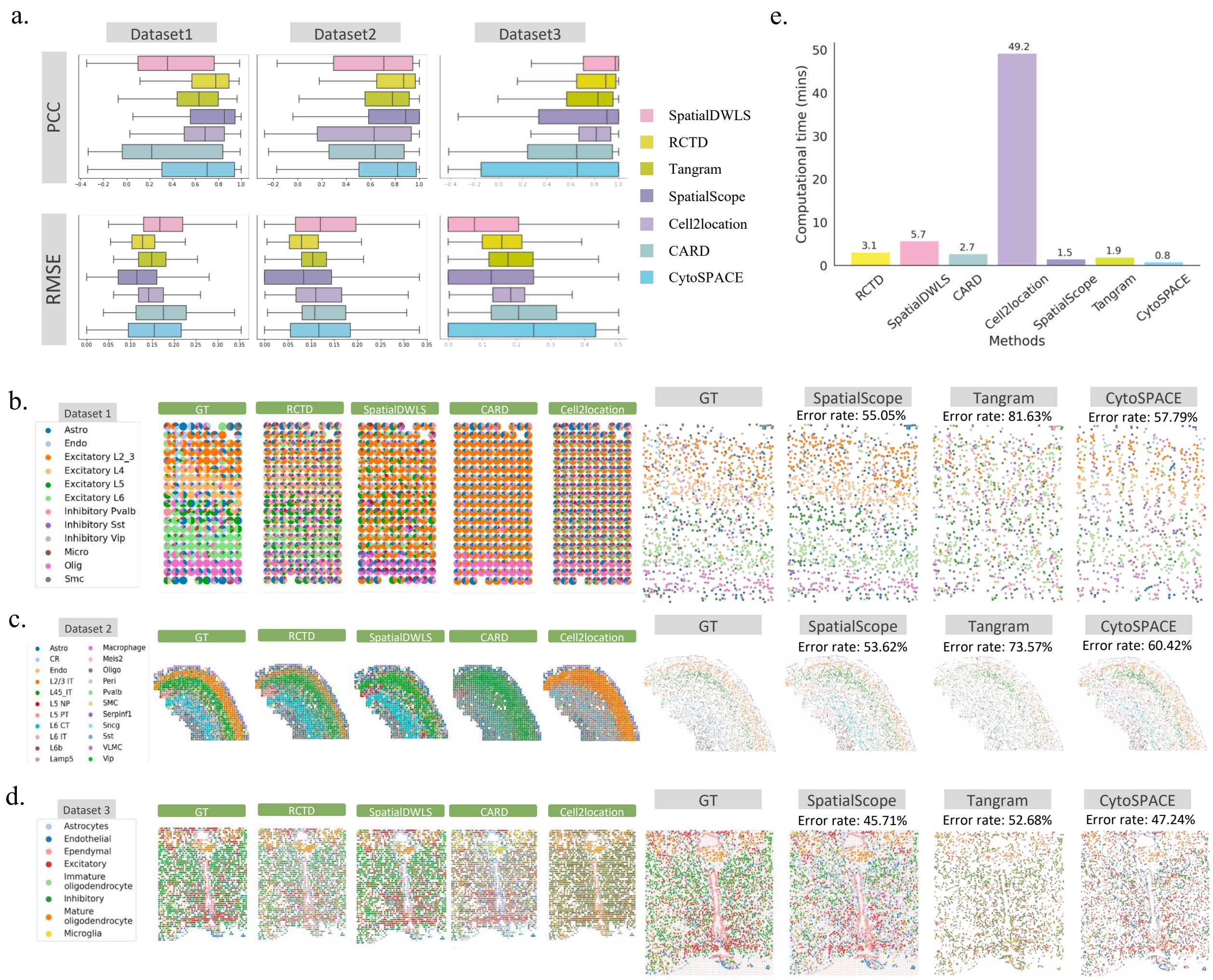

**Figure S3: Comparison of cell type identification/deconvolution under more simulation studies. a**, Summary of cell type deconvolution performance of the compared methods across four simulation datasets. Compared deconvolution methods include SpatialDWLS, RCTD, Tangram, Cell2location, CARD, CytoSPACE and SpatialScope. Top, Pearson correlation (PCC) and Bottom, root-mean-square error (RMSE), both of which quantify the similarity between the inferred cell-type composition from different methods (y-axis) and the ground truth, are displayed with box plots. **b-d** Left, a spatial scatter pie plot displays ground truth and inferred cell-type composition on each spatial location from different deconvolution methods; Right, A spatial scatter plot displays identified single cell types on each cell location from ground truth and different methods. **b** shows the results of the STARmap-based simulation dataset, and **c-d** shows the results of MERFISH-based simulation datasets. The detail simulation settings are given in supplementary note section 2.7.1. **e**, Computational times of SpatialScope and the compared methods when using MERFISH dataset as the simulated spot-level ST data.

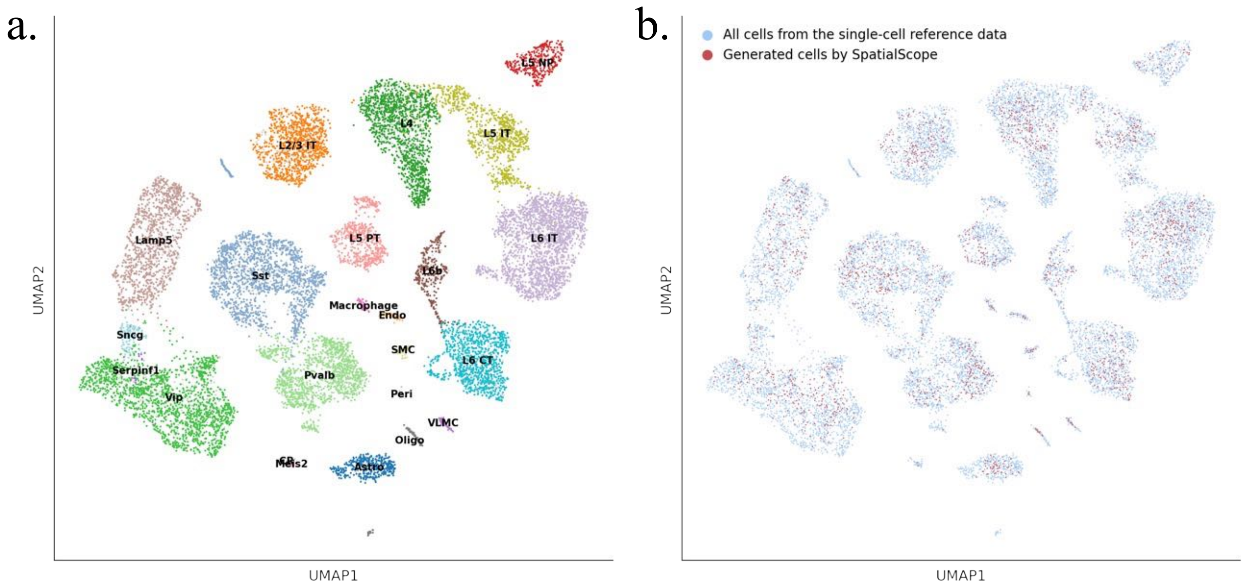

**Figure S4: scRNA-seq reference of mouse brain cortex a**, UMAP plot of scRNA-seq reference data with cell type annotations. **b**, UMAP plot of true cells and cells sampled from the learned distribution.

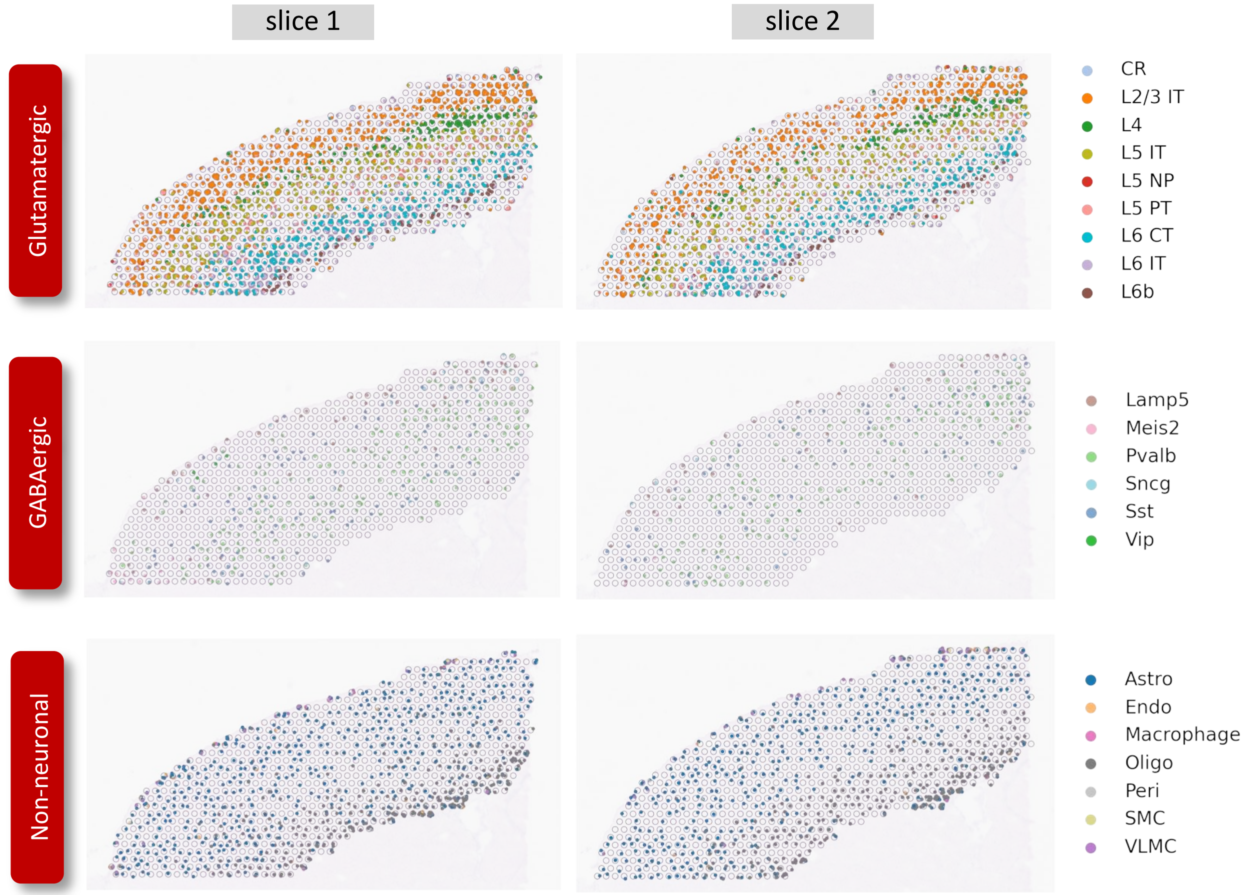

**Figure S5: Cell type identification results by SpatialScope for the stacked 3D Visium mouse brain cortex data.** Cell type identification results for slice 1 and slice2.

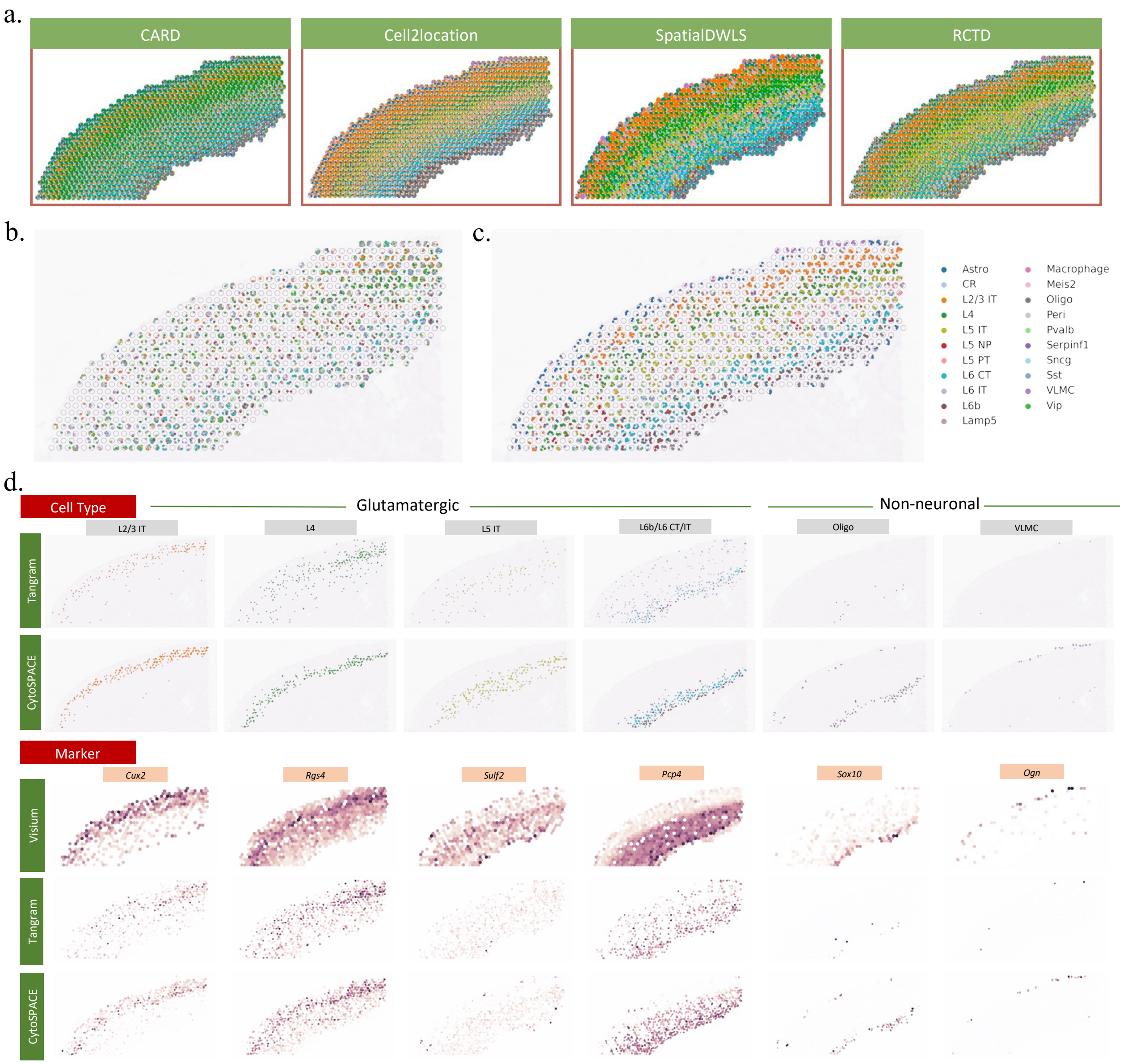

**Figure S6: Visium mouse brain cortex's cell type identification and gene expression decomposition results by the compared methods. a**, Cell type deconvolution results by spatialDWLS, RCTD, CARD, Cell2location. RCTD and Cell2location coarsely depicted the multi-layer structure, while CARD mistakenly assign many cells to the L4 layer and spatialDWLS did not provide clear boundaries across the layers. **b-c**, Cell type identification at single-cell resolution by Tangram and CytoSPACE. The canonical cortical four-layer structure is almost indistinguishable for Tangram due to missing cells caused by the soft regularized of cell number in each spot. CytoSPACE roughly reconstructed the four main layers but over-smoothed the cell type organization. For example, the right upper layer of the cortex was predicted to be Astrocytes only, while it should contain multiple cell types. **d**, The spatial cell type organizations (top) inferred by Tangram/CytoSPACE and Visium-measured spot-level (middle) and SpatialScope/CytoSPACE-decomposed single-cell level (bottom) expressions of a few marker genes.

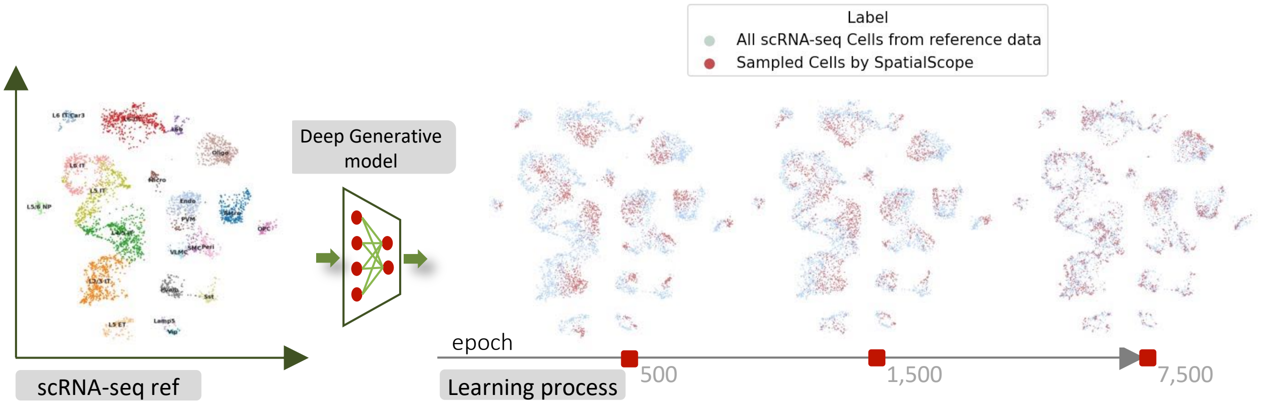

**Figure S7: The UMAP plots of single-cell reference data and learned distribution at 500 and 7500 epochs.** SpatialScope uses score-based generative modeling to learn the distribution of single-cell reference data.

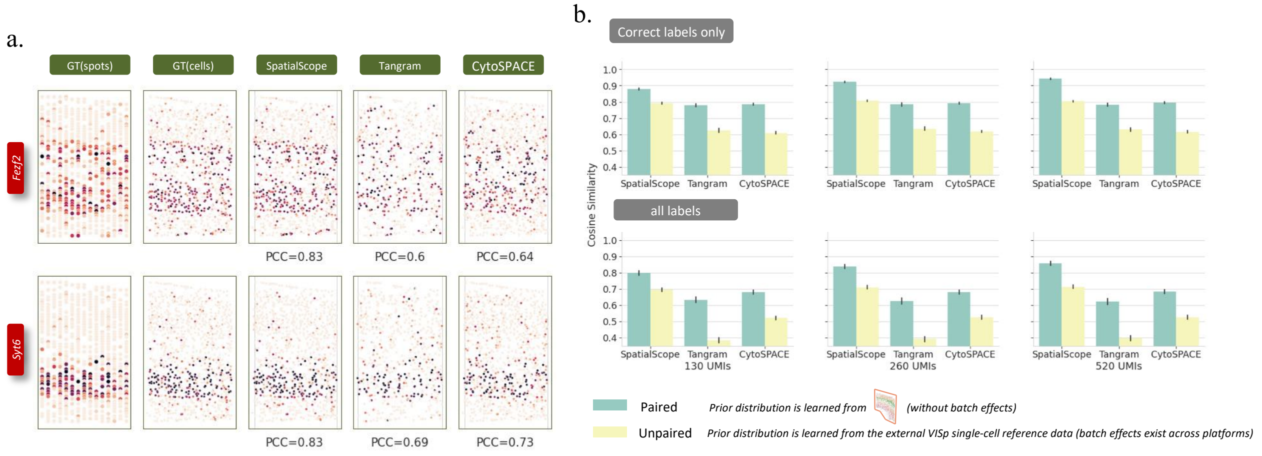

**Figure S8: Comparison of gene expression decomposition under more simulation studies. a**, Inspection of decomposed single-cell level expressions for a few layer marker genes. **b**, The cosine similarities between the ground truth and predicted gene expressions for cells with correctly identified cell type label (top) or all cells (bottom) under different combination scenarios of UMI subsample rate and single-cell reference data

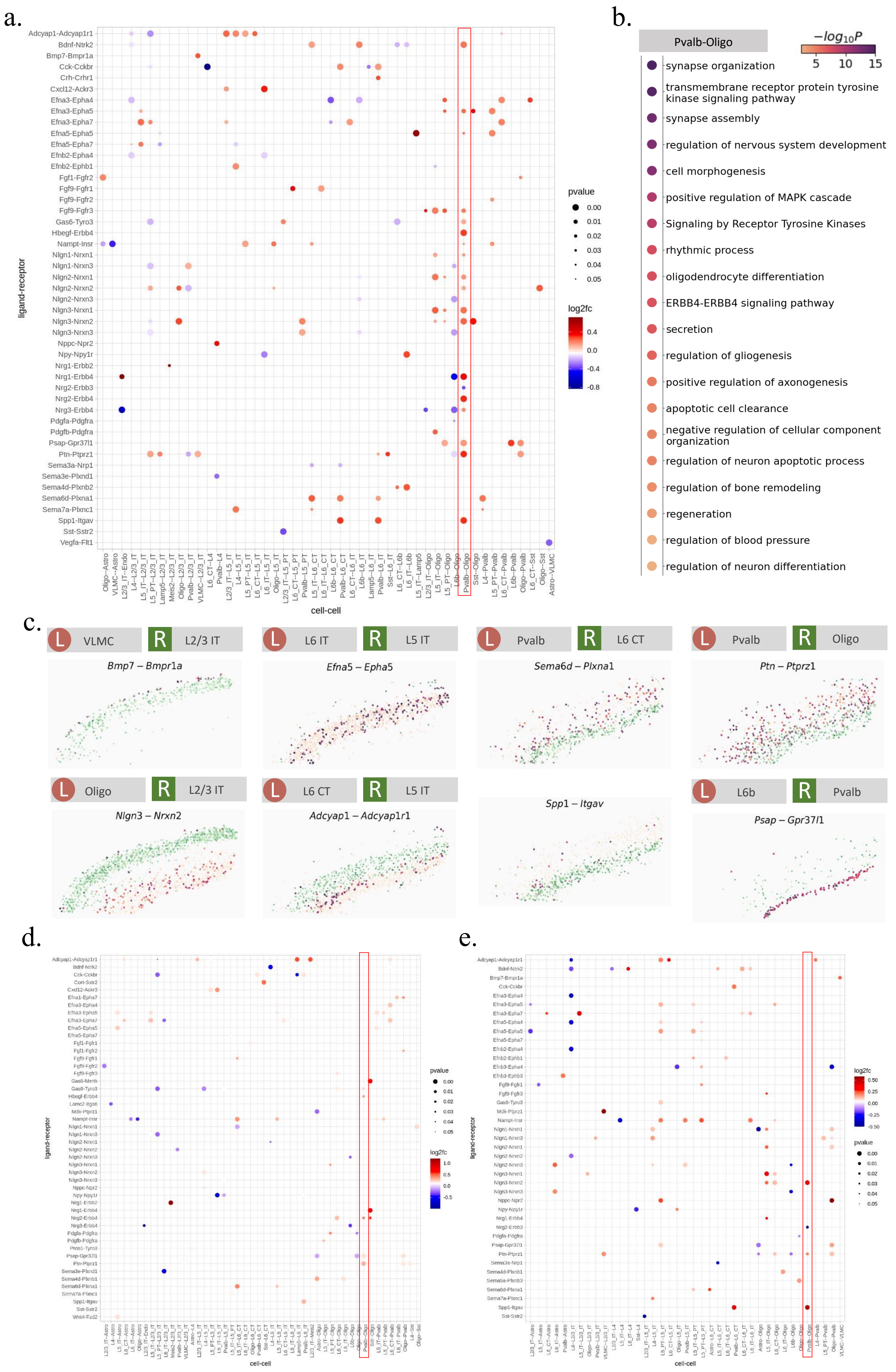
a.
b.
c.
d.
e.
ligand-receptor
cell-cell
pvalue
0.00
0.01
0.02
0.03
0.04
0.05
log2fc
0.4
0.0
-0.4
-0.8
Pvalb-Oligo
$-log_{10}P$
5
10
15
synapse organization
transmembrane receptor protein tyrosine kinase signaling pathway
synapse assembly
regulation of nervous system development
cell morphogenesis
positive regulation of MAPK cascade
Signaling by Receptor Tyrosine Kinases
rhythmic process
oligodendrocyte differentiation
ERBB4-ERBB4 signaling pathway
secretion
regulation of gliogenesis
positive regulation of axonogenesis
apoptotic cell clearance
negative regulation of cellular component organization
regulation of neuron apoptotic process
regulation of bone remodeling
regeneration
regulation of blood pressure
regulation of neuron differentiation
L VLMC
R L2/3 IT
Bmp7 – Bmpr1a
L L6 IT
R L5 IT
Efna5 – Epha5
L Pvalb
R L6 CT
Sema6d – Plxna1
L Pvalb
R Oligo
Ptn – Ptprz1
L Oligo
R L2/3 IT
Nlgn3 – Nrxn2
L L6 CT
R L5 IT
Adcyap1 – Adcyap1r1
Spp1 – Itgav
L L6b
R Pvalb
Psap – Gpr37l1

**Figure S9: Cell-cell interaction analysis in SpatialScope generated stacked 3D single-cell resolution ST mouse brain cortex data. a**, Dot plot of ligand-receptor pairs that exhibit spatially resolved cell-cell communications when analyzing the 3D aligned ST data. **b**, Significantly enriched Gene ontology biological processes determined with the Metascape web tool for the ligands and receptors from Pvalb and Oligo. **c** Visualization of a few representative cellular communications detected in the SpatialScope generated stacked 3D single-cell resolution ST mouse brain cortex data. **d**. Dot plot of ligand-receptor pairs that exhibit spatially resolved cell-cell communications when analyzing slice 1 only. **e**. Dot plot of ligand-receptor pairs that exhibit spatially resolved cell-cell communications when analyzing slice 2 only.

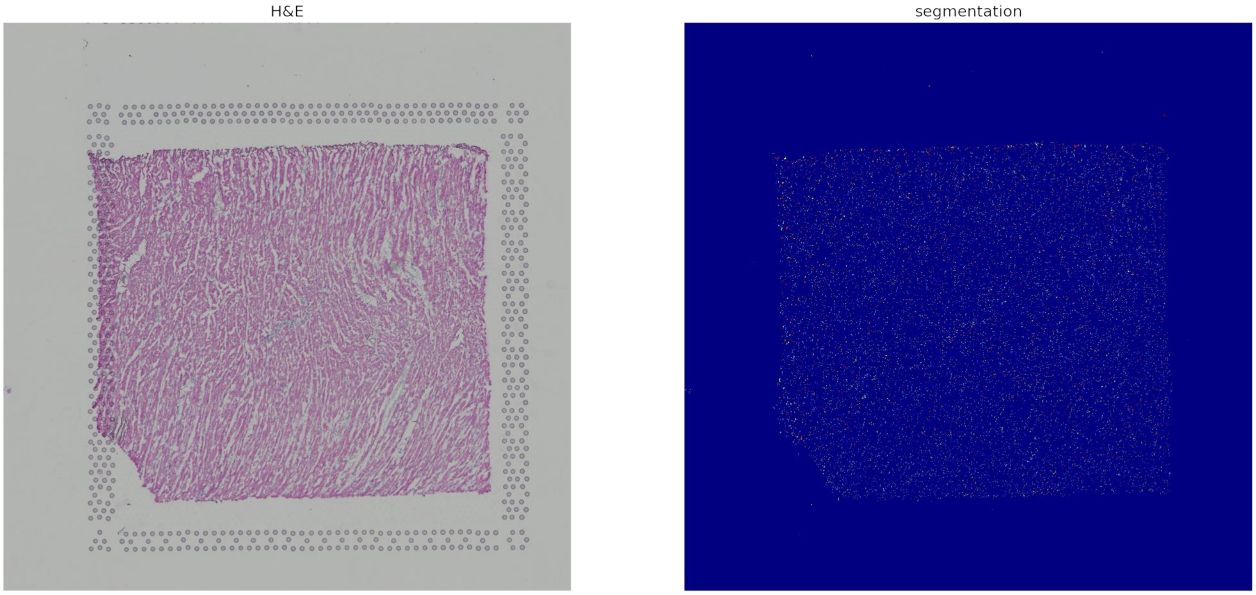

**Figure S10: Nuclei segmentation result for 10X Visium heart data. a**, The paired H&E stained histological image. **b**, The nuclei segmentation results

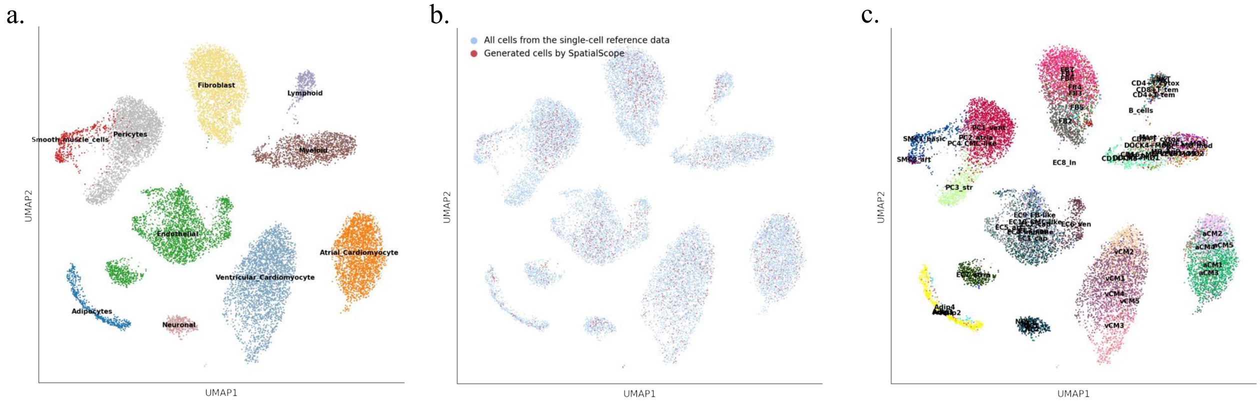

**Figure S11: snRNA-seq reference of human heart a**, UMAP plot of snRNA-seq reference data with cell type annotations. **b**, UMAP plot of true cells and cells sampled from the learned distribution, **c**, UMAP plot of snRNA-seq reference data with cell type subgroup annotations.

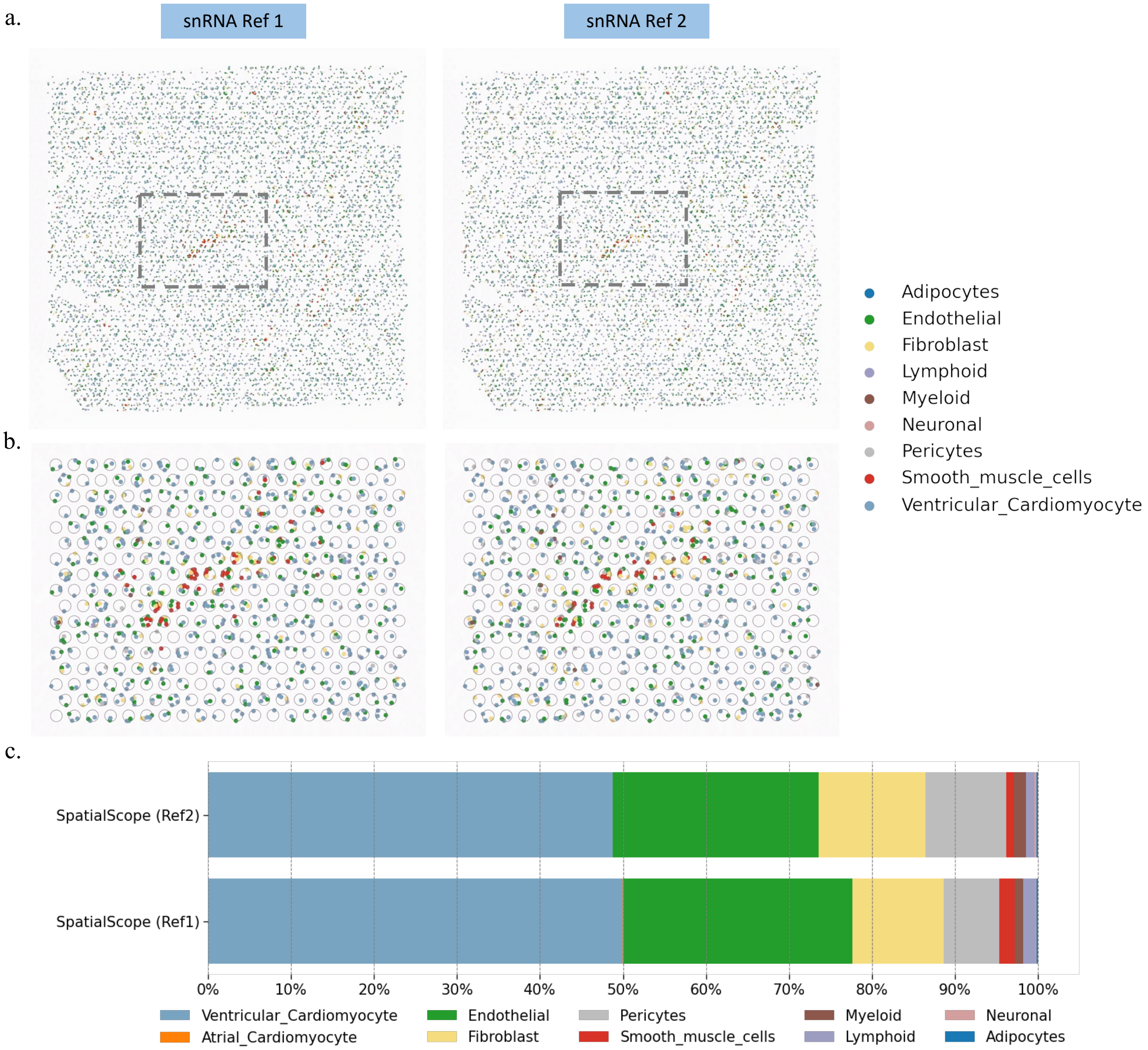

**Figure S12: Visium heart data cell type identification results by SpatialScope when distinct snRNA-seq reference was used.** **a**, Cell type identification at single-cell resolution for the whole slice. **b**, Cell type identification at single-cell resolution for ROI. **c**, Inferred cell type compositions across the whole slice when distinct snRNA-seq reference was used.

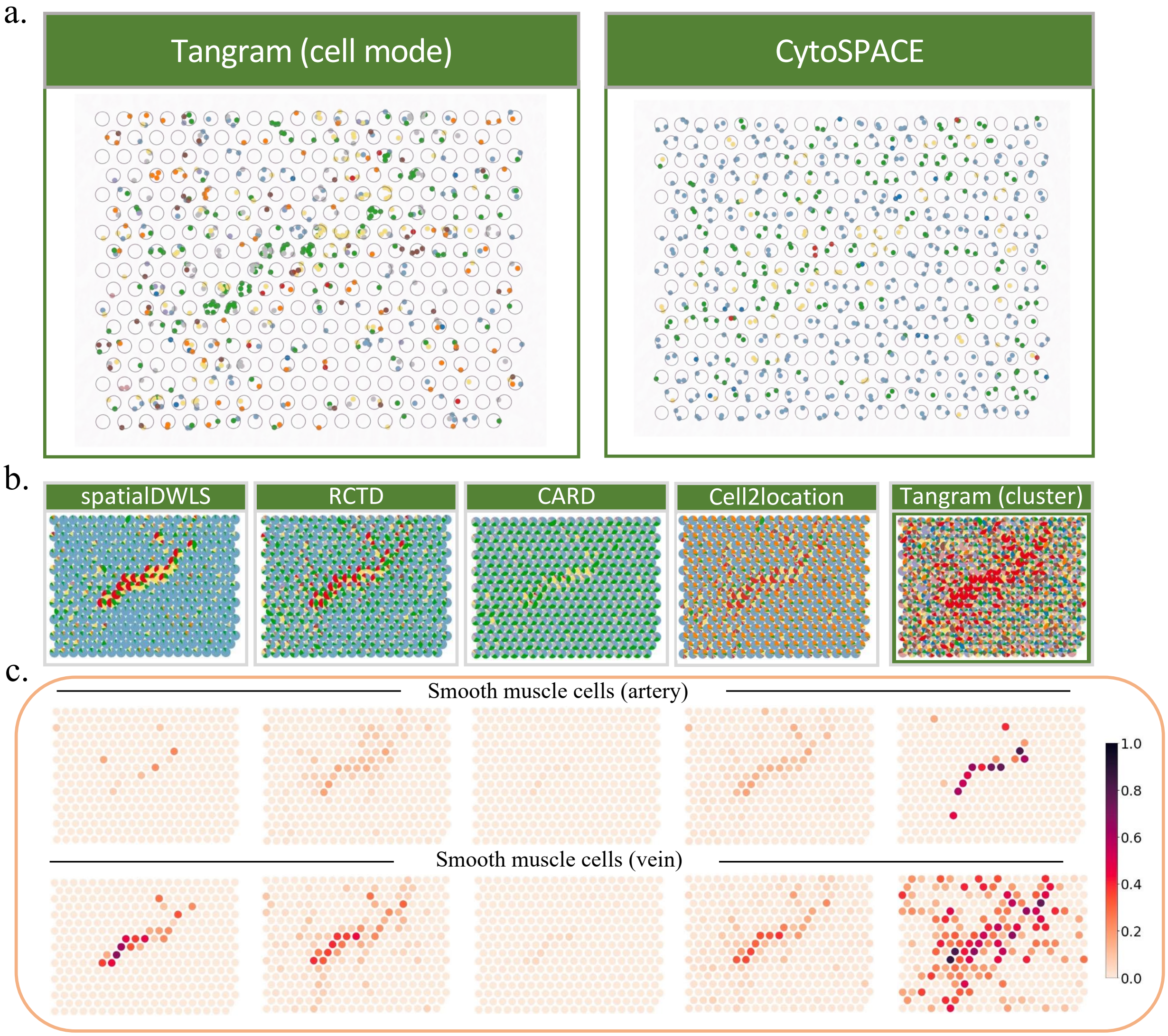

**Figure S13: Visium heart data cell type identification results by the compared methods a**, Cell type identification at single-cell resolution for ROI by Tangram (cell mode) and CytoSPACE. **b**, Cell type deconvolution results by spatialDWLS, RCTD, CARD, Cell2location and Tangram (cluster mode). **c**, Inferred cell type proportion for the two subgroups of SMC. We used the separated SMC cell type labels (SMC_artery, SMC_vein) in snRNA-seq reference rather than a unified SMC cell type label during the deconvolution, then we compared the estimated cell type proportion for these two subgroups to evaluate the discrimination abilities of the compared methods.

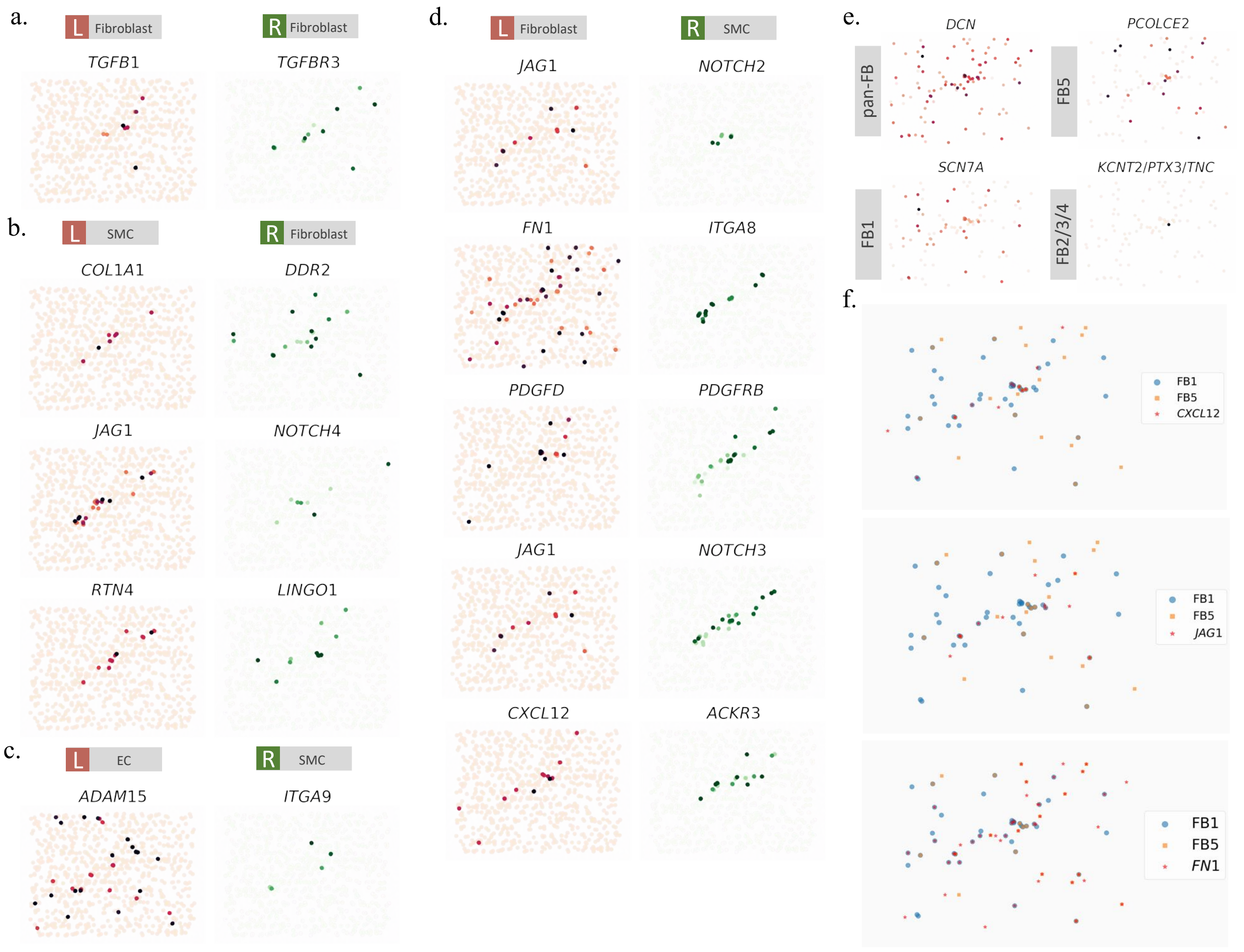

**Figure S14: More cellular communications detected in the SpatialScope generated single-cell resolution human heart ST data.**

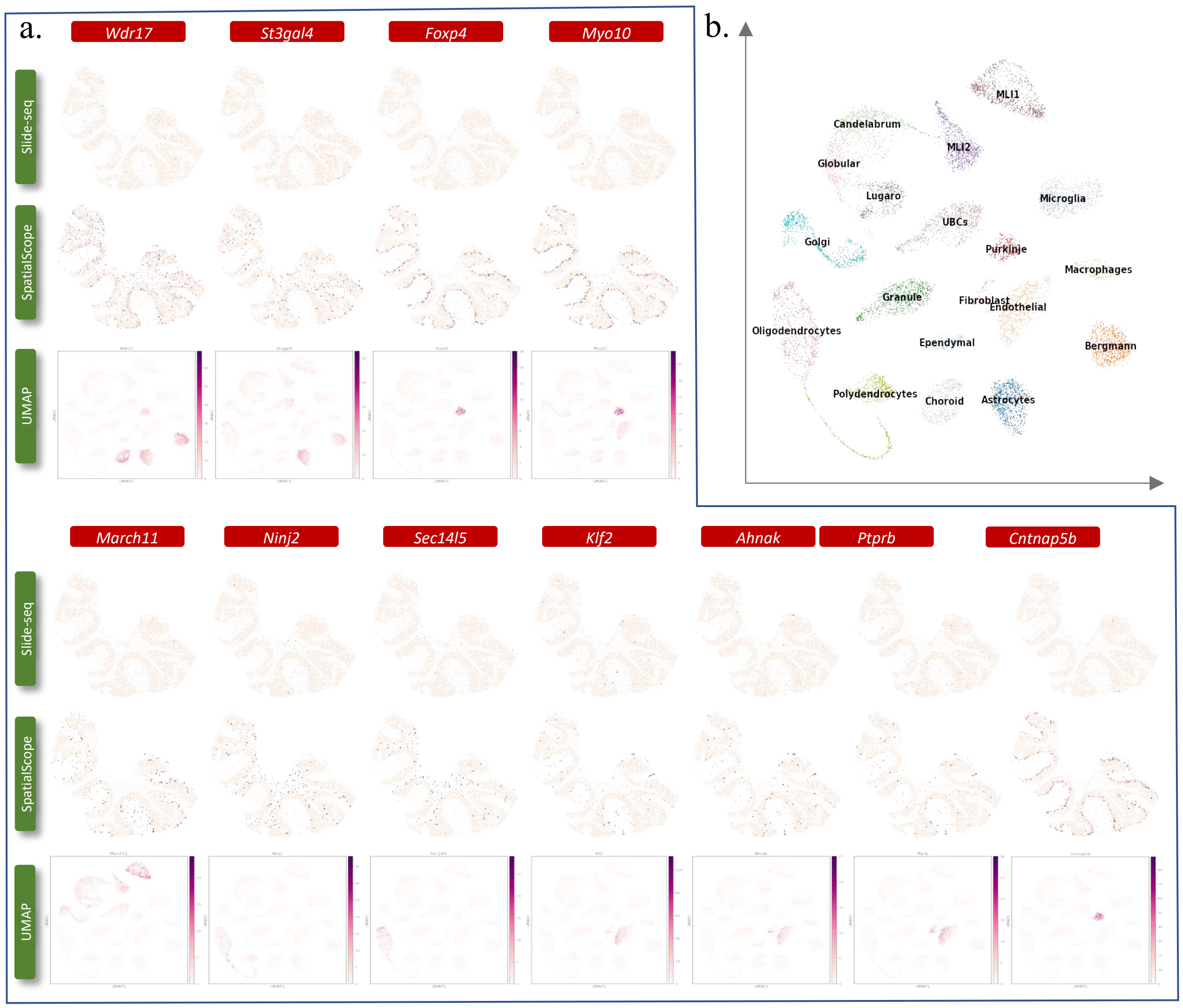

**Figure S15: Dropouts corrections by SpatialScope in Slide-seq data a**, Slide-seq measured (top) and SpatialScope corrected (middle) expressions of highly sparse marker genes. The marker gene expression signatures were displayed with UMAP plots. **b**, UMAP plot of snRNA-seq reference data with cell type annotations.

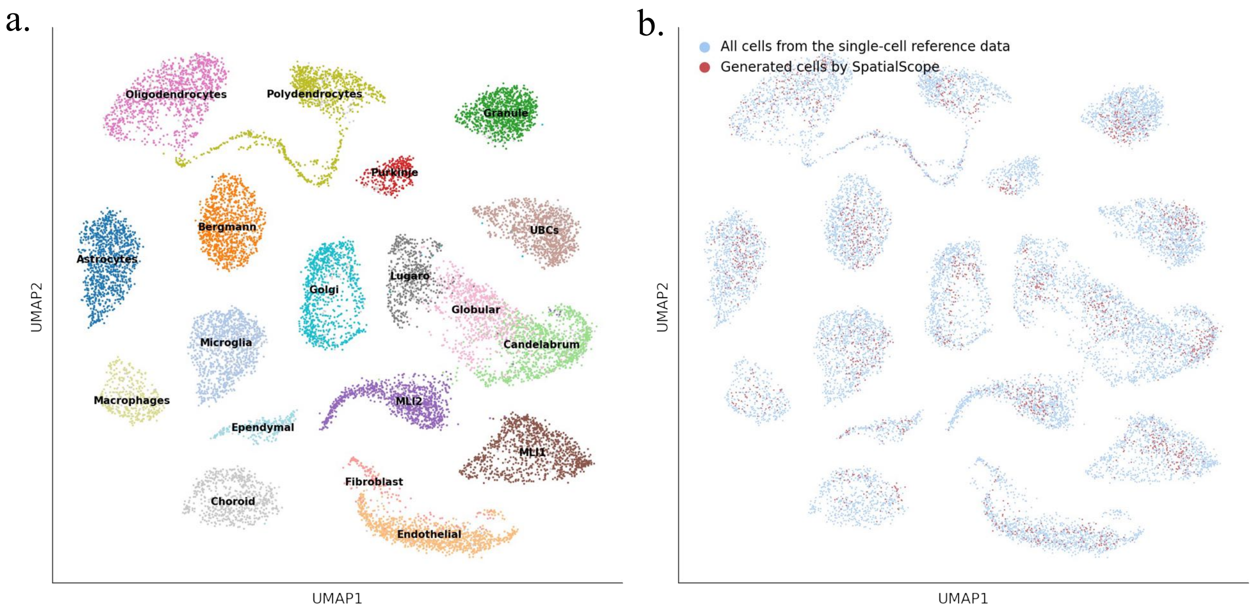

**Figure S16: snRNA-seq reference of mouse cerebellum a**, UMAP plot of snRNA-seq reference data with cell type annotations. **b**, UMAP plot of true cells and cells sampled from the learned distribution.

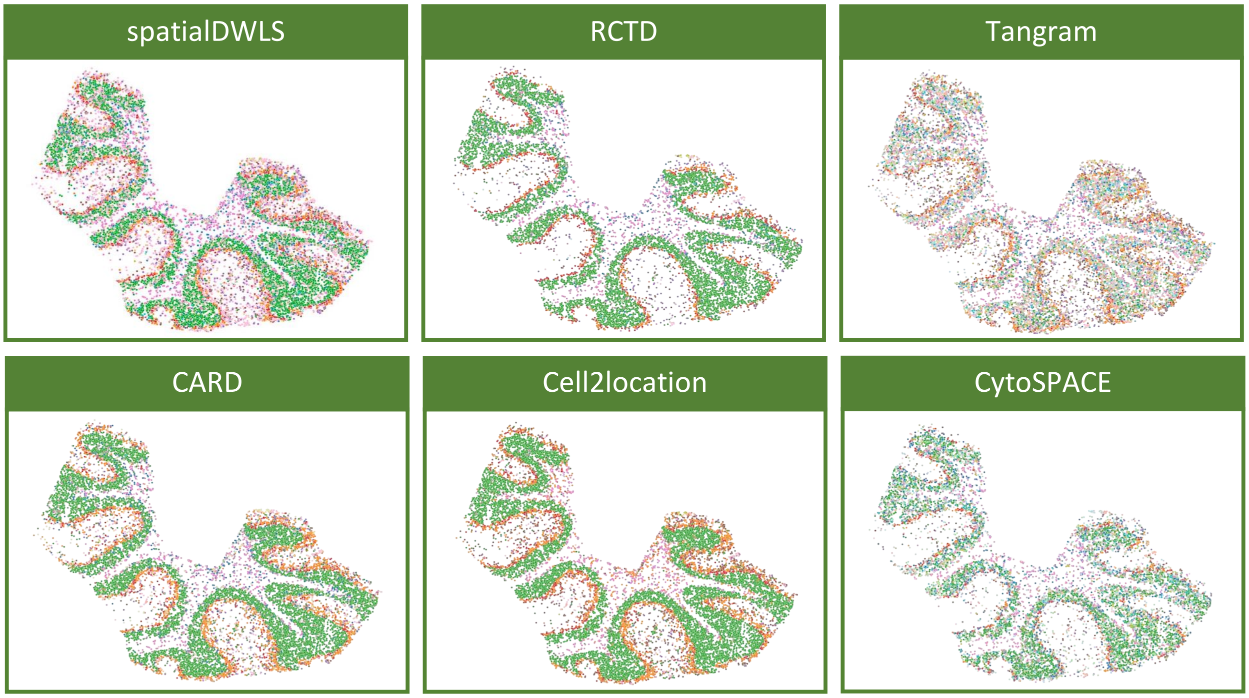

**Figure S17: Cell type identification results by the compared methods for Slide-seq V2 mouse cerebellum data.**

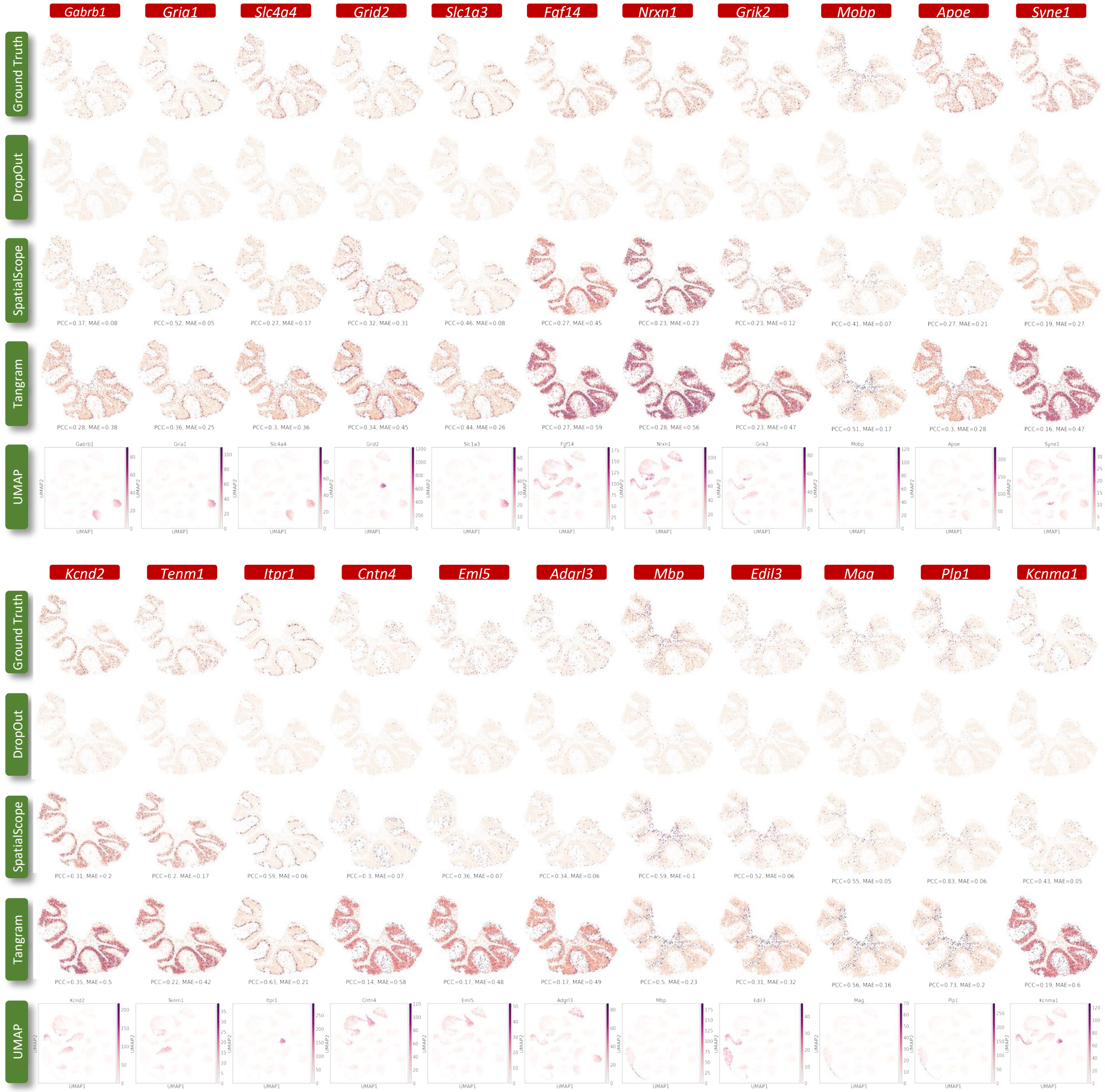

**Figure S18: Simulated dropouts and correction results by SpatialScope and Tangram based on the Slide-seq data.**

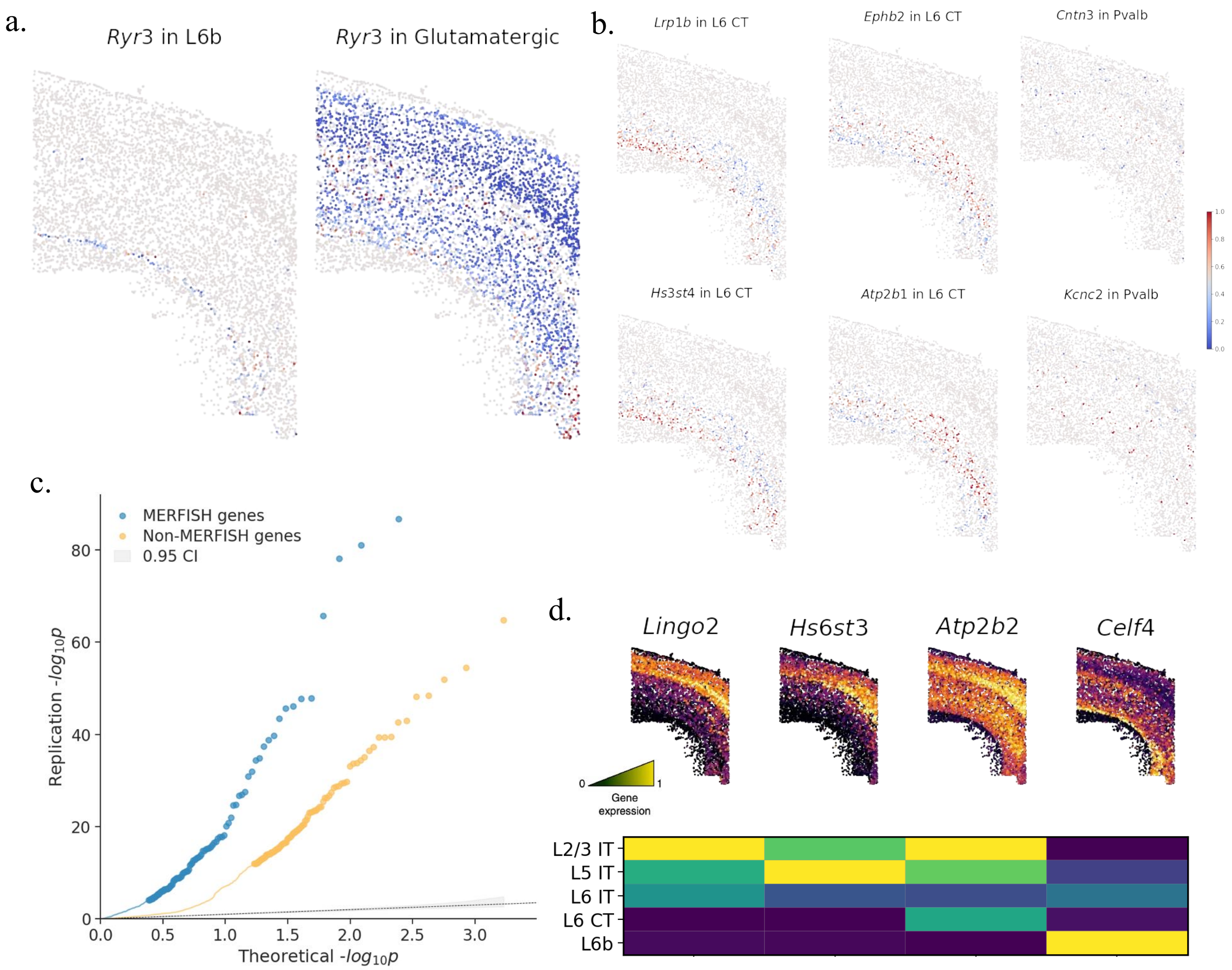

**Figure S19: Spatially DE genes detection results on MERFISH data.** **a**, The expression profile of *Ryr3* in L6b and Glutamatergic cell types, respectively. **b**, Representative examples of significant cell-type specific Non-MERFISH DE genes. **c**, QQ-plot of p-values for MERFISH and Non-MERFISH genes in the detection of spatially DE genes with SPARK-X. **d**, Visualization of a few representative Non-MERFISH spatially DE genes detected by SPARK-X, the gene expression signatures in snRNA-seq reference were displayed with heatmap plot.

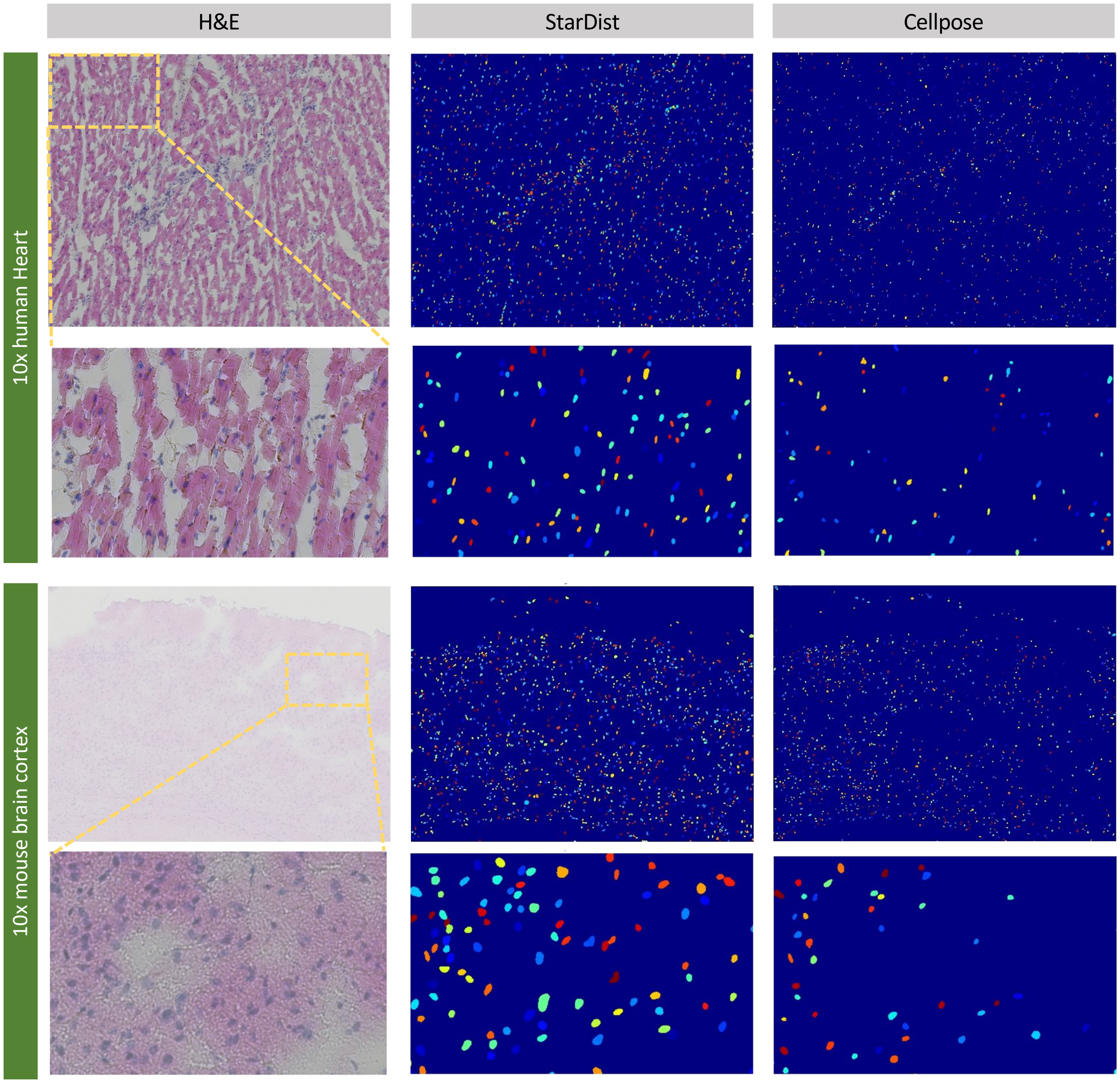

**Figure S20: Comparison of nucleus segmentation performance between StarDist and Cellpose in two 10x Visium datasets.** We applied Squidpy, which provides the interface of StarDist and Cellpose, to segment nuclei in the pair HE images. We used the default parameters following the instruction (`https://squidpy.readthedocs.io/en/stable/index.html`). In the first 10x human heart data, the H&E-stained histological image (first column) was used as input. The segmentation results of StarDist and Cellpose were shown in the second and last column, respectively, where StarDist located 1797 single cells and Cellpose only found 1301 cells. Clearly, Cellpose performed worse as a result of substantial missing cells, especially in the zoom-in region. For the second 10x mouse brain cortex dataset, we observed similar results that StarDist (n=1563) segments more cells than Cellpose (n=1250).

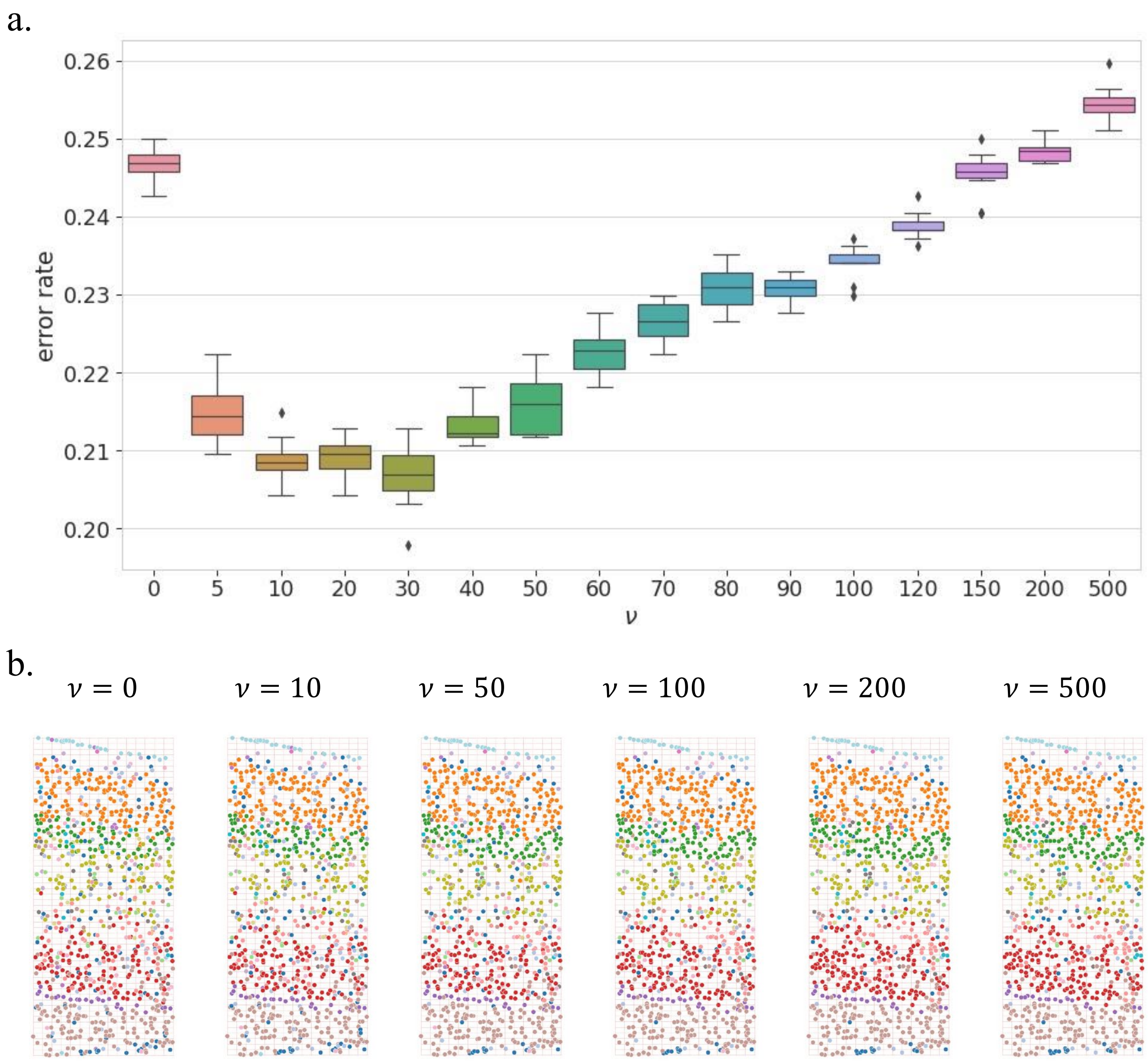

**Figure S21: Influence of hyperparameter $\nu$ in cell type identification task. a**. The cell type identification performance of SpatiaScope when different hyperparameter $\nu$ are used in the MERFISH benchmarking analysis. The performance is evaluated by error rate, which is defined as the proportion of mis-identified cell labels. **b**. Cell type identification results at $\nu = 0, 10, 50, 100, 200, 500$. Clearly, we observe that the smoothness of cell type distribution increases as the $\nu$ increases.

### 2 Supplementary Methods

#### 2.1 Cell type identification.

Suppose we have $K$ cell types in single-cell reference data. The expression counts of $G$ genes have been measured to capture the whole transcriptome in the scRNA-seq data. Let $k_{i,m} \in \{1, 2, \ldots, K\}$ be the cell type of the $m$-th cell at spot $i$, where $m = 1, \ldots, M_i$. Our goal is to infer the cell type vector $\mathbf{k}_i = \{k_{i,m}\}$ at spot $i$ by integrating scRNA-seq and ST data.

##### 2.1.1 Model setting

In this section, we revisit the probabilistic model for cell type identification in spatial transcriptomics (ST) data by incorporating scRNA-seq reference data. Inspired by RCTD [1], cell type means $\mu_{k,g}$ for cell type $k \in K$ and gene $g \in G$ are first estimated from annotated single-cell reference data:

$$\hat{\mu}_{k,g} \equiv \frac{1}{I_k} \sum_{n=1}^{I_k} \frac{x_{n,k,g}}{N_{n,k}}, \tag{S1}$$

where $I_k$ is the number of cells in reference of cell type $k$, $N_{n,k}$ is the number of UMIs of cell $n$ and cell type $k$ in single-cell reference data, and $x_{n,k,g}$ is observed counts of gene $g$ in this cell.

Next, we build a probabilistic model for each unique molecular identifier (UMI) in spots. For each UMI, the source cell is first probabilistically determined. Second, the gene that the UMI belongs to is determined based on the cell type of that cell. Formally, for each spot $1 \le i \le I$ and for each read $1 \le r \le N_i$, the probability that the read belongs to cell $1 \le \theta_{r,i} \le M_i$ and gene $1 \le z_{r,i} \le G$ are:

$$P\left(\theta_{r,i} = m | M_i, \mathbf{k}_i\right) = \frac{1}{M_i}, \quad P\left(z_{r,i} = g | \theta_{r,i}, \gamma, \varepsilon\right) \propto \delta_{i,\theta_{r,i},g}. \tag{S2}$$

Here, $\delta_{i,m,g}$ is defined as following

$$\log\left(\delta_{i,m,g}\right) = \alpha_i + \log\left(\hat{\mu}_{k_{i,m},g}\right) + \gamma_g + \varepsilon_{i,g}, \tag{S3}$$

where $\varepsilon_{i,g} \sim \mathcal{N}\left(0, \sigma_\varepsilon^2 \mathbf{I}\right)$ is a random effect to account for additional noise, and $\mu_{k,g}$ is defined as (S1). Both $\gamma_g$ and $\alpha_i$ are designed to address the batch effect between single-cell reference and ST data. More specifically, $\gamma_g \sim \mathcal{N}(0, \sigma_\gamma^2 \mathbf{I})$ represents a gene-specific random effect of accounting for expression differences of a gene $g$ between single-cell and ST platforms, and $\alpha_i$ is the spot-specific effect to account for differences of a gene set across platforms (See section 2.1.2 and section 2.1.3).

By combining formula (S2)(S3) and integrating $\theta$ out, the following holds,

$$\lambda_{i,g} \equiv P\left(z_{r,i} = g | \alpha_i, \gamma_g, \varepsilon_{i,g}, M_i, \mathbf{k}_i\right) \propto \frac{1}{M_i} \sum_{m=1}^{M_i} \hat{\mu}_{k_{i,m},g}. \tag{S4}$$

Consider $y_{i,g}$, the observed gene expression counts of gene $g$ at spot $i$ in ST data, as the sum of the reads that belong to gene $g$ in spot $i$, $y_{i,g} = \sum_{r=1}^{N_i} \mathbb{I}\left(z_{r,i} = g\right)$. Then we have

$$y_{i,1}, y_{i,2}, \ldots, y_{i,G} | \lambda_{i,1}, \lambda_{i,2}, \ldots, \lambda_{i,G} \sim \text{Multinomial}\left(N_i, \lambda_{i,1}, \lambda_{i,2}, \ldots, \lambda_{i,G}\right). \tag{S5}$$

Following RCTD, this distribution can be approximated by the Poisson distribution. Assuming that $N_i$ follows a Poisson distribution, $N_i \sim \text{Poisson}(\mu_i)$, and $y_{i,1}, y_{i,2}, \ldots, y_{i,G} \mid N_i \sim \text{Multinomial}\left(N_i, \lambda_{i,1}, \lambda_{i,2}, \ldots, \lambda_{i,G}\right)$. This induces $y_{i,g} \overset{\text{ind}}{\sim} \text{Poisson}\left(\mu_i \cdot \lambda_{i,g}\right)$. We can estimate $\hat{\mu}_i = N_i$ when $\mu_i$ is large. In practice, We filter spots with low UMIs and only consider spots with $N_i \ge 100$ to ensure that we are working in the regime of large $\mu_i$. Finally, we can model the counts $y_{i,g}$ as following,

$$\begin{aligned} y_{i,g} | \lambda_{i,g} &\overset{\text{ind}}{\sim} \text{Poisson}\left(N_i \lambda_{i,g}\right), \\ \log\left(\lambda_{i,\mathrm{g}}\right) &= \alpha_i + \log\left(\frac{1}{M_i} \sum_{m=1}^{M_i} \hat{\mu}_{k_{i,m},g}\right) + \gamma_{\mathrm{g}} + \varepsilon_{i,\mathrm{g}}. \end{aligned} \tag{S6}$$

##### 2.1.2 Spot-specific effect $\alpha_i$ accounting for platform differences

Parameter $\alpha_i$ is a free parameter that accounts for the difference in the total probability of observing a gene in the gene set $G = \{1, 2, \ldots, G\}$ between scRNA-seq and spot $i$. For example, highly expressed genes selected in scRNA-seq or snRNA-seq datasets may not be detected or less expressed in spot $i$. The spot index $i$ in $\alpha_i$ allows us to model this gene-set-level batch effect spot-by-spot, as the various cell type compositions across spatial spots may be associated with the gene-set-level batch effect.

##### 2.1.3 Gene-specific effect $\gamma_g$ accounting for platform difference

The platform batch effect characterized by the gene expression shift between scRNA-seq and spatial transcriptomics data limits the cell type knowledge transfer between the two RNA sequencing technologies. Therefore, following RCTD, we estimate and then correct the batch effect by summarizing the spatial transcriptomics data as a single pseudo-bulk measurement $S_g$:

$$S_g \equiv \sum_{i=1}^{I} y_{i,g} \sim \text{Poisson}\left(\sum_{i=1}^{I} N_i \lambda_{i,g}\right). \tag{S7}$$

Next we derive $N_i \lambda_{i,g}$. Plug (S6) into (S7), We have $\forall i, g$,

$$\begin{aligned}
\sum_{i=1}^{I} N_i \lambda_{i,g} &= \sum_{i=1}^{I} \sum_{m=1}^{M_i} \frac{1}{M_i} N_i \mu_{k_{i,m},g} e^{\gamma_g + \alpha_i + \varepsilon_{i,g}} = \sum_{i=1}^{I} \sum_{k=1}^{K} N_i \frac{M_i^k}{M_i} \mu_{k,g} e^{\gamma_g + \alpha_i + \varepsilon_{i,g}} \\
&= e^{\gamma_g} \sum_{k=1}^{K} \mu_{k,g} \sum_{i=1}^{I} N_i \frac{M_i^k}{M_i} e^{\alpha_i} e^{\varepsilon_{i,g}} \\
&= I e^{\gamma_g} \bar{N} \sum_{k=1}^{K} \mu_{k,g} B_{k,g},
\end{aligned} \tag{S8}$$

where $M_i^k$ is the number cells in spot $i$ that belongs to cell type $k$ and

$$\bar{N} = \frac{1}{I} \sum_{i=1}^{I} N_i, \quad B_{k,g} = \frac{1}{I} \sum_{i=1}^{I} \frac{N_i}{N} \frac{M_i^k}{M_i} \exp\left(\alpha_i + \varepsilon_{i,g}\right). \tag{S9}$$

Notice that $B_{k,g}$ only related to $g$ though $\varepsilon_{i,g}$. To get the point estimate of $\gamma_g$, we use the mean of $\varepsilon_{i,g}$ to replace $\varepsilon$ and define $W_k = \frac{1}{I} \sum_{i=1}^{I} \frac{N_i}{N} \frac{M_i^k}{M_i} e^{\alpha_i} e^{\sigma_\varepsilon^2/2}$ as a new unconstrained parameter representing the average bulk cell type proportion of cell type $k$. Then we have

$$S_g | \gamma_g \sim \text{ Poisson } \left(I \bar{N} e^{\gamma_g} \sum_{k=1}^{K} \mu_{k,g} W_k\right), \quad \gamma_g \sim \text{Normal}\left(0, \sigma_\gamma^2\right). \tag{S10}$$

When $I$ is large, the bulk Poisson mean is large for most genes. Therefore, we can approximate $S_g$ by it's mean:

$$\bar{S}_g \approx e^{\gamma_g} \sum_{k=1}^{K} \mu_{k,g} W_k \Longrightarrow \gamma_g | \hat{W} \approx \log\left(\bar{S}_g\right) - \log\left(\sum_{k=1}^{K} \mu_{k,g} \hat{W}_k\right) \equiv \hat{\gamma}_g, \tag{S11}$$

where $\bar{S}_g = S_g/(I\bar{N})$ and $W_k$ is estimated by solving the optimization problem $\hat{W}_k = \arg\min_{W_k} \frac{1}{2}\| \log\left(\bar{S}_g\right) - \sum_{k=1}^{K} \hat{\mu}_{k,g} W_k \|^2$.

##### 2.1.4 Cell type identification

After estimating platform effects, we treat the estimates $\hat{\gamma}_g$ and also $\hat{\mu}_{k,g}$ as fixed and then use SpatialScope to obtain the MAP estimate of cell type label $k_{i,m}$, where $i = 1, 2, \ldots, I$, and $m = 1, 2, \ldots, M_i$. Before the estimation, we first discuss the initialization of $\{k_{i,m}\}$. A warm start of RCTD will be applied to make the model more efficient. The major cell type of each spot estimated by RCTD will serve as the initial cell type for all cells in that spot. Besides, the platform effect $\alpha_i$ will also be estimated in the warm start, and we denote it as $\hat{\alpha}_i$. Next, we apply an iterative algorithm to identify cell type label $\{k_{i,m}\}$ and $\sigma_\varepsilon$. Recall that the MAP estimate for $\{k_{i,m}\}$ when given $\sigma_\varepsilon$ is as Eq. (6) in the main text method, where we update two labels $k_{i,m}, k_{i,\tilde{m}}$ at a time. The prior of $\{k_{i,m}\}$ is given as Eq. (4).

To make Eq. (6) in the main text feasible, we have the following derivation for $\log p\left(y_{i,g}|\hat{\boldsymbol{\theta}}_c, k_{im}, k_{i\tilde{m}}, k_{-\{(i,m),(i,\tilde{m})\}}\right)$ and $\log p\left(k_{im}, k_{i\tilde{m}}|k_{-\{(i,m),(i,\tilde{m})\}}\right)$. First, we derive $\log p\left(y_{i,g}|\hat{\boldsymbol{\theta}}_c, k_{im}, k_{i\tilde{m}}, k_{-\{(i,m),(i,\tilde{m})\}}\right)$. Define $w_{k,i} = \frac{1}{M_i} M_i^k e^{\hat{\alpha}_i}$, $\mathbf{w}_i = \{w_{k,i}\}_{k=1}^K$ and

$$\bar{\lambda}_{i,g}(\mathbf{w}_i) = \sum_{k=1}^{K} w_{k,i} \hat{\mu}_{k,g} e^{\hat{\gamma}_g} = \sum_{k=1}^{K} w_{k,i} \bar{\mu}_{k,g}. \tag{S12}$$

Based on (S6), we have following holds:

$$y_{i,g}|\bar{\lambda}_{i,g} \sim \text{Poisson}\left(e^{\varepsilon_{i,g}} N_i \bar{\lambda}_{i,g}(\mathbf{w}_i)\right), \quad \varepsilon_{i,g} \sim \text{Normal}\left(0, \hat{\sigma}_\varepsilon^2\right). \tag{S13}$$

By integrating out $\varepsilon_{i,j}$, we derive $\log p\left(y_{i,g}|\hat{\boldsymbol{\theta}}_c, k_{im}, k_{i\tilde{m}}, k_{-\{(i,m),(i,\tilde{m})\}}\right) = \log p\left(y_{i,g}|\bar{\lambda}_{i,g}\right)$ as following,

$$\begin{aligned} p\left(y_{i,g}|\bar{\lambda}_{i,g}\right) &= \int_{-\infty}^{\infty} p_\sigma(z) p\left(y_{i,j}|\lambda_{i,g} = \bar{\lambda}_{i,g} e^z\right) \mathrm{dz} \\ &= \int_{-\infty}^{\infty} p_\sigma(z) e^{-\bar{\lambda}_g N_i e^z} \frac{\left(N_i e^z \bar{\lambda}_g\right)^{y_{i,g}}}{y_{i,g}!} \mathrm{dz} \\ &= Q_{y_{i,g}}\left(\bar{\lambda}_{i,g}\right). \end{aligned} \tag{S14}$$

Here, $p_\sigma$ is the probability density function of $\varepsilon$. When $\hat{\sigma}_\varepsilon$ is obtained, $p_\sigma(z) = p_{\hat{\sigma}_\varepsilon}(z) = \frac{1}{\hat{\sigma}_\varepsilon\sqrt{2\pi}} e^{-\frac{1}{2}\left(\frac{z}{\hat{\sigma}_\varepsilon}\right)^2}$. The probability in (S14) is only related to the values of $y_{i,g}$ and $\bar{\lambda}_{i,g}$. To make the algorithm more efficient, a value table of the integration values in (S14) with respect to the values of $y_{i,g}$ and $\bar{\lambda}_{i,g}$ will be prepared in advance and $p\left(y_{i,g}|\bar{\lambda}_{i,g}\right)$ is calculated by searching the table in the algorithm.

Next, we derive the prior distribution $\log p\left(k_{im}, k_{i\tilde{m}}|k_{-\{(i,m),(i,\tilde{m})\}}\right)$. Recall that the prior of $\mathbf{K}$ is defined as Eq. (4) in the main text. To simplify the notation, omit subscript $i$ and use $k_m, k_{\tilde{m}}$ to denote the cell types of the cells $(i,m)$ and $(i,\tilde{m})$ respectively. We aim to calculate $p\left(k_{im}, k_{i\tilde{m}}|k_{-\{(i,m),(i,\tilde{m})\}}\right) = p(k_m, k_{\tilde{m}})$, where we omit $k_{-\{(i,m),(i,\tilde{m})\}}$ conditioning. To make the notation simple, we also omit $k_{-\{(i,m),(i,\tilde{m})\}}$ in the following derivation but keep in mind that all the probability in the following derivation of $p\left(k_{im}, k_{i\tilde{m}} | k_{-\{(i,m),(i,\tilde{m})\}}\right)$ are conditional on $k_{-\{(i,m),(i,\tilde{m})\}}$. Notice that $p(k_m, k_{\tilde{m}})$ can be rewrite as,

$$p(k_m, k_{\tilde{m}}) = p(k_{\tilde{m}}|k_m)p(k_m). \tag{S15}$$

To simplify the notation, denote $v_{m\tilde{m}} = p(k_m|k_{\tilde{m}}), v_{\tilde{m}m} = p(k_{\tilde{m}}|k_m)$. Since $p(k_{\tilde{m}})$ is a probability density function, it satisfies

$$\sum_j \frac{p(k_m)v_{\tilde{m}m}}{v_{m\tilde{m}}} = \sum_j p(k_{\tilde{m}}) = 1. \tag{S16}$$

Using this condition, we can rewrite $p(k_m)$ as $\frac{1}{\sum_j \frac{v_{\tilde{m}m}}{v_{m\tilde{m}}}}$. Plug $p(k_m) = \frac{1}{\sum_j \frac{v_{\tilde{m}m}}{v_{m\tilde{m}}}}$ into (S15), we have

$$p(k_m, k_{\tilde{m}}) = v_{\tilde{m}m} \cdot p(k_m) = v_{\tilde{m}m} \frac{1}{\sum_j \frac{v_{\tilde{m}m}}{v_{m\tilde{m}}}}. \tag{S17}$$

After taking logarithms on both sides of (S17):

$$\log p(k_m, k_{\tilde{m}}) = \log v_{\tilde{m}m} - \log \sum_j \frac{v_{\tilde{m}m}}{v_{m\tilde{m}}}. \tag{S18}$$

Because both $v_{\tilde{m}m}$ and $v_{m\tilde{m}}$ can be calculated by Eq. (4) in the main text, the prior term $\log p\left(k_{im}, k_{i\tilde{m}} | k_{-\{(i,m),(i,\tilde{m})\}}\right)$ in Eq. (6) in the main text now becomes feasible by using (S18).

Finally, plug (S14) and (S18) into Eq. (6) in the main text and then we can obtain MAP estimate of $k_{i,m}, k_{i,\tilde{m}}$ by maximizing the posterior distribution Eq. (6) in the main text . By finding the MAP estimate, we not only use information from gene expression levels $y_{i,g}$ to determine the cell type labels $k_{i,m}$, but also incorporate information from its neighbors.

***Algorithm*** We iteratively perform the following two steps: finding MAP estimate of $\{k_{i,m}\}$ and finding MLE for $\sigma_\varepsilon$. When finding MAP for $\{k_{i,m}\}$ given $\sigma_\varepsilon$, we maximize Eq. (6) in the main text one cell pair in a spot at a time and then iterate over all cell pairs. For each spot, the five cell types with the largest proportion are selected from the warm start results as candidate cell types for all cell pairs at that spot. When maximizing Eq. (6) in the main text, we search for all possible combinations of the two cell types among the five candidate cell types for computational efficiency. The combination that has the largest posterior Eq. (6) in the main text will be chosen as the MAP estimate for $k_{i,m}, k_{i,\tilde{m}}$.

Next, we compute MLE for $\sigma_\varepsilon$. When assume other parameters $\hat{\mu}_{k,g}, \hat{\gamma}_g, \hat{\alpha}_i, \hat{\mathbf{K}}$ are fixed, the log-likelihood of $\sigma_\varepsilon$ is given by (S14). The difference is that now we maximize (S14) with respect to $\sigma_\varepsilon$. Inspired by RCTD, we initialize $\sigma_\varepsilon = 1$. For each iteration, we randomly choose 500 spots, and then the MLE estimate of $\sigma_\varepsilon$ is calculated over these 500 spots. We maximize (S14) with respect to $\sigma_\varepsilon$ by searching 16 neighbors of previous $\sigma_\varepsilon$ value and the values of (S14) are obtained according to the pre-calculated table. The new $\sigma_\varepsilon$ value is chosen with the largest log-likelihood.

***"Smoothness" hyper-parameters*** In cell type identification, spatial information of spatial transcriptomic data is used to determine the labels by incorporating a smoothing prior, as shown in Eq. (4) in the main text. This is based on the intuition that cells that are close to each are more likely to have the same cell type. The "smoothness" hyper-parameters allow us to incorporate spatial information and make our model more robust over the noise. There are two hyper-parameters that are related to "smoothness". One is $\nu$ in Eq. (4) in the main text, and the other is $\mathcal{N}_{i,m}$, the neighbor for each cell. Both of them are tunable parameters. In practice, 1-norm distance is used to find neighbors. Ten neighbors are assigned to each cell in the default setting. Based on simulation studies, we set $\nu = 10$ to add moderate smoothing (Fig. S21). Users can decide to increase both $\mathcal{N}_{i,m}$ and $\nu$, which will result in a smoother spatial distribution of single-cell level cell types.

#### 2.2 Score-based generative models

We briefly review score-based generative models and then show how to leverage a conditional score-based generative models for SpatialScope in section 2.3. Let $\mathbf{x}$ be the log-scale single-cell expression level in single-cell reference data, following distribution $\mathbf{x} \sim p(\mathbf{x})$. The goal of score-based generative modeling is to obtain $p(\mathbf{x})$ by learning the score function: $\nabla_{\mathbf{x}} \log p(\mathbf{x})$ of probability density $p(\mathbf{x})$. Recall that multiple levels of Gaussian noise are added to the data. Let $\{\sigma_l\}_{l=1}^{L}$ be a sequence of positive noise level that satisfies $\sigma_L > \sigma_{L-1} > \cdots > \sigma_1 \approx 0$, and $\mathbf{x}^{(l)}$ be a sample perturbed by the noise level $\sigma_l^2$ with distribution $p_{\sigma_l}\left(\mathbf{x}^{(l)}\right) = \int p(\mathbf{x})\mathcal{N}\left(\mathbf{x}^{(l)}|\mathbf{x}, \sigma_l^2\mathbf{I}\right) d\mathbf{x}$. We aim to train a score network $s_\theta\left(\mathbf{x}^{(l)}, \sigma_l\right)$ to jointly learn all the *scores* of perturbed data distribution $\nabla_{\mathbf{x}^{(l)}} \log p_{\sigma_l}\left(\mathbf{x}^{(l)}\right), \forall l$. Formally, we consider the following objective function, which is called Explicit Score Matching (ESM) [2]:

$$\mathbb{E}_{p(\mathbf{x}^{(l)})}\left[\frac{1}{2}\left\|s_\theta\left(\mathbf{x}^{(l)}, \sigma_l\right) - \nabla_{\mathbf{x}^{(l)}} \log p_{\sigma_l}\left(\mathbf{x}^{(l)}\right)\right\|^2\right]. \tag{S19}$$

However, since $\nabla_{\mathbf{x}^{(l)}} \log p_{\sigma_l}\left(\mathbf{x}^{(l)}\right)$ cannot be computed, we consider the following denoising score matching (DSM) objective [2],

$$\ell(\boldsymbol{\theta}; \sigma_l) \triangleq \frac{1}{2}\mathbb{E}_{p(\mathbf{x})}\mathbb{E}_{\mathbf{x}^{(l)}\sim\mathcal{N}\left(\mathbf{x}, \sigma_l^2 I\right)}\left[\left\|\mathbf{s}_{\boldsymbol{\theta}}(\mathbf{x}^{(l)}, \sigma_l) - \nabla_{\mathbf{x}^{(l)}} \log p_{\sigma_l}\left(\mathbf{x}^{(l)}|\mathbf{x}\right)\right\|_2^2\right]. \tag{S20}$$

Note that

$$\nabla_{\mathbf{x}^{(l)}} \log p_{\sigma_l}(\mathbf{x}^{(l)}|\mathbf{x}) = -\frac{\mathbf{x}^{(l)} - \mathbf{x}}{\sigma_l^2}. \tag{S21}$$

After plugging (S21) into (S20), the denoising score matching objective (S20) is,

$$\ell(\boldsymbol{\theta}; \sigma_l) \triangleq \frac{1}{2}\mathbb{E}_{p(\mathbf{x})}\mathbb{E}_{\mathbf{x}^{(l)}\sim\mathcal{N}\left(\mathbf{x}^{(l)}, \sigma_l^2 I\right)}\left[\left\|\mathbf{s}_{\boldsymbol{\theta}}(\mathbf{x}^{(l)}, \sigma_l) + \frac{\mathbf{x}^{(l)} - \mathbf{x}}{\sigma_l^2}\right\|_2^2\right]. \tag{S22}$$

From the above, we can clearly see the intuition of (S20). The denoising process is to recover clean data from the data that is corrupted by the noise. The direction in (S21) is exactly from the noisy data to the clean data. Also, this is what we learned in the neural network (S22).

Then we combine (S22) for all noise level $\{\sigma_l\}_{l=1}^{L}$ to get the final objective,

$$\mathcal{L}\left(\boldsymbol{\theta}; \{\sigma_l\}_{l=1}^{L}\right) \triangleq \frac{1}{L}\sum_{l=1}^{L} \lambda\left(\sigma_l\right) \ell\left(\boldsymbol{\theta}; \sigma_l\right). \tag{S23}$$

Because we learn all the scores at the same time, we need to add the coefficient $\lambda(\sigma_l)$ to balance each term and make sure that we learn all the scores successfully. Notice that $\left|\frac{\mathbf{x}^{(l)}-\mathbf{x}}{\sigma_l^2}\right| \propto \frac{1}{\sigma_l}$. To make sure that the outputs of the score network have the same scale for different noise levels, we choose $\lambda_l(\sigma_l)$ to be $\sigma_l^2$.

Then we run annealed Langevin dynamics [3] (see Algorithm 1) to generate new samples from $p(\mathbf{x})$. First, we initialize $\mathbf{x}^{(0)}$ randomly and apply Langevin dynamics with the score network estimated at the largest noise level: $\mathbf{s}_{\boldsymbol{\theta}}(\mathbf{x}, \sigma_L) \approx \nabla_{\mathbf{x}^{(L)}} \log p_{\sigma_L}\left(\mathbf{x}^{(L)}\right)$. Then gradually annealed down the noise level from $l = L$ to $l = 1$ with initialization $\mathbf{x}^{(l,t=1)} = \mathbf{x}^{(l+1,t=T)}$. At the same time, the step size $\eta$ is also reducing:

$$\mathbf{x}^{(l,t+1)} = \mathbf{x}^{(l,t)} + \eta \mathbf{s}_{\boldsymbol{\theta}}\left(\mathbf{x}^{(l,t)}, \sigma_l\right) + \sqrt{2\eta}\boldsymbol{\varepsilon}^{(l,t)}. \tag{S24}$$

Finally, with the noise level and the step size becoming smaller and smaller, we obtain samples from $p_{\sigma_1}(\mathbf{x})$ which is close to the real clean data distribution $p(\mathbf{x})$ when $\sigma_1 \approx 0$.

**Algorithm 1** Annealed Langevin dynamics

**Require:** $\{\sigma_l\}_{l=1}^{L}, \eta_0, T$
  Initialize $\mathbf{x}^{(0)}$
  **for** $l = L, L-1, \ldots, 1$ **do**
    $\eta = \eta_0 \cdot \sigma_l^2/\sigma_1^2$
    **for** t = 1,2,...,T **do**
      Draw $\boldsymbol{\varepsilon}^{(l,t)} \sim \mathcal{N}(\mathbf{0}, \mathbf{I})$,

$$\mathbf{x}^{(l,t+1)} = \mathbf{x}^{(l,t)} + \eta \mathbf{s}_{\boldsymbol{\theta}}(\mathbf{x}^{(l,t)}, \sigma_l) + \sqrt{2\eta}\boldsymbol{\varepsilon}^{(l,t)}. \tag{S25}$$

    **end for**
    $\mathbf{x}^{(0)} = \mathbf{x}^{(T)}$
  **end for**

#### 2.3 SpatialScope: a conditional score-based generative model for single-cell reference data

Conditional generative models have been long time studied in the generative model field [4, 5, 6, 7, 8]. Many works choose to incorporate the conditioning information into the network for generative models. When the conditional information is discrete variables, one of the challenges is how to match the dimension of the discrete variable and some middle layer of the network, and at the same time, the conditioning information is well incorporated. Discrete conditioning information is always embedded in high dimensional space first, then use strategies like concatenating. For example, the embedding methods include a learnable map [7], or sinusoidal positional encoding [9]. Here, to encode the cell type conditioning information, we propose to learn the score function $\mathbf{s}_{\boldsymbol{\theta}}\left(\mathbf{x}^{(l)}, \sigma_l, \boldsymbol{\mu}_k\right)$ which takes the mean expression level of cell type $k$ as input. The benefits are two-fold. First, $\boldsymbol{\mu}_k$ provides precise information about cell type $k$. Second, $\boldsymbol{\mu}_k \in \mathbb{R}^G$ has the same dimension of $\mathbf{x}^{(l)}$ such that it will not be ignored. With this key idea, we can design a novel network architecture to learn the score function

$\mathbf{s}_{\boldsymbol{\theta}}\left(\mathbf{x}^{(l)}, \sigma_l, \boldsymbol{\mu}_k\right)$ (See network architecture for details). Besides, as empirically noted in [10], $\|\mathbf{s}_{\boldsymbol{\theta}}(\mathbf{x}, \sigma, \boldsymbol{\mu}_k)\| \propto 1/\sigma$ for trained score fuction on real data. We also find that incorporating the noise information by rescaling the score function is more stable and more widely applicable. Therefore, we use noise unconditional score network [11] and inject the information of the noise level by scaling, *i.e.* $\mathbf{s}_{\boldsymbol{\theta}}(\mathbf{x}, \sigma, \boldsymbol{\mu}_k) \approx \mathbf{s}_{\boldsymbol{\theta}}(\mathbf{x}, \boldsymbol{\mu}_k)/\sigma$.

#### 2.4 Network Architectures

The UNet architecture [12] is widely used in image diffusion models and has shown outstanding performance [13, 14]. Here we also use UNet architecture for our conditional score function and found it works well for learning gene expression distribution of single-cell reference data. There are two main differences between our architecture and image diffusion models. First, image data is two dimensional while single-cell data are one dimensional and therefore dilated 1 dimensional convolution is used as the smallest block in our network. Second, as we said before that we use a conditional score function where we take the cell type means $\boldsymbol{\mu}_k, k = 1, 2, \ldots, K$ as the actual input for score function instead of some embedding of cell type $k$. More specifically, we add another UNet (Fig. S22) taking $\boldsymbol{\mu}_k$ as input and in the middle of which produces both scale and bias vectors for feature-wise affine transformation applying on feature vector at some middle layer of main UNet that taking $\mathbf{x}$ as input.

The network architecture is shown in Fig. S22. There are two UNets taking $\mathbf{x}$ and $\boldsymbol{\mu}_k$ as input, respectively. In each UNet, three main blocks (MBlock) or conditional blocks (CBlock) are applied to gradually downsample the gene dimension by factors $3, 4, 5$ with the number of channels of $128, 256$, $512$, respectively. Then another three MBlocks or CBlocks, which are symmetric to the downsampling process, are applied to gradually upsample the gene dimension. The downsampling process and upsampling process are connected by additional MBlocks or CBlocks without downsampling or upsampling and yield a U-shaped architecture. The MBlock is illustrated in Fig. S23a. Each MBlock includes two residual blocks and four convolutional layers with dilation factors $1, 2, 1, 2$. Each CBlock (Fig. S23b) only includes one residual blocks and the dilation factors of three convolutional layers inside are $1, 2, 4$. Inspired by [15], the feature-wise linear modulation (FiLM) (Fig. S22, S23c) module is applied to produce feature-wise affine parameters, scale $\mathbf{W}$ and bias $\mathbf{b}$ vectors, to add cell type information in score function. Formally,

$$\mathbf{x}_{mid} = \mathbf{W}(\boldsymbol{\mu}_k) \odot \mathbf{x}_{mid} + \mathbf{b}(\boldsymbol{\mu}_k), \tag{S26}$$

where $\mathbf{W}$ and $\mathbf{b}$ correspond to the scaling and shift vectors produced by the FiLM module, $\mathbf{x}_{mid}$ is the corresponding middle layer output from MBlock.

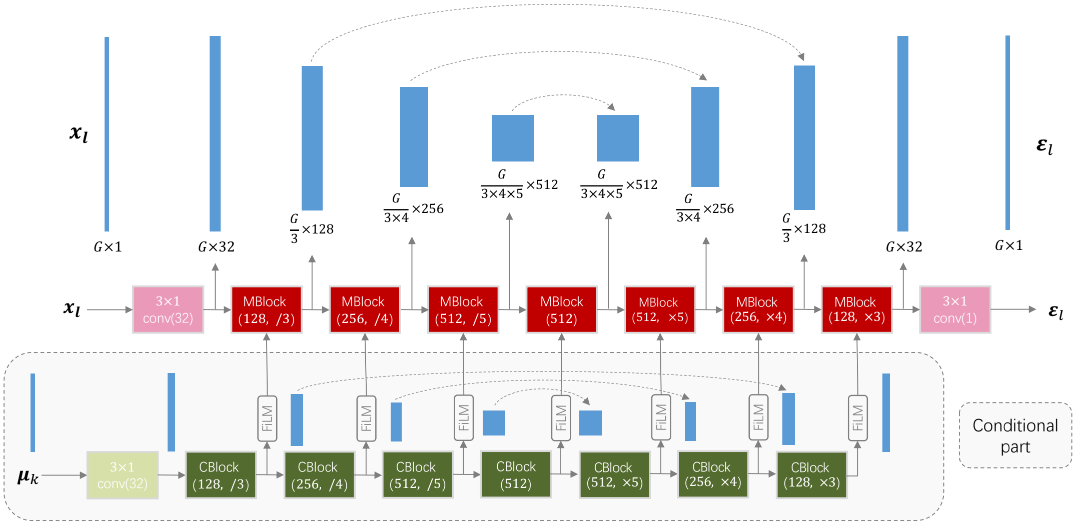

**Figure S22:** Architecture of the conditional *score network* $\mathbf{s}_{\boldsymbol{\theta}}(\mathbf{x}, C)$. The red color line is the UNet that takes $\mathbf{x}$ as input, and the green one is the conditional UNet that takes $\boldsymbol{\mu}_k$ as input, where $k$ in $\boldsymbol{\mu}_k$ is the cell type that $\mathbf{x}$ belongs to. We condition cell type information into score function by FiLM module, which produces both scale and bias vectors for feature-wise affine transformation as shown in S26.

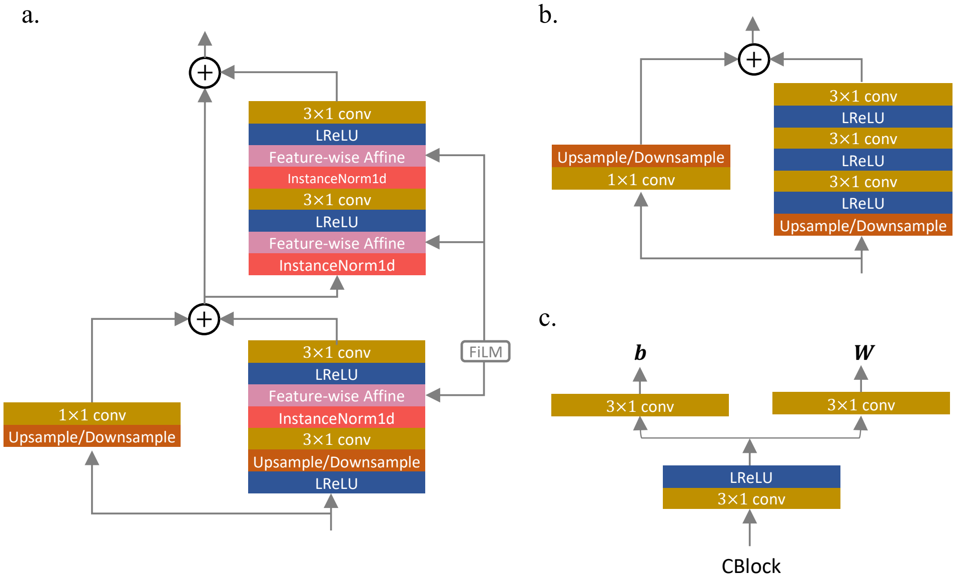

**Figure S23: The diagrams of MBlock, CBlock and FiLM**. **a**, The diagram of MBlock. Two residual blocks are used. The output of the FiLM block will be the input to the Feature-wise Affine block, as shown in (S26). **b**, The diagram of CBlock. One residual block is used, and this block's output will be the FiLM block's input (Fig. S23c). **c**, The diagram of FiLM. The block will take the output of CBlock as input and output scale $\mathbf{W}$ and shift $\mathbf{b}$.

#### 2.5 Hyper-parameters

Here we give the hyperparameters for training the score function and the decomposition process. As suggested by [11], we determine the values of $L, T, \{\sigma_l\}_{l=1}^{L}$, and $\eta_0$ as follows. In our default setting, we selected around 2,000 marker genes. We set $L = 232, T = 5$, and chose $\{\sigma_l\}_{l=1}^{L}$ to be a geometric progression:

$$c = \frac{\sigma_L}{\sigma_{L-1}} = \cdots = \frac{\sigma_2}{\sigma_1} > 1, \tag{S27}$$

where $c$ is a constant. Further, $\sigma_L$ is chosen to be as large as the maximum Euclidean distance between all pairs of different cells from single-cell reference data (See Table S1). We provide detailed hyperparameters setting in Table S1. We use the Adam optimizer [16] for all models with a learning rate 0.0001. Exponential moving average(EMA) is used in training, and the averaged parameters are used when sampling.

We find that the relative order of magnitude between the posterior $\nabla_{\mathbf{X}_i} \log p\left(\mathbf{y}_i|\mathbf{X}_i^{(t)}\right) + \nabla_{\mathbf{X}_i} \log p\left(\mathbf{X}_i^{(t)}|\mathbf{k}_i\right)$ and the injected noise $\sqrt{2\eta}\boldsymbol{\varepsilon}^{(t)}$ will affect the results of decomposition. Therefore, we use a little bit larger $\eta_0$ in decomposition (See Table S1) and let $\sigma_{yl} = \sigma_l^{0.5}$ in main text method Algorithm 1.

| Dataset | training dimension | $\sigma_1$ | $\sigma_L$ | $\eta_0$ | $\eta_0$ (decomposition) | downsampling channel dimension |
|---|---|---|---|---|---|---|
| Heart | 2195 | 0.002 | 50 | 3e-7 | 1e-6 | 128, 256, 512 |
| MOp | 1938 | 0.01 | 100 | 6.6e-6 | 1e-5 | 128, 256, 512 |
| VISp | 2027 | 0.01 | 100 | 6.6e-6 | 1e-5 | 128, 256, 512 |
| Cerebellum | 2394 | 0.01 | 100 | 6.6e-6 | 1e-5 | 128, 256, 512 |
| Hippocampus | 1923 | 0.002 | 50 | 3e-7 | 1e-6 | 128, 256, 512 |
| MERIFISH | 254 | 0.002 | 50 | 3e-7 | 1e-6 | 64, 128, 256 |

**Table S1:** Hyperparameters of SpatialScope for different datasets.

#### 2.6 Correction of the batch effects between single-cell reference and ST data

Recall that the batch effects between ST and single-cell reference data will hinder gene expression decomposition. To correct the batch effects, we adjust the gene-specific cross-platform effects using

$$\mathbf{y}_i = [y_{i,1}/\exp(\hat{\gamma}_1), \ldots, y_{i,G}/\exp(\hat{\gamma}_G)], \tag{S28}$$

where $y_{i,g}$ are the observed expression counts of gene $g$ at spot $i$ and $\hat{\gamma}_g$ is the batch effect of gene $g$ estimated under model (S6). Next, we account for the difference in sequencing depth by normalizing the total count of $\mathbf{y}_i$ to the mean of the total transcript counts of individual cells from single-cell reference data:

$$\mathbf{y}_i \leftarrow \frac{\mathbf{y}_i}{\sum_g y_{i,g}} \cdot \left(\frac{1}{N_{sc}} \sum_{n=1}^{N_{sc}} \sum_g x_{n,g}\right), \tag{S29}$$

where $N_{sc}$ is the total cell number of single-cell reference data and $x_{n,g}$ is the count data of cell $n$ and gene $g$ from single-cell reference data.

#### 2.7 Simulation design

##### 2.7.1 Main simulation analysis

*MERFISH MOp dataset* The MERFISH MOp dataset was profiled by the image-based ST approach with single-cell resolution. This dataset contains 254 genes and about 300,000 single cells located in 64 mouse brain MOp slices from 12 different samples [17]. As a concrete example demonstrated in the MERFISH paper, we first used the slice "mouse1_slice180" from "mouse1_sample4" to construct a simulation dataset shown in Fig. 2 of the main text. This slice consists of 5,551 cells and shows a multi-layer structure horizontally. Therefore, we vertically partitioned this data into two parts: the right part contains ∼4K cells and serves as paired single-cell reference data; the left part contains ∼1K cells and is used to generate low-resolution ST data by aggregating the cells on uniform grids to make simulated spots. The spot-level cell type compositions, cell type labels, and gene expression profiles of each cell within the simulated spots were used as the ground truth. To comprehensively evaluate the performance of SpatialScope and the compared methods, we varied the grid size from 16 × 16 µm to 42 × 42 µm, resulting in 1-3 and 1-6 cells within the simulated spots, respectively. We also varied the mean subsampled UMIs within the simulated spots ranging from 130 (averaged half-cell UMIs in MERFISH data) to 520 (averaged two-cell UMIs in MERFISH data). Besides, except for the paired single-cell reference data from the same slice, we additionally used an unpaired scRNA-seq data reference collected from the mouse Primary visual area [18] to show the robustness of SpatialScope.

Furthermore, we used another slice, "mouse1_slice122", to construct a new simulation dataset (Fig. S3c) with the same pipeline to demonstrate the generality of SpatialScope. The grid size in Fig. S3c is 62 × 61 µm, leading to 1-14 cells within the simulated spots. The subsampled UMIs is 600, approximately 2 cell UMIs in the slice "mouse1_slice122".

*STARmap cortex dataset* The STARmap dataset was also profiled by the image-based ST approach with single-cell resolution. This dataset was collected from the mouse visual cortex and comprised 1,020 genes measured in 973 cells [19]. We used the same simulation pipeline to generate simulated spots with grid size equal to 64 × 70 µm, leading to 1-14 cells within the simulated spots (Fig. S3b). The subsampled UMIs were set to 932, approximately 2 cell UMIs in raw STARmap data. We used paired scRNA-seq data collected from the mouse's Primary visual area as reference [18].

*MERFISH hypothalamus dataset* The MERFISH hypothalamus dataset was obtained from the mouse preoptic region of the hypothalamus from Dryad [20]. This dataset imaged about 1.1 million single cells from multiple slices. We used the tissue slice at Bregma-11 mm from animal ID 25 as an example, which contains expression values of 161 genes on 2447 single cells after preprocessing. Similarly, we applied the same simulation pipeline to generate simulated spots with grid size equal 24 × 30 µm, leading to 1-6 cells within the simulated spots (Fig. S3d). We set subsampled UMIs as 519, approximately 2 cell UMIs in raw MERFISH hypothalamus data.

We used paired scRNA-seq data collected from the preoptic region of the hypothalamus from multiple male and female mice [20].

##### 2.7.2 Missing cell types in single-cell reference

We performed additional simulations to evaluate the impact of missing cell types in reference. Specifically, following the MERFISH simulation analysis in the main text, we removed the L6b and Lamp5 cells from the single-cell reference, then we evaluated the effect of missing cell type on the cell type identification for SpatialScope and the compared methods. Overall, SpatialScope achieved the highest robustness over the compared methods by predicting the cells as the most transcriptionally similar cell type in the reference when the ground truth cell types were missing (Fig. S24). For example, most L6b cells were predicted to be L6 CT cells by SpatialScope as L6 CT is the closest cell type for L6b, and most Lamp5 cells were still classified as GABAergic neurons: Pvalb or Vip. In contrast, substantial L6b and Lamp5 cells were classified as L6 IT Car3 and L4/5 IT by Tangram, respectively. CytoSPACE unreasonably misclassified most Lamp5 cells as Endo, which is a non-neuronal cell type.

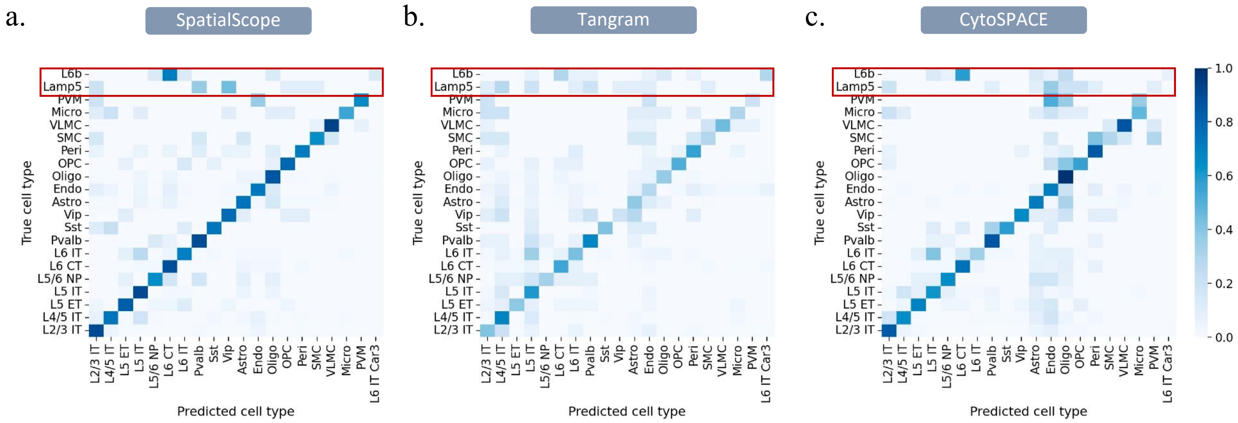

**Figure S24: Performance of SpatialScope and the compared method on cell types that do not appear in the single-cell reference**. **a**, Confusion matrix for SpatialScope's performance on missing cell types simulation analysis. Color represents the proportion of the cell type on the y-axis, classified as the cell type on the x-axis. **b**, Confusion matrix of Tangram. **c**, Confusion matrix of CytoSPACE.

##### 2.7.3 Inconsistent cell number in gene expression decomposition task

Although deep nucleus segmentation tools have shown great success in identifying cells in microscopy images, the segmentation result is not 100 percent accurate. For example, some cells were missing due to the weak signal, and some noise pixels may be misidentified as cells. Therefore, it is necessary to evaluate the performance of SpatialScope and the compared method when the ground truth cell number is inconsistent with the estimated cell number in the spots. Following the MERFISH simulation analysis in the main text, we created additional simulations in the presence of inconsistent cell numbers. Specifically, based on the original ST data (the second part of the partitioned MERFISH simulation dataset, Fig. S25a), we randomly selected some cells as missing cells whose gene expressions still contribute to the simulated spot-level expression profile, but they are not observed in the new ST data. The remaining cells existing in the original ST data (denoted as existing cells) are still observed and contribute to the spot-level gene expressions. Besides, we further randomly added some cells as mis-added cells in this new ST data to mimic the misidentified cells due to the noise pixels. Those mis-added cells have no gene expressions and thus cannot contribute to the spot-level expression profiles, but they are observed in the new ST data (Fig. S25b). Next, following the same analysis pipeline as in the main text, we aggregated all observed cells on uniform grids to generate simulated spots with the grid size: 34×30(µm) and varied subsampled UMIs ranging from 130 to 520. Then we used the paired single-cell reference (the first part of the partitioned MERFISH simulation dataset) to perform cell type identification and gene expression decomposition. Of note, we only used the gene expression profiles for the existing cells within the simulated spots as ground truth because the missing cells are not observed, and the mis-added cells have no ground truth.

Fig. S25c shows the overall gene expression decomposition results for this new ST data when considering the spots with missing or mis-added cells. Clearly, SpatialScope achieved significantly higher accuracy of decomposition in all settings. The mean cosine similarity between the ground truth and predicted single-cell level gene expressions by SpatialScope is as high as 0.886 when the subsample UMIs count is 260 and the cell type labels are correctly identified, while the alignment-based methods, Tangram and CytoSPACE, only achieved 0.747 and 0.791 mean cosine similarities, respectively. As some concrete examples of spots with missing cells (Fig. S26), spot 108 contains three cells from L6 CT, Lamp5 and SMC, but the SMC cell was missing. SpatialScope successfully identified the remaining two ground truth cells with the highly matched transcriptional profiles, while the compared methods failed in both cell type identification and gene expression decomposition; Spot 54 contains three cells from L4/5 IT, Endo and L5 ET, but the L5 ET cell was missing. Although SpatialScope misidentified the L4/5 IT cell as L5 IT, the cosine similarity between the predicted and ground truth gene expression is still as high as 0.94 for L4/5 IT cell. This is because L4/5 IT and L5 IT are transcriptionally similar cell types in the reference and the L4/5 IT cell in this spot is very close to L5 IT cluster in the UMAP plot, which leads to the misidentified L5 IT cell by SpatialScope. Nevertheless, with the gene expression distribution approximated by the score-based generative model, SpatialScope can still find the best-matched transcriptional profile as much as possible even though the cell type label is misidentified. For spots with mis-added cells, we also provided four examples in Fig. S27. Spot 463 contains four existing cells from L4/5 IT and Endo, and one mis-added cell. SpatialScope successfully identified three of the four ground truth cells with a mean cosine similarity of 0.82, and the mis-added cell was identified as the major cell type in this spot: L4/5 IT. Spot 8 contains three existing cells from L4/5 IT and L2/3 IT, and two mis-added cells. SpatialScope successfully identified two of the three ground truth cells with a mean cosine similarity of 0.89, and the two mis-added cells were also identified as the major cell type in this spot.

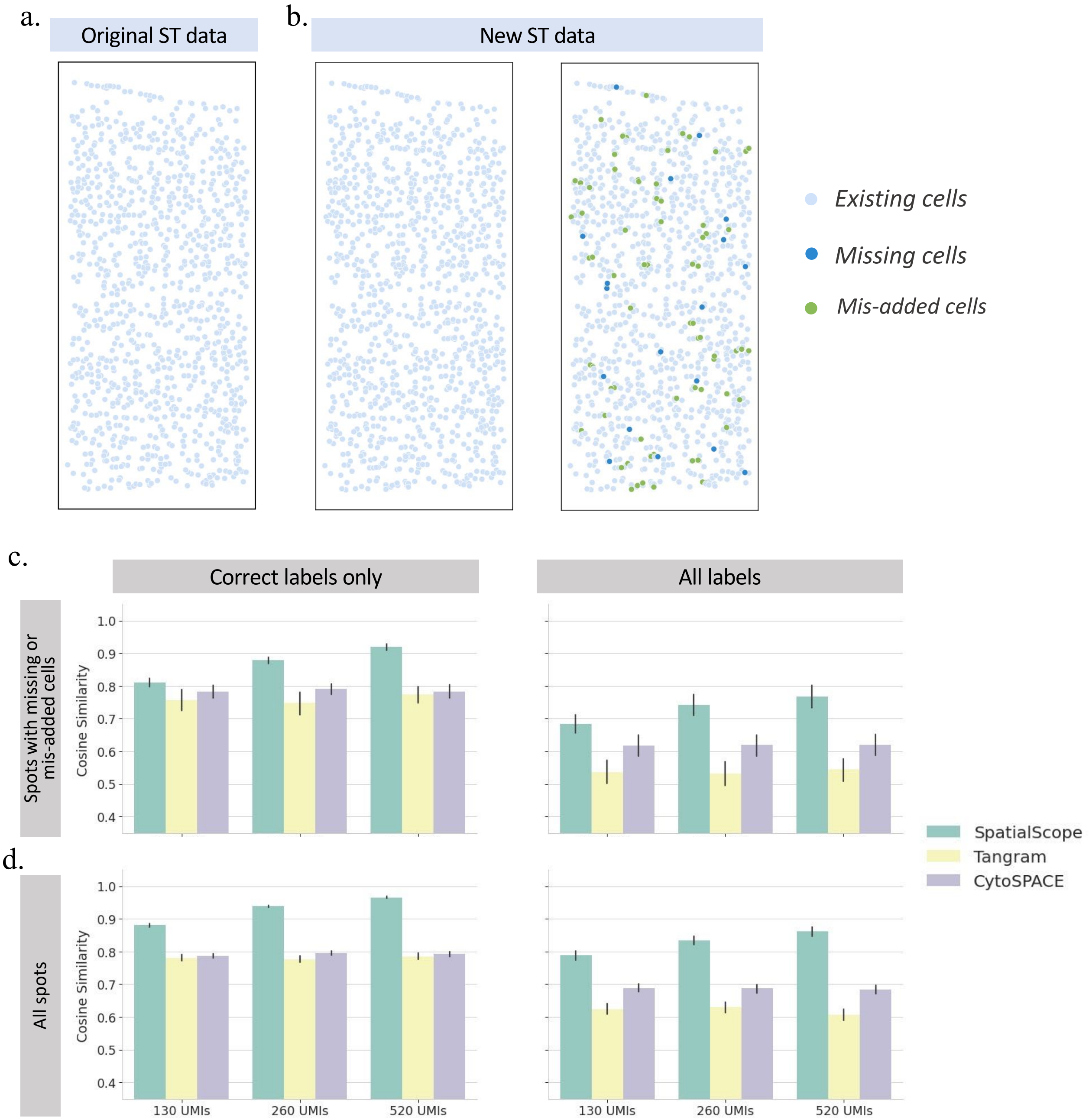

**Figure S25: Influence of inconsistent cell number on gene expression decomposition task**. **a**, Original ST cells in the MERFISH simulation dataset. **b**, The new ST data after randomly removing and adding some cells. Missing cells are colored blue, and mis-added cells are colored green. **c-d**, The cosine similarities between the ground truth and predicted gene expressions by the considered methods for cells in spots with inconsistent cell number (c) or all spots (d) under different scenarios of subsampled UMIs count. We further considered cells with correctly identified cell type labels (left) or all cells (right) as in the main text.

a.

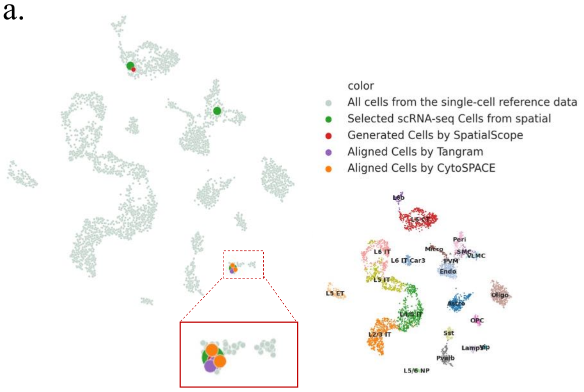

| | | cell1 | cell2 | cell3 |
|---|---|---|---|---|
| | True label | L6 CT | Lamp5 | SMC |
| SpatialScope | Estimated. | L6 CT | Lamp5 | ------ |
|  | Cos. sim. | 0.94 | 0.96 | ------ |
| Tangram | Estimated. | Lamp5 | Lamp5 | ------ |
|  | Cos. sim. | 0.38 | 0.80 | ------ |
| CytoSPACE | Estimated. | Lamp5 | Lamp5 | ------ |
|  | Cos. sim. | 0.41 | 0.78 | ------ |

b.

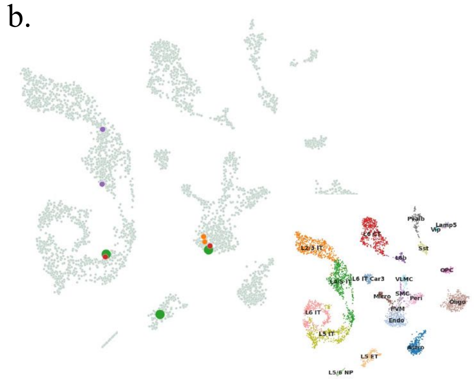

| | | cell1 | cell2 | cell3 |
|---|---|---|---|---|
| | True label | L4/5 IT | Endo | L5 ET |
| SpatialScope | Estimated. | L5 IT | Endo | ------ |
|  | Cos. sim. | 0.94 | 0.88 | ------ |
| Tangram | Estimated. | L4/5 IT | L4/5 IT | ------ |
|  | Cos. sim. | 0.60 | 0.16 | ------ |
| CytoSPACE | Estimated. | Endo | Endo | ------ |
|  | Cos. sim. | 0.36 | 0.70 | ------ |

c.

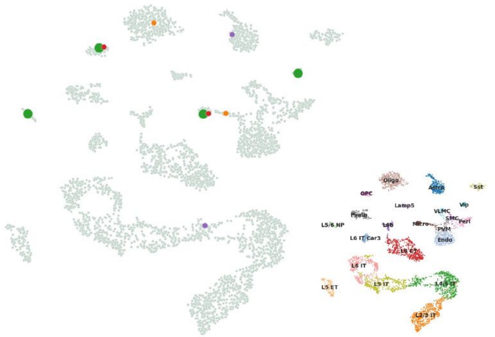

| | | cell1 | cell2 | cell3 | cell4 |
|---|---|---|---|---|---|
| | True label | Micro | OPC | Vip | L5/6 NP |
| SpatialScope | Estimated. | Micro | OPC | ------ | ------ |
|  | Cos. sim. | 0.93 | 0.98 | ------ | ------ |
| Tangram | Estimated. | L4/5 IT | Astro | ------ | ------ |
|  | Cos. sim. | 0.19 | 0.22 | ------ | ------ |
| CytoSPACE | Estimated. | Micro | Oligo | ------ | ------ |
|  | Cos. sim. | 0.61 | 0.37 | ------ | ------ |

d.

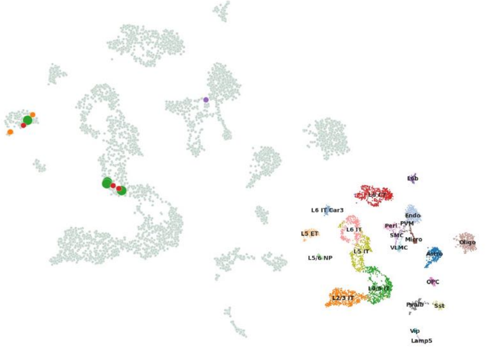

| | | cell1 | cell2 | cell3 | cell4 |
|---|---|---|---|---|---|
| | True label | L5 IT | L5 ET | L4/5 IT | L5 IT |
| SpatialScope | Estimated. | L5 IT | L5 ET | L5 IT | ------ |
|  | Cos. sim. | 0.86 | 0.89 | 0.90 | ------ |
| Tangram | Estimated. | ------ | ------ | L5 IT | ------ |
|  | Cos. sim. | ------ | ------ | 0.88 | ------ |
| CytoSPACE | Estimated. | L5 ET | L5 ET | L5 ET | ------ |
|  | Cos. sim. | 0.64 | 0.80 | 0.58 | ------ |

**Figure S26: Examples of spots with missing cells in the gene expression decomposition task**. **a**, The UMAP plot of gene expression decomposition example for spot 108, which contains three cells from L6 CT, Lamp5 and SMC, but the SMC cell was missing. The cell type identification and gene expression decomposition results were shown in the table below. Estimate.: Estimated cell type label by SpatialScope and the compared method. Cos. sim.: Cosine similarity between the ground truth and predicted gene expressions. **b**, The UMAP plot of gene expression decomposition example for spot 54, which contains three cells from L4/5 IT, Endo and L5 ET, but the L5 ET cell was missing. **c**, The UMAP plot of gene expression decomposition example for spot 140, which contains four cells from Micro, OPC, Vip and L5/6 NP, but the Vip and L5/6 NP cells were missing. **d**, The UMAP plot of gene expression decomposition example for spot 593, which contains four cells from L5 IT, L4/5 IT and L5 ET, but one of the L5 IT cells was missing.

a.

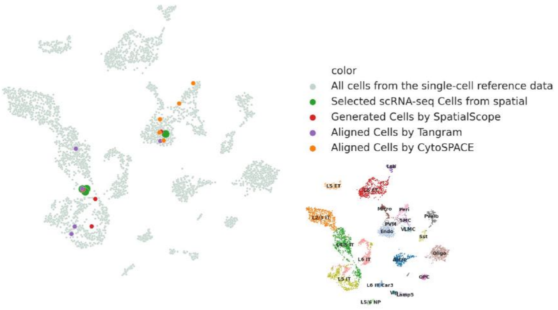

| | | cell1 | cell2 | cell3 | cell4 | cell5 |
|---|---|---|---|---|---|---|
|  | True label | L4/5 IT | L4/5 IT | L4/5 IT | Endo | ------ |
| SpatialScope | Estimated. | L4/5 IT | L5 IT | L4/5 IT | Endo | L4/5 IT |
|  | Cos. sim. | 0.83 | 0.68 | 0.79 | 0.82 | ------ |
| Tangram | Estimated. | L4/5 IT | L4/5 IT | L5 IT | L5 IT | Endo |
|  | Cos. sim. | 0.81 | 0.60 | 0.68 | 0.18 | ------ |
| CytoSPACE | Estimated. | Endo | Endo | Peri | Peri | PVM |
|  | Cos. sim. | 0.43 | 0.29 | 0.39 | 0.71 | ------ |

b.

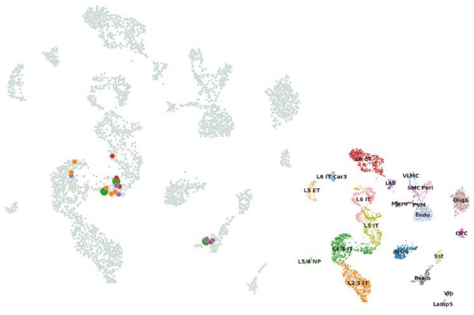

| | | cell1 | cell2 | cell3 | cell4 | cell5 |
|---|---|---|---|---|---|---|
|  | True label | L4/5 IT | Pvalb | L4/5 IT | ------ | ------ |
| SpatialScope | Estimated. | L4/5 IT | Pvalb | L5 IT | L5 IT | Pvalb |
|  | Cos. sim. | 0.76 | 0.90 | 0.73 | ------ | ------ |
| Tangram | Estimated. | L4/5 IT | L4/5 IT | L4/5 IT | Pvalb | Pvalb |
|  | Cos. sim. | 0.71 | 0.38 | 0.42 | ------ | ------ |
| CytoSPACE | Estimated. | L4/5 IT | L4/5 IT | L4/5 IT | L4/5 IT | L4/5 IT |
|  | Cos. sim. | 0.79 | 0.53 | 0.64 | ------ | ------ |

c.

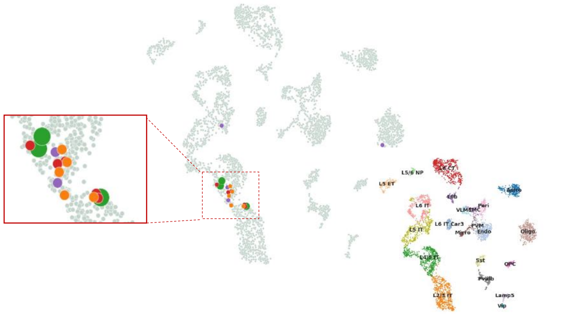

| | | cell1 | cell2 | cell3 | cell4 | cell5 |
|---|---|---|---|---|---|---|
|  | True label | L4/5 IT | L4/5 IT | L2/3 IT | ------ | ------ |
| SpatialScope | Estimated. | L2/3 IT | L4/5 IT | L2/3 IT | L4/5 IT | L4/5 IT |
|  | Cos. sim. | 0.79 | 0.87 | 0.90 | ------ | ------ |
| Tangram | Estimated. | L4/5 IT | L4/5 IT | L5 IT | Oligo | ------ |
|  | Cos. sim. | 0.71 | 0.78 | 0.72 | ------ | ------ |
| CytoSPACE | Estimated. | L2/3 IT | L4/5 IT | L4/5 IT | L4/5 IT | L4/5 IT |
|  | Cos. sim. | 0.80 | 0.82 | 0.89 | ------ | ------ |

d.

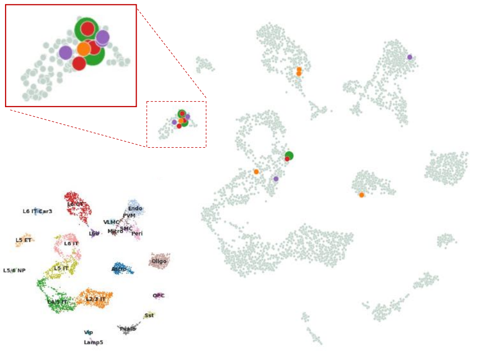

| | | cell1 | cell2 | cell3 | cell4 | cell5 |
|---|---|---|---|---|---|---|
|  | True label | L5 ET | L6 IT | L5 ET | ------ | ------ |
| SpatialScope | Estimated. | L5 ET | L5 ET | L6 IT | L5 ET | L5 ET |
|  | Cos. sim. | 0.95 | 0.63 | 0.61 | ------ | ------ |
| Tangram | Estimated. | L5 IT | Endo | L5 ET | L5 ET | L5 ET |
|  | Cos. sim. | 0.60 | 0.20 | 0.76 | ------ | ------ |
| CytoSPACE | Estimated. | Astro | L5 ET | L5 IT | L6 CT | L6 CT |
|  | Cos. sim. | 0.34 | 0.60 | 0.60 | ------ | ------ |

**Figure S27: Examples of spots with mis-added cells in the gene expression decomposition task**. **a**, The UMAP plot of gene expression decomposition example for spot 463, which contains four existing cells from L4/5 IT and Endo, and one mis-added cell. The cell type identification and gene expression decomposition results were shown in the table below. Estimate.: Estimated cell type label by SpatialScope and the compared method. Cos. sim.: Cosine similarity between the ground truth and predicted gene expressions. **b**, The UMAP plot of gene expression decomposition example for spot 13, which contains three existing cells from L4/5 IT and Pvalb, and two mis-added cells. **c**, The UMAP plot of gene expression decomposition example for spot 8, which contains three existing cells from L4/5 IT and L2/3 IT, and two mis-added cells. **d**, The UMAP plot of gene expression decomposition example for spot 437, which contains three existing cells from L5 ET and L6 IT, and two mis-added cells.

##### 2.7.4 Different training epochs

Next, we investigate the performance of SpatialScope under score-based generative models with different training epochs. Precisely, we followed the simulation analysis in the main text to evaluate the gene expression decomposition performance but varied the training epochs of the score-based generative model saved at 500, 1500, 7500, 12500, and 25000 epochs. Among them, 7500 epoch was used in the simulation analysis of the main text. To fairly compare the performance of models with different training epochs, we only considered cells with the correct estimated cell type label. In the simple case when the simulated spots only contain one cell (Fig. S28a), increasing the number of epochs improves the performance significantly when the subsample UMIs count is low. This result suggests that the score-based generative model can better approximate the gene expression distribution with the increase in the number of epochs. As expected, when considering the spots with cell numbers larger than one (Fig. S28b), the gene expression decomposition performance improved steadily as the number of epochs increased for all settings. These results indicate that the score-based generative model can benefit from increasing the training epochs in most cases. However, due to the trade-off between the performance and the time cost, we recommend that the number of epochs ranges from 5,000 to 10,000, as the improvement after the 10,000 epoch is minor.

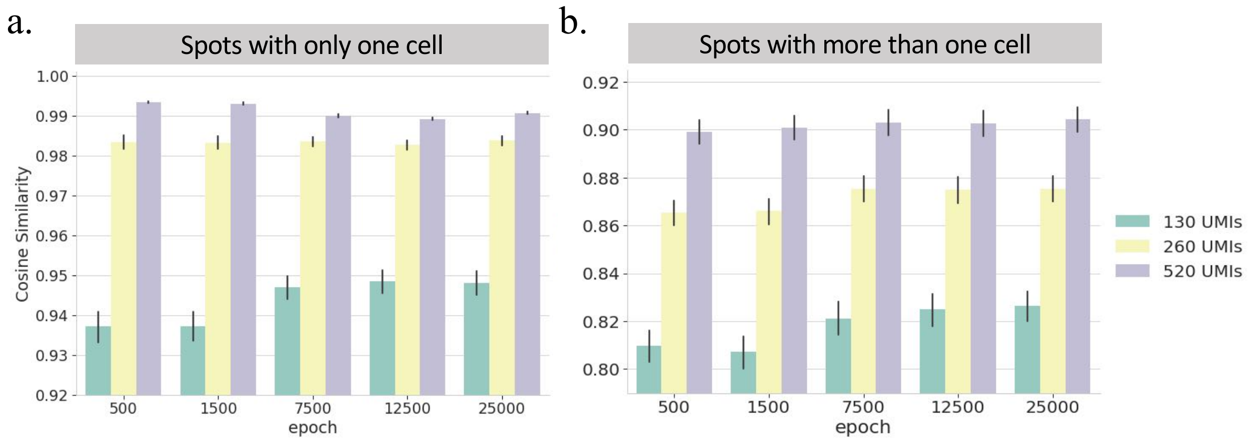

**Figure S28: The gene expression decomposition performance for spots with only one cell (a) or cell number larger than one (b) when different checkpoints of the score-based generative model were used in the MERFISH simulation dataset.**

##### 2.7.5 The comparison of conditional and unconditional score function

As illustrated in Section 2.3 (Gene expression decomposition), we learned conditional score-based generative models from single-cell reference data by additionally embedding the cell type mean gene expression as inputs, which increases the flexibility of our model to accommodate the heterogeneity across cell types. More specifically, the unconditional network requires learning the gene expression distribution of all cell types with one shared network. By contrast, embedding the cell-type information in the conditional network allows capturing cell-type specific information and retaining modeling flexibility. To see this, we trained an unconditional score-based generative model using the paired single-cell reference from the simulation dataset and compared the results with those of the corresponding conditional network. Unsurprisingly, the cells sampled from the distribution learned by the conditional network (Fig. S29a) gradually overlapped with the original single-cell reference data with training epoch increases, while cells generated by the unconditional network continuously showed strong evidence of deviation from the reference, especially for L4/5 IT and L5 IT cell types (Fig. S29b). We further applied the kBET to quantitatively assess how well the single-cell reference and generated cells mixed. kBET hypothesized that the proportions of the batch labels in any neighborhood do not differ from the global distribution in the absence of batch-effect, and it used the rejection rates to quantify the degree of mixing: low rejection rates imply well-mixed datasets, a low rejection rate of 0.2 roughly corresponds to 1% of biased genes (mean gene expression was varied) between the two datasets while a high rejection rate of 0.8 corresponds to 20% of biased genes. [21]. We observed that the kBET rejection rates decreased from 0.771 to 0.296 as the number of epochs increased from 500 to 7500 for the conditional network, while kBET rejection rates of the unconditional network only reduced a little (from 0.809 to 0.759).

Next, we followed the simulation analysis in the main text and evaluated the gene expression decomposition performance between conditional and unconditional networks. We used the checkpoints saved in 7500 epochs for both networks. As shown in Fig. S29c, in the naïve case when the simulated spots only contain one cell, the conditional network is slightly better than the unconditional network when the subsampling UMIs count is as low as 130, and two kinds of networks are comparable when subsampling UMIs larger than 130. However, when considering the spots with cell number larger than one (Fig. S29d), the conditional network shows better performance over the unconditional network in all settings, indicating that the conditional network can improve the gene expression decomposition by learning the gene expression distribution of different cell types in some sense specifically.

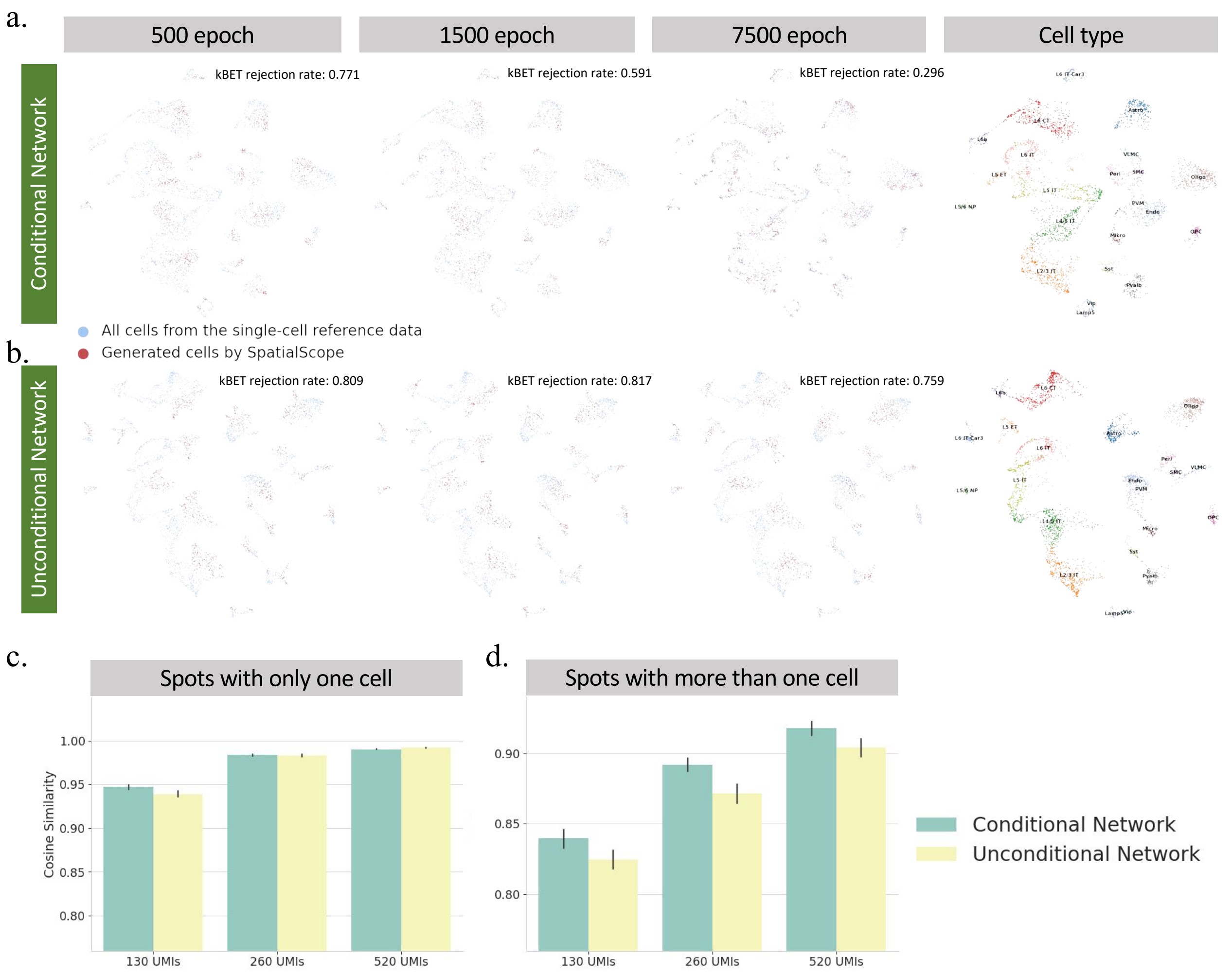

**Figure S29: Comparison of conditional and unconditional networks in training the score-based generative model**. The learning process of the score-based generative model for the conditional network (**a**) and unconditional network (**b**). UMAP of single cell reference data and the pseudo cells generated by the deep generative model at different epochs. The blue dots represent the existing cells from scRNA-seq data and the red dots represents cells generated by SpatialScope. **c**, Gene expression decomposition performance between conditional and unconditional networks for spots with only one cell. **d**, Gene expression decomposition performance for spots with cell number larger than one.

##### 2.7.6 Unbalanced cell types in single-cell reference data

Due to the large variation in the proportions of different cell types, we created additional simulations to compare the performance of gene expression decomposition in different cell types using the MERFISH simulation dataset. Specifically, for each cell type, we simulated 100 spots by randomly selecting two cells of this cell type from the second part (∼1K cells) of the partitioned MERFISH simulation dataset. Then we evaluated the gene expression decomposition performance using the simulated spots as ST data and the first part of the partitioned MERFISH simulation dataset as single-cell reference data. We observed that there is no association between decomposition performance and cell type proportions (Fig. S30a). For example, L4/5 IT (12.6%) and Vip (0.5%) are the most abundant and rare cell types in the single cell reference, respectively, but they showed similar gene expression decomposition performance, suggesting the score function is not influenced by the unbalanced cell types. In addition, we also showed that the heterogeneity within the cell types mainly drove the difference in decomposition performance (Fig. S30b). For example, decomposition within the cell type Micro performs worst as Micro has the maximum heterogeneity with an average mean cosine similarity of 0.51 (See Fig. S30b).

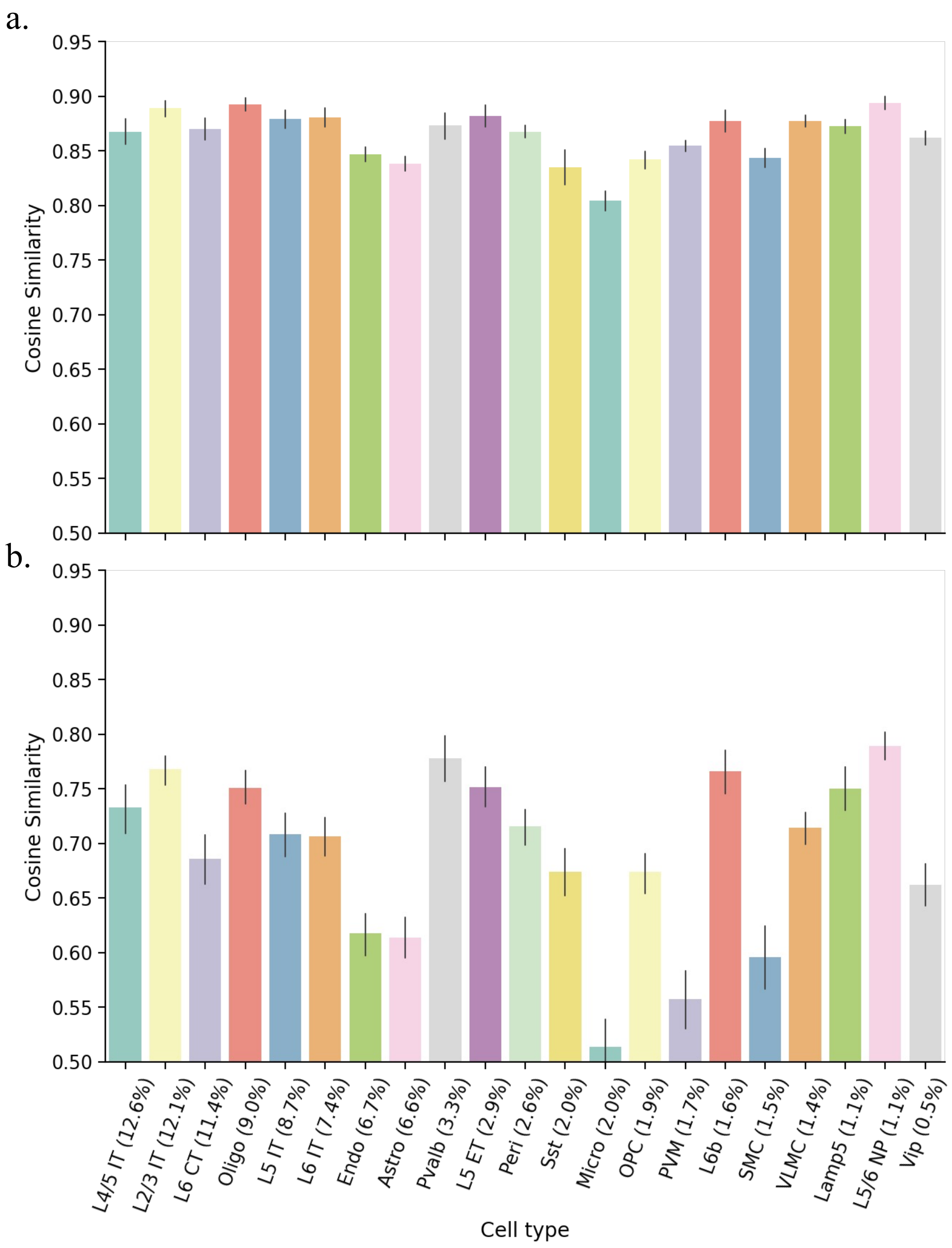

**Figure S30: Influence of unbalanced cell types in training the score-based generative model**. **a**, Gene expression decomposition performance of SpatialScope for each cell type in the MERFISH simulation dataset, the cell types are sorted by their proportion (shown in the x-ticks) in the single-cell reference data. **b**, The mean cosine similarities between two randomly selected cells for each cell type in the paired single-cell reference of the MERFISH simulation dataset.

### References

[1] Dylan M Cable, Evan Murray, Luli S Zou, Aleksandrina Goeva, Evan Z Macosko, Fei Chen, and Rafael A Irizarry. Robust decomposition of cell type mixtures in spatial transcriptomics. *Nature Biotechnology*, 40(4):517–526, 2022.

[2] Pascal Vincent. A connection between score matching and denoising autoencoders. *Neural computation*, 23(7):1661–1674, 2011.

[3] Max Welling and Yee W Teh. Bayesian learning via stochastic gradient langevin dynamics. In *Proceedings of the 28th international conference on machine learning (ICML-11)*, pages 681–688. Citeseer, 2011.

[4] Mehdi Mirza and Simon Osindero. Conditional generative adversarial nets. *arXiv preprint arXiv:1411.1784*, 2014.

[5] Kihyuk Sohn, Honglak Lee, and Xinchen Yan. Learning structured output representation using deep conditional generative models. *Advances in neural information processing systems*, 28, 2015.

[6] Andrew Brock, Jeff Donahue, and Karen Simonyan. Large scale gan training for high fidelity natural image synthesis. *arXiv preprint arXiv:1809.11096*, 2018.

[7] Takeru Miyato and Masanori Koyama. cgans with projection discriminator. *arXiv preprint arXiv:1802.05637*, 2018.

[8] Augustus Odena, Christopher Olah, and Jonathon Shlens. Conditional image synthesis with auxiliary classifier gans. In *International conference on machine learning*, pages 2642–2651. PMLR, 2017.

[9] Matthew Tancik, Pratul Srinivasan, Ben Mildenhall, Sara Fridovich-Keil, Nithin Raghavan, Utkarsh Singhal, Ravi Ramamoorthi, Jonathan Barron, and Ren Ng. Fourier features let networks learn high frequency functions in low dimensional domains. *Advances in Neural Information Processing Systems*, 33:7537–7547, 2020.

[10] Yang Song and Stefano Ermon. Generative modeling by estimating gradients of the data distribution. *Advances in Neural Information Processing Systems*, 32, 2019.

[11] Yang Song and Stefano Ermon. Improved techniques for training score-based generative models. *Advances in neural information processing systems*, 33:12438–12448, 2020.

[12] Olaf Ronneberger, Philipp Fischer, and Thomas Brox. U-net: Convolutional networks for biomedical image segmentation. In *International Conference on Medical image computing and computer-assisted intervention*, pages 234–241. Springer, 2015.

[13] Jonathan Ho, Ajay Jain, and Pieter Abbeel. Denoising diffusion probabilistic models. *Advances in Neural Information Processing Systems*, 33:6840–6851, 2020.

[14] Prafulla Dhariwal and Alexander Nichol. Diffusion models beat gans on image synthesis. *Advances in Neural Information Processing Systems*, 34:8780–8794, 2021.

[15] Nanxin Chen, Yu Zhang, Heiga Zen, Ron J Weiss, Mohammad Norouzi, and William Chan. Wavegrad: Estimating gradients for waveform generation. *arXiv preprint arXiv:2009.00713*, 2020.

[16] Diederik P Kingma and Jimmy Ba. Adam: A method for stochastic optimization. *arXiv preprint arXiv:1412.6980*, 2014.

[17] Meng Zhang, Stephen W Eichhorn, Brian Zingg, Zizhen Yao, Kaelan Cotter, Hongkui Zeng, Hongwei Dong, and Xiaowei Zhuang. Spatially resolved cell atlas of the mouse primary motor cortex by merfish. *Nature*, 598(7879):137–143, 2021.

[18] Bosiljka Tasic, Zizhen Yao, Lucas T Graybuck, Kimberly A Smith, Thuc Nghi Nguyen, Darren Bertagnolli, Jeff Goldy, Emma Garren, Michael N Economo, Sarada Viswanathan, et al. Shared and distinct transcriptomic cell types across neocortical areas. *Nature*, 563(7729):72–78, 2018.

[19] Xiao Wang, William E Allen, Matthew A Wright, Emily L Sylwestrak, Nikolay Samusik, Sam Vesuna, Kathryn Evans, Cindy Liu, Charu Ramakrishnan, Jia Liu, et al. Three-dimensional intact-tissue sequencing of single-cell transcriptional states. *Science*, 361(6400):eaat5691, 2018.

[20] Jeffrey R Moffitt, Dhananjay Bambah-Mukku, Stephen W Eichhorn, Eric Vaughn, Karthik Shekhar, Julio D Perez, Nimrod D Rubinstein, Junjie Hao, Aviv Regev, Catherine Dulac, et al. Molecular, spatial, and functional single-cell profiling of the hypothalamic preoptic region. *Science*, 362(6416):eaau5324, 2018.

[21] Maren Büttner, Zhichao Miao, F Alexander Wolf, Sarah A Teichmann, and Fabian J Theis. A test metric for assessing single-cell rna-seq batch correction. *Nature methods*, 16(1):43–49, 2019.
